# A yeast with muscle doesn’t run faster: full humanization of the glycolytic pathway in *Saccharomyces cerevisiae*

**DOI:** 10.1101/2021.09.28.462164

**Authors:** Francine J. Boonekamp, Ewout Knibbe, Marcel A. Vieira-Lara, Melanie Wijsman, Marijke A.H. Luttik, Karen van Eunen, Maxime den Ridder, Reinier Bron, Ana Maria Almonacid Suarez, Patrick van Rijn, Justina C. Wolters, Martin Pabst, Jean-Marc Daran, Barbara Bakker, Pascale Daran-Lapujade

**Author notes:** These authors have contributed equally to this work (Francine J. Boonekamp, Ewout Knibbe).

## Abstract

While transplantation of single genes in yeast plays a key role in elucidating gene functionality in metazoans, technical challenges hamper the humanization of full pathways and processes. Empowered by advances in synthetic biology, this study demonstrates the feasibility and implementation of full humanization of glycolysis in yeast. Single gene and full pathway transplantation revealed the remarkable conservation of both glycolytic and moonlighting functions and, combined with evolutionary strategies, brought to light novel, context-dependent responses. Remarkably, human hexokinase 1 and 2, but not 4, required mutations in their catalytic or allosteric sites for functionality in yeast, while hexokinase 3 was unable to complement its yeast ortholog. Comparison with human tissues cultures showed the preservation of turnover numbers of human glycolytic enzymes in yeast and human cell cultures. This demonstration of transplantation of an entire, essential pathway paves the way to the establishment of species, tissue and disease-specific metazoan models.

**One Sentence Summary:** This work demonstrates the successful humanization of an entire pathway in *Saccharomyces cerevisiae* and establishes an attractive strategy to study (human) glycolysis architecture and regulation.

**Highlights:**

- The successful humanization of the entire glycolytic pathway in yeast offers new microbial models for both fundamental and applied studies.
- Both glycolytic and moonlighting functions and turnover numbers of glycolytic enzymes are highly conserved between yeast and human.
- Functionality of human hexokinases 1 and 2 in yeast requires mutations in the catalytic or allosteric binding sites.
- Combination of single gene and full transplantation with laboratory evolution reveals context-dependent activity and evolution of glycolytic enzymes.

## Introduction

Due to its tractability and genetic accessibility, *S. cerevisiae* has played and still plays a key role as simplified model organism for higher eukaryotes. Many discoveries in yeast native processes such as the cell cycle and ribosome biogenesis were pivotal for understanding their mammalian equivalents [1, 2]. In addition, yeast is used to study a wide range of diseases such as cancer, diabetes and neurodegenerative diseases [3]. In a large part of these studies, the heterologous expression of human genes in yeast enables the detailed investigation of human biology and disease-specific variations of human genes [4]. As the yeast and human genome share over 2000 groups of orthologs [4], several large initiatives have explored the complementarity of human genes in yeast and shown a high degree of functional conservation [4–11]. These studies are however complicated by the genetic redundancy of eukaryotic genomes [12], which is even more prominent for genes encoding proteins with metabolic functions [13, 14].

While individual gene complementation in yeast is an interesting approach to characterize single human proteins, humanization of entire pathways or processes would greatly increase their usefulness. Such ‘next level’ yeast models hold the potential to capture the native functional context of the humanized proteins and to enable the study of more complex, multigene phenotypes, and epistatic interactions between genes. The feasibility of such extensive humanization projects depend largely on the replaceability of yeast genes by their human orthologs. Recent large scale humanization and bacterialization efforts of the yeast genome suggested that replaceability was better predicted on pathway- or process-basis than by sequence conservation [8, 15]. To date, reports of full humanization of pathways or protein complexes are scarce [5, 16–19]. However, rapid developments in synthetic biology have tremendously increased the ability to extensively remodel microbial genomes, and promise to bring more examples of large scale humanization in the future.

The Embden-Meyerhof-Parnas (EMP) pathway of glycolysis, which is near ubiquitous to eukaryotes, has a central role in carbon metabolism and is involved in a wide range of diseases in mammals, including cancer with the well-known Warburg effect [20]. So far, few single human glycolytic enzymes have been transplanted into yeast, mostly in large-scale complementation studies [6, 8, 9, 22–24]. Whether all human glycolytic enzymes can complement their yeast orthologs is however unknown. It is a particularly fascinating question as glycolytic enzymes, both in yeast and human are characterized by their versatility in moonlighting capabilities [25, 26]. The degree of conservation of these moonlighting functions between these two distant organisms has hardly been explored to date, with the exception of the human aldolase B (*Hs*ALDOB) and the glucokinase (*Hs*HK4) [23, 24].

To overcome the difficulty caused by genetic redundancy, a yeast strain in which the set of genes encoding glycolytic enzymes has been minimized from 19 to 11 was previously constructed [27]. This minimal glycolysis (MG) strain is a perfect platform for single glycolytic gene complementation. Furthermore, a strain in which this minimized set of yeast glycolytic genes has been fully relocated to a single chromosomal locus (SwYG strain) enables the swapping of the entire yeast glycolytic pathway by any designer glycolysis with minimal genetic engineering [28]. In the present study, glycolysis swapping with the SwYG strain was used to demonstrate the functionality of an entire human muscle glycolytic pathway in yeast. A combination of single gene complementation, full pathway humanization and adaptive laboratory evolution was used to explore the functionality of all human glycolytic genes in yeast. This led to the identification of mutations in human hexokinase 1 and 2 related to allosteric inhibition by glucose-6-phosphate, which appear to be required for functional expression in yeast. Finally the validity of yeast strains with humanized glycolysis as a model was evaluated by comparing the protein turnover number (k_cat_) of the human glycolytic enzymes expressed in yeast with enzymes in their native environment from human skeletal muscle myotube cell cultures.

## Results

### All human glycolytic genes directly complement their yeast ortholog except for hexokinases 1-3

With the exception of hexokinases and F1,6bP aldolases, human and yeast glycolytic enzymes are highly conserved with 43% to 65% identity at protein level, as compared to the 32% identity at whole proteome level [8] (Fig. 1). The human and yeast F1,6bP-aldolases belong to two different classes of enzymes and do not share homology at all at protein level [29] (Fig. 1 and Table S1). Among the four human hexokinases (*Hs*HK1 to *Hs*HK4), *Hs*HK4 is closest in size and sequence to *Sc*Hxk2 (ca. 30% protein identity) while *Hs*HK1, *Hs*HK2 and *Hs*HK3 are roughly twice the size of their yeast orthologs, with each subunit sharing ca. 30% identity with *Sc*Hxk2 ([30], Table 1). Due to the genetic redundancy of metabolic pathways in eukaryotes [13, 14], and associated difficulty of complementation studies, so far complementation in *S. cerevisiae* was only tested for eight human glycolytic genes [7-9, 22, 24], of which only *HsPGAM2* was unsuccessful (Table S1, [9, 31]). Implementation of the MG yeast strain, which carries a single isoenzyme for each glycolytic step, with the notable exception of the *Sc*Pfk1 and *Sc*Pfk2 subunits of the hetero-octameric phosphofructokinase [27], considerably facilitates complementation studies (Fig. 1). The ability of 25 human glycolytic genes to complement their yeast ortholog(s) was systematically explored by individual gene complementation in the MG strain. For enzymes with multiple splicing variants, the canonical version was used (Fig. 1 and Table S1). However, as the two pyruvate kinase genes *HsPKLR* and *HsPKM* have tissue-specific splicing variants (*HsPKL* and *HsPKR for HsPKLR*, and *HsPKM1* and *HsPKM2* for *HsPKM*), all four variants were tested (Fig. 1, Table S1 [32]). The 25 genes were codon-optimized, cloned downstream strong, constitutive promoters (Table S2) and individually cloned in the MG strain, after which the yeast ortholog(s) were removed (Fig. S1). Remarkably, 22 out of these 25 genes demonstrated direct complementation of their yeast orthologs for growth on glucose (Fig. 1, Fig. S2). Additionally, *HsHK1* and *HsHK2* but not *HsHK3* also complemented their yeast orthologs, but only after a period of adaptation of several days. While most strains were only marginally affected by single humanization of the glycolytic genes, strains harbouring a human hexokinase 2, the aldolases, phosphoglycerate mutases and the glyceraldehyde-3P dehydrogenase GAPDH variant S had a strongly reduced growth rate, the strongest decrease (30%) occurring with *Hs*ALDOB (Fig. 1). No clear correlation could be found between growth rate and conservation between human and yeast gene sequences or promoter strength (Fig S3). All human genes were Sanger-sequenced in the complementation strains, revealing that all besides *HsHK1* and *HsHK2* had the expected sequence (see following section). This study therefore demonstrated the absence of complementation of the native human *HsHK3* and the remarkable complementation by 22 out of 25 human genes of their yeast orthologs.

**Figure 1.**
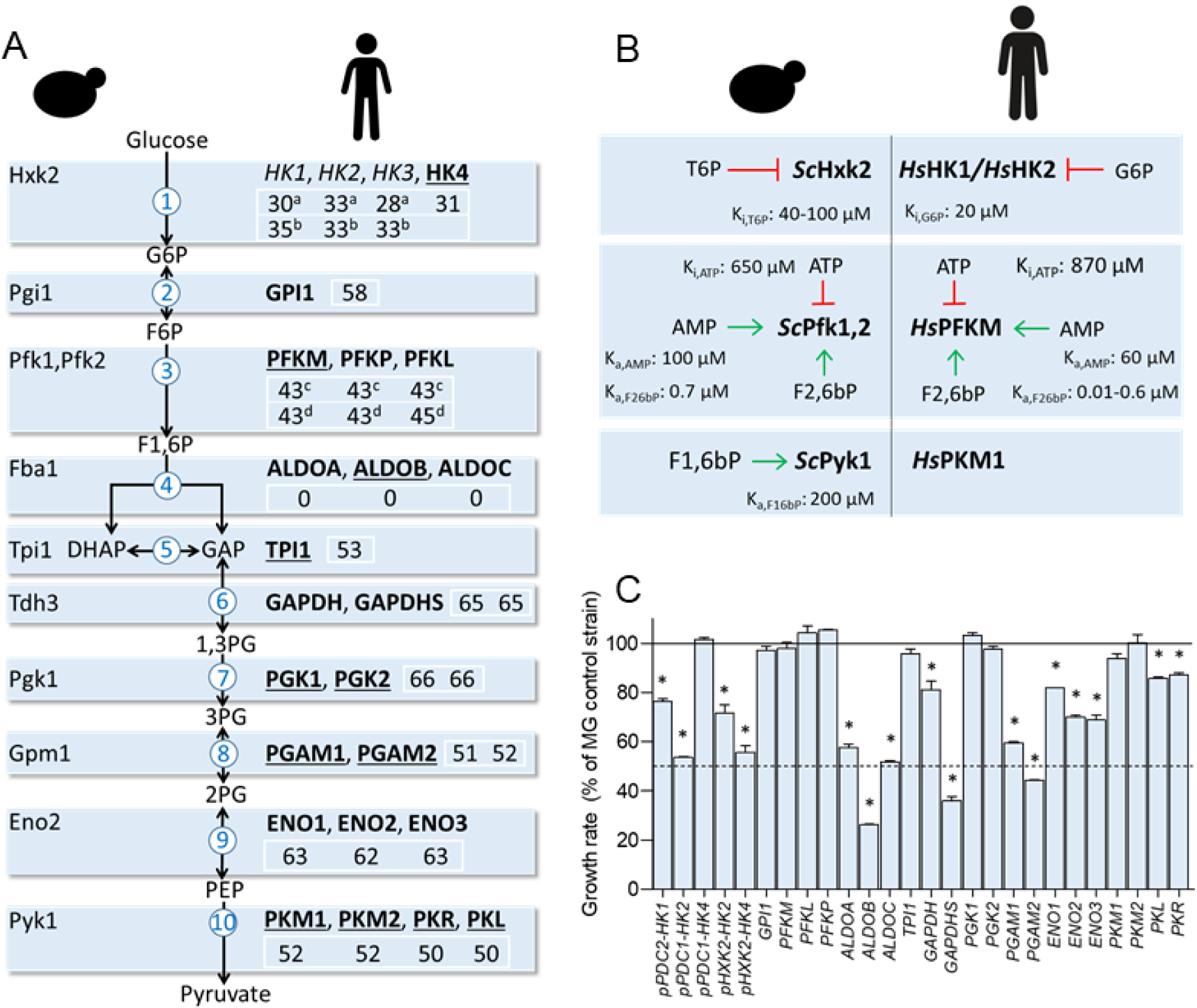
Glycolytic human and yeast enzymes relevant for this study and single complementation assays. **A)** Major glycolytic isoenzymes in yeast (left) and human enzymes used in this study (right). Underlined human enzymes were previously shown to complement their yeast ortholog. Bold enzymes complemented their yeast counterpart in this study. Percentage identity at protein level is shown of the human enzymes as compared to their yeast orthologs: ^a, b^, 1^st^ subunit and 2^nd^ of human hexokinases vs Hxk2, respectively, ^c,d^ human phosphofructokinases vs ScPfk1 and ScPfk2, respectively. Also see Table S1. **B)** Main allosteric regulators of the glycolytic yeast and human kinases and their regulation constants [36, 38, 84, 146–148]. **C)** Specific growth rates of the single gene complementation strains grown in SM glucose shown as percentage of the MG control strain IMX372, see also Fig S2. *HsHK2* and *HsHK4* were expressed with the *PDC1* promoter or the *HXK2* promotor, leading to different growth rates. Average and standard deviation of at least three independent replicates. * p-values between complementation and control strain below 0.01 (Student t-test, two-tailed, homoscedastic).

### Human HsHK1 and *Hs*HK2 can only complement the yeast hexokinases upon mutation

Upon transformation, strains expressing the human *HsHK1* or *HsHK2* as sole hexokinase grew well on galactose, a carbon source phosphorylated by galactokinase that does not require hexokinase activity, while exposure to glucose led to 1-2 days lag phase. Strains solely cultured on galactose displayed native *HsHK1* and *HsHK2* sequences, whereas exposure to glucose led to the systematic occurrence of single mutations in these genes, leading to an amino acid substitution or deletion (Fig. 2). *Hs*HK1 and *Hs*HK2 of strains solely exposed to galactose were active *in vitro* (IMX1689 and IMX2419 for *Hs*HK1 and *Hs*HK2 respectively), revealing that impaired growth on glucose was most likely not caused by lack of functionality of the human hexokinases in yeast (Fig. 2). Considering that native and mutated alleles of human hexokinases (strains IMS1137 and IMX1690) had similar catalytic activities *in vitro* (Fig. 2), growth defects upon exposure to glucose might result from inhibition of native human hexokinases in the yeast context. The observed mutations could then alleviate this inhibition to enable hexokinase activity *in vivo*. Mutations were observed in different regions of the *HsHK2* sequence in different strains, while in three separate *HsHK1* mutants the mutations were reproducibly localized at the glucose-6-phosphate binding site (Fig. 2). The activity of both human hexokinases is sensitive to substrate concentration [33, 34], but is also allosterically inhibited by the product of the reaction, glucose-6-phosphate (G6P) [34, 35]. The elevated intracellular G6P concentrations reported for yeast (0.5-2 mM) are well above the k_i,G6P_ of *Hs*HK1 and *Hs*HK2 (0.02 mM) and might inhibit these enzymes when expressed in yeast [36, 37].

**Figure 2.**
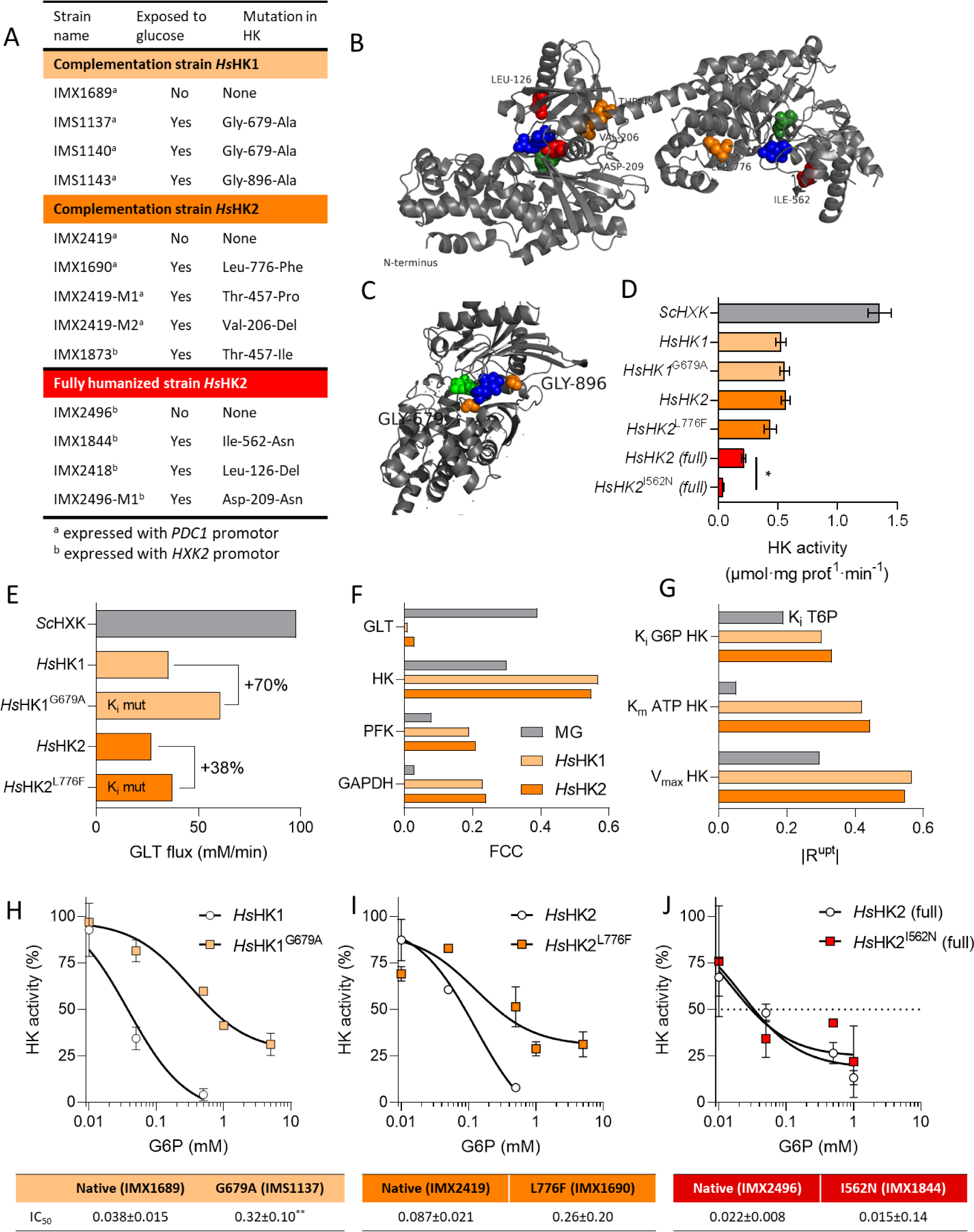
Characterization of human hexokinase mutants. **A)** Mutations in human hexokinases after growth on glucose. **B) and C)** Localization of the amino acid substitutions in *Hs*HK2 and *Hs*HK1 variants, respectively. Colour coding as in panel A. Green, glucose binding site in the catalytic domain and blue, glucose-6P allosteric binding site. *Hs*HK2 crystal structure from [33] and *Hs*HK1 from [114]. **D)** Hexokinase *in vitro* activity assay from *S. cerevisiae* strains grown on galactose. *Sc*Hxk2 control activity was measured using strain IMX2015. Non-mutated *Hs*HK1 and *Hs*HK2 were assayed using strains IMX1689, IMX2419, and IMX2496 (*Hs*Gly-HK2), that were never exposed to glucose. Activities of the mutated variants were measured in extracts obtained from IMS1137, IMX1690 and IMX1844. In the complementation strains and the control *Sc*Hxk strain, hexokinase was expressed with the strong *ScPDC1* promoter while in the fully humanized strains the *ScHXK2* promotor was used. * significant change in activity as compared to unmutated enzymes (p<0.01, n=2, unpaired t-test). **E)** Simulated glucose uptake rate (GLT, glucose transporter) for the MG control, native *Hs*HK1 and *Hs*HK2 complementation strains. The effect of the observed change in K_i_,G6P on the flux is modelled for each enzyme. **F)** Flux Control Coefficients (FCC’s) of the four enzymes with the highest control over the flux, GLT, hexokinase (HK), phosphofructokinase (PFK) and glyceraldehyde 3-phosphate dehydrogenase (GAPDH). **G)** Absolute values of the response coefficient |R^upt^| of the three parameters with highest control over the steady-state glucose uptake flux. **H)-I)** *In vitro* activity of the native and mutated hexokinase variants at various concentrations of the competitive inhibitor glucose-6-phosphate, expressed as percentage of activity without inhibitor. IC_50_: half maximal inhibitory concentration of G6P (mM). ** significant difference of the mutated as compared to the native variant (p<0.01, extra-sum-of-squares F test, n=2).

A computational ‘hexokinase complementation model’ built to address this phenomenon (Appendix 1) predicted a functional glycolytic pathway, able to reach a stable flux for both *Hs*HK1 and *Hs*HK2 with glucose as carbon source (Fig. 2E), but with a remarkable shift in the control of the glycolytic flux from glucose import (as predicted with yeast hexokinases) to hexokinase (Fig. 2F). The glycolytic flux was more specifically predicted to be sensitive to the magnitude of the K_i,G6P_, K_m,ATP_ and V_max_ of both hexokinases (Fig 2G). In agreement with this prediction, the tested hexokinase variants of both *Hs*HK1 and *Hs*HK2 (*Hs*HK1^G679A^ from IMS1137 and *Hs*HK2^L776F^) were less sensitive to G6P inhibition than the native alleles (Fig. 2H-I). For the *Hs*HK1^G679A^ mutation, our results are in direct contradiction to a previous study, where this mutation was found to have no impact on inhibition by the glucose-6P analog 1,5-anhydroglucitol-6P in purified *Hs*HK1 expressed in *E.coli* [38], which could be due to the different host organism. The K_i,G6P_ of the humanized computational glycolytic model was modified using our experimental data (see Fig. 2H-I) to mimic the response of the mutated *Hs*HK1^G679A^ and *Hs*HK2^L776F^ variants, resulting in a predicted increase of the glycolytic flux of 70% for *Hs*HK1^G679A^ and 38% for *Hs*HK2^L776F^, as compared to their native variants (Fig. 2E). Conversely, the sensitivity of the native and L776F variants to ATP, ADP and glucose measured *in vitro* were similar (Fig. S4) and trehalose-6-phosphate, a major inhibitor of yeast hexokinase not present in human cells, only mildly affected human hexokinase activity *in vitro* and the predicted *in silico* glycolytic flux (Fig S4, Appendix 1) [38]. Glucose-6-phosphate inhibition is therefore most likely the main mechanism underlying the inability of native *Hs*HK1 and *Hs*HK2 to complement their yeast ortholog.

### Successful humanization of the entire glycolytic pathway in yeast

The successful complementation of individual glycolytic enzymes suggested that transplantation of a complete human glycolytic pathway might be possible. The success of humanization at full pathway level would depend on a combination of expression, kinetic and moonlighting properties of the whole set of enzymes (Fig. 1B) [39, 40], which together may have a severe impact on the growth of the yeast host. The muscle glycolytic pathway, characterized by fast *in vivo* rates, was chosen for transplantation in yeast [41–43]. While *Hs*HK1 and *Hs*HK2 are the most abundant enzymes in muscle tissue [44, 45], *Hs*HK4 was initially chosen as hexokinase due to its ability to readily complement *Sc*Hxk2 (Fig. 1). *Hs*HK2 was also transplanted in a second glycolysis version, to test whether mutations were also required for its functionality in a human glycolytic context. Using both *Hs*HK4 and *Hs*HK2 was also interesting considering their difference in sequence and in kinetic and regulatory properties, *Hs*HK4 having a substantially lower affinity for glucose and being insensitive to glucose-6P [30, 46, 47]. Transplantation in a SwYG strain [28] of the entire set of human glycolytic genes resulted in the *Hs*Gly-HK2 strain with *Hs*HK2 as hexokinase and the *Hs*Gly-HK4 strain with *Hs*HK4 (Fig. 3). Expression of the human genes was driven by strong, constitutive yeast promoters (Table S2). Note that expression of *HsHK2* and *HsHK4* using the yeast *ScHXK2* promoter in these fully humanized strains, led to a slower growth rate than complementation using the stronger *ScPDC1* promoter (Fig. 1 and Fig S2). Pathway transplantation was successful as both the *Hs*Gly-HK2 and *Hs*Gly-HK4 strains displayed remarkably fast growth (ca. 0.15 h^-1^, around 40% of the control SwYG strain with native glycolysis *Sc*Gly (IMX1821)) in minimal medium with glucose as sole carbon source (Fig. 3). Exposure to glucose of the *Hs*Gly-HK2 strain led to long lag phase, and sequencing of *HsHK2* from several culture isolates revealed the systematic presence of single mutations in the vicinity of the catalytic and glucose-6P binding sites (Fig. 2, Table S3). Remarkably a *ca.* 5-fold decrease in hexokinase V_max_ was observed in an isolate carrying the *Hs*Gly-HK2^I562N^ variant, while no evidence was found for other changes in kinetic parameters (Fig 2 and Fig S5). In another isolate, the *Hs*HK2^D209N^ mutation affected an amino acid key to the activity of the C-terminal active site [34]. The decrease in V_max_ but maintenance of glucose-6P sensitivity of *Hs*HK2^I562N^ was in stark contrast with the stable V_max_ but decreased glucose-6P sensitivity observed for *Hs*HK2 variants in single complementation strains (Fig. 2). This suggested that the humanized glycolytic context might result in a different intracellular environment (particularly metabolite concentrations) and thereby lead to different requirements for hexokinase functionality and different evolutionary solutions.

**Figure 3.**
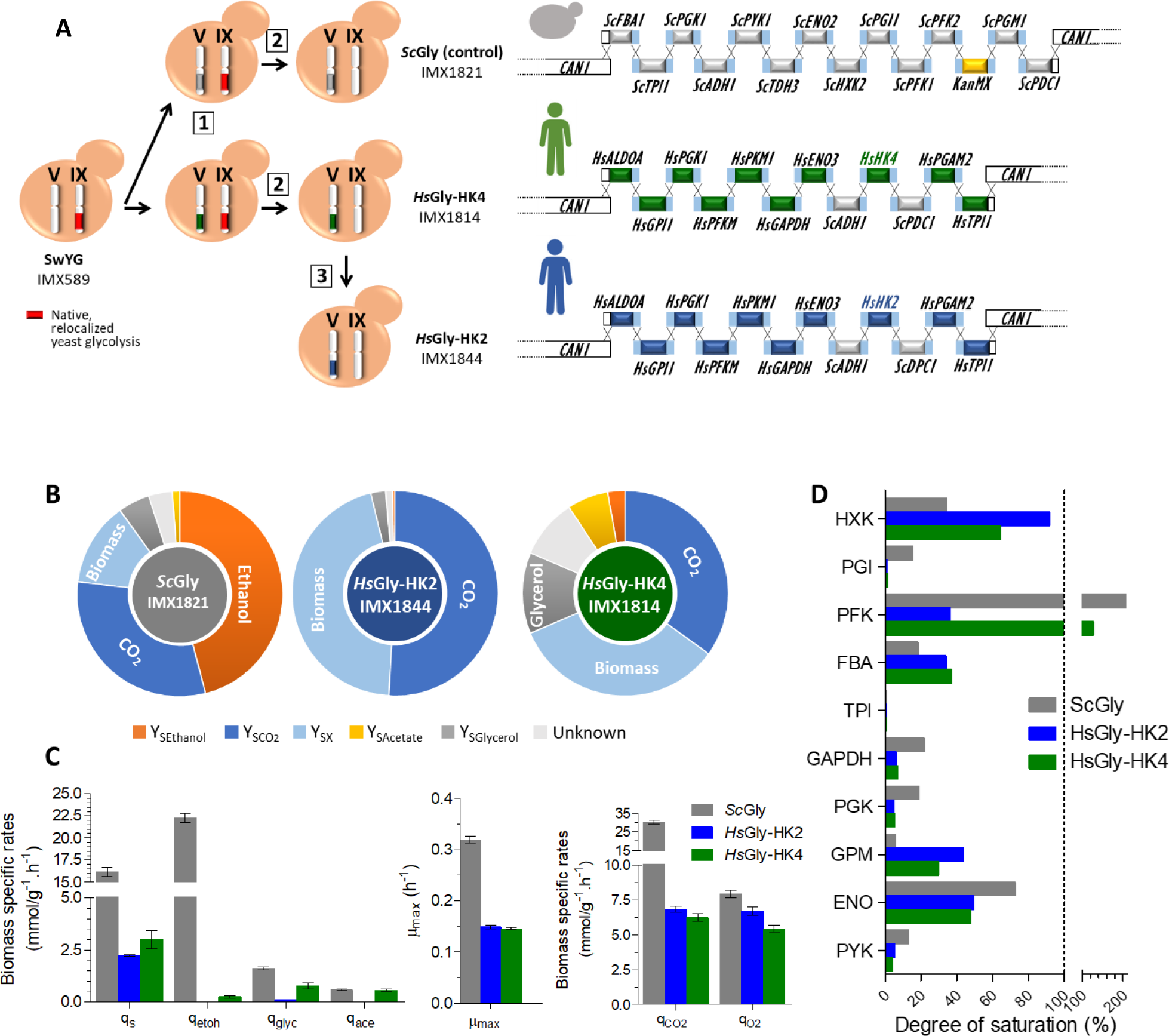
Construction and physiological characterization of strains with fully humanized glycolysis. **A)** Strain construction strategy and glycolytic pathway strains with native, and humanized co-localized glycolysis. **B), C) and D)** Physiological characterization of strains shown in panel A in bioreactors on SM glucose. **B)** Yields on glucose (CMol/CMol) of ethanol, CO_2_, biomass, acetate and glycerol are indicated (Y_SEthanol_, Y_SCo2_, Y_SX_, Y_SAcetate_, Y_SGlycerol_ respectively). **C)** Specific rates for glucose and oxygen uptake (q_glu_ and q_O2_), and for ethanol (q_Eth_), glycerol (q_gly_), acetate (q_acet_), CO_2_ (q_CO2_) and biomass (µ_max_) production. Average and SEM of biological duplicates. **D)** Estimation of the degree of saturation of glycolytic enzymes based on *in vitro* assays from cell extracts (activities reported in Fig. S8). *in vivo* fluxes were approximated from the q_glu_. The dashed line indicates 100% saturation.

Next to hexokinase, human and yeast pyruvate kinases also differ in allosteric regulations, *Hs*PKM1 being insensitive to the feed-forward activation by F1,6bP characteristic of *Sc*Pyk1 (Fig. 1B, Fig. S6). Despite the proposed role of this allosteric regulation for yeast cellular adaption to transitions [40, 48], the ability of the humanized yeast strains was not visibly impaired during transition between alternative (galactose) and favourite (glucose) carbon source (Fig. S7).

*S. cerevisiae* favors a mixed respiro-fermentative metabolism when glucose is present in excess (see IMX1821 in Fig. 3), a phenomenon known as the Crabtree effect, analogous to the Warburg effect in mammalian cells [49]. *Hs*Gly-HK2 mostly respired glucose, with only traces of ethanol and glycerol being produced, while *Hs*Gly-HK4 displayed a respiro-fermentative metabolism, more similar to that of the *Sc*Gly control strain, although with far lower substrate uptake and ethanol production rates (Fig. 3 and Table S4). The fact that these physiological responses were similar to those observed for the respective *Hs*HK2 and *Hs*HK4 single complementation strains suggested that the human hexokinases strongly contributed to this switch between fermentative and respiratory metabolism, but they might not be the only players (Fig. 4A). Excepted *Hs*GPI1, the activity of the human enzymes was two to fifty times lower than the activity of their yeast ortholog (Fig. 3 and Fig. S8). With the notable exception of phosphofructokinase, sensitive *in vivo* to many effectors, the yeast glycolytic enzymes generally operate at overcapacity ([50–52] and Fig. 3). In the humanized yeast strains hexokinase, aldolase and phosphoglycerate mutase showed higher degrees of saturation compared to the control strain with the native yeast glycolysis, suggesting that the activity of these enzymes could exert higher control on the glycolytic flux in the humanized strains (Fig. 3). In line with this hypothesis, these three enzymes also led to low growth rates in single complementation strains (Fig. 1). Remarkably, the activity of *Hs*PFKM was 2.6-fold lower in *Hs*Gly-HK4 than in *Hs*Gly-HK2, while the same protein abundance was found (Fig. S8). Consequently, while *Hs*PFKM *in vivo* operated above its *in vitro* capacity in *Hs*Gly-HK4, similarly to what is typically observed in *S. cerevisiae* and in IMX1821, the flux through *Hs*PFKM in *Hs*Gly-HK2 was only at ca. 30% of its *in vitro* capacity (Fig. 3).

**Figure 4.**
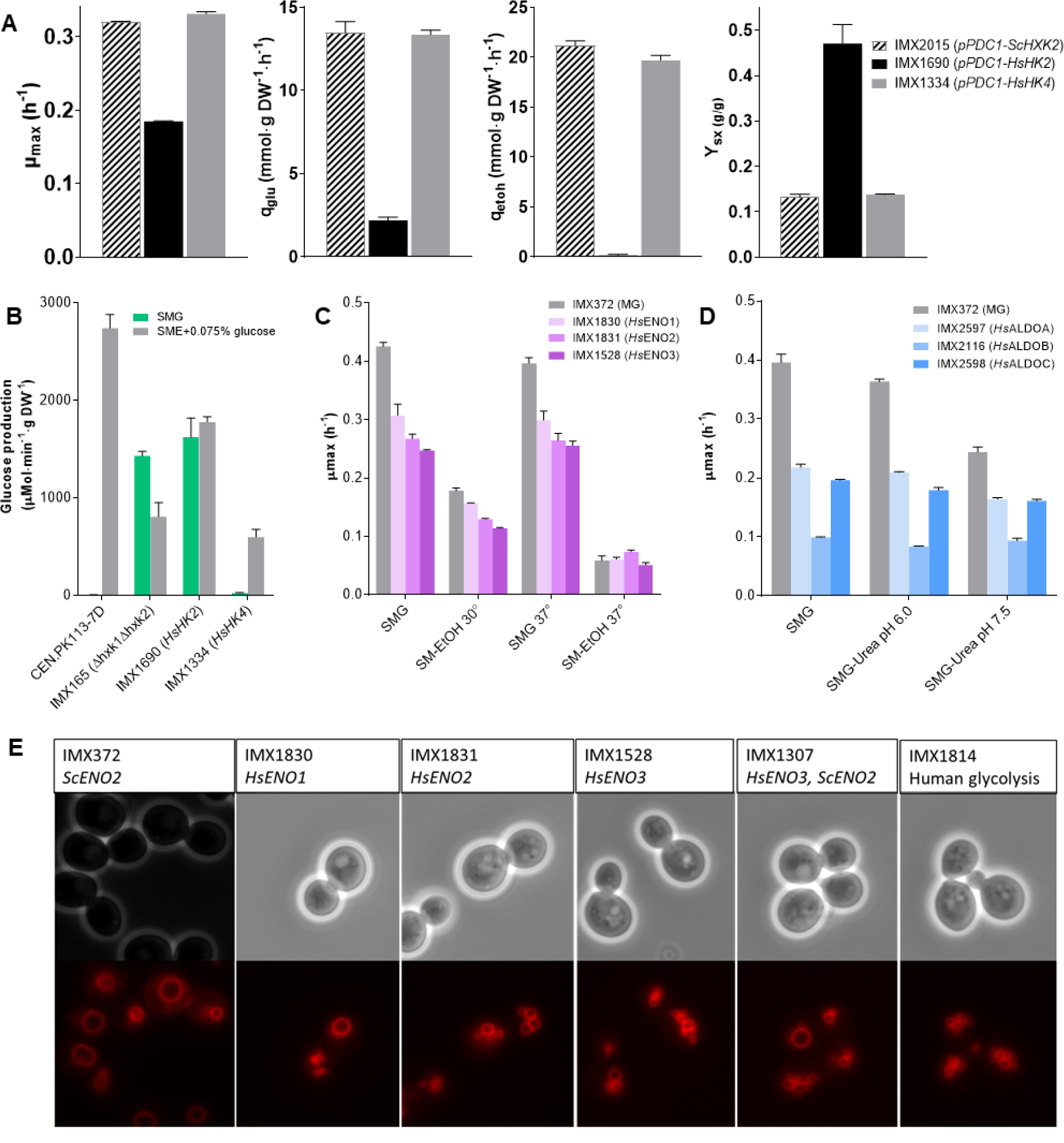
Complementation of moonlighting functions. **A)** Specific growth (µ_max_), glucose consumption (q_glu_) and ethanol production (q_Eth_) rates, and biomass yield (Y_SX_) of single hexokinase complementation strains grown in shake flask in SM glucose. Average and SEM of two biological replicates. **B)** Extracellular invertase activity of cultures with SM glucose (repressing condition) or SM ethanol + 0.075% glucose (inducing condition). Average and SEM of two biological replicates. **C)** and **D)** Specific growth rate of strains with single complementation of the three human enolases and the three human aldolases, and of the MG control strain at 30 °C and 37 °C, with glucose or ethanol as carbon source or at different pH as indicated. Average and standard deviation of biological triplicates. SM medium was used, but ammonium was replaced by urea to maintain pH in panel D). **E)** Staining of membranes with FM4-64 in *S. cerevisiae* strains expressing the yeast *Sc*Eno2 ( IMX372 control) or the human *Hs*ENO3, *Hs*ENO2 and *Hs*ENO3 (IMX1830, IMX1831, IMX1528) as single complementation or as fully humanized glycolysis (*Hs*ENO3, IMX1814). IMX1307 carries one copy of the human *HsENO3* and the yeast *ScENO2* gene.

Overall the transplantation of a complete human glycolytic pathway to yeast was successful despite the structural, kinetic and regulatory differences between yeast and human enzymes. Global proteomics revealed increases in protein abundance of mainly metabolic enzymes, corresponding to the altered physiology (Fig. S9). The fully humanized glycolysis strains grew remarkably well, where the reduced glycolytic flux and growth rate as compared to yeast strains with a native glycolysis (μ_max_ ca. 60% slower), were in agreement with the lower *in vitro* enzymatic capacity of the human glycolytic enzymes.

### Complementation of moonlighting functions

Many eukaryotic glycolytic enzymes, next to their glycolytic functions, have other cellular activities. These moonlighting functions might not be conserved across species [26] and failure of the human orthologs to complement the yeast moonlighting activities could strongly affect the humanized yeast strains. Three glycolytic enzymes in *S. cerevisiae* have documented moonlighting functions: hexokinase, aldolase and enolase.

*Sc*Hxk2 is involved in glucose repression and the Crabtree effect by partially localizing to the nucleus in the presence of excess glucose where it represses the expression of genes involved in respiration and the utilization of alternative carbon sources such as the sucrose hydrolysing invertase *SUC2* [53, 54]. Accordingly, invertase activity is not detected in *S. cerevisiae* cultures with excess glucose, while it is expressed and active when glucose repression is alleviated (Fig. 4B). Conversely, double deletion of *ScHXK1* and *ScHXK2* alleviates glucose repression and enables invertase expression and activity in the presence of excess glucose condition. Invertase assays suggested that *Hs*HK4 but not *Hs*HK2^L776F^ was able to complement the role in glucose repression of *Sc*Hxk2 (Fig. 4). However, since invertase repression is known to be sensitive to growth rate and the *Hs*HK2-Gly strains grew slowly, our findings do not completely rule out the possibility that *Hs*HK2 plays a role in glucose repression, although based on the difference in sequence, size and structure with yeast hexokinase, this seems unlikely [56]. The role of *Hs*HK4 in glucose repression, suggested in an earlier report [24], is in line with the Crabtree effect we observed for the complementation and fully humanized strains carrying *Hs*HK4 despite their slow growth rate (Fig 3, [55–57]).

Yeast aldolase is involved in assembly of vacuolar proton-translocating ATPases (V-ATPases), leading to the inability of aldolase deficient strains to grow at alkaline pH [58]. This function has been reported to be highly conserved between the yeast Fba1 and the human *Hs*ALDOB despite the absence of any sequence homology between the two proteins [58, 59]. The present study shows that the other human aldolases (*Hs*ALDOA and *Hs*ALDOC), which share ca. 70% identity with *Hs*ALDOB, can complement the moonlighting functions of *Sc*Fba1. All three human aldolase complementation strains as well as the fully humanized strains *Hs*Gly-HK2 and *Hs*Gly-HK4 showed no growth defects at pH 7.5 (Fig. 4 and Fig. S10), indicating that the vacuolar function was complemented by all three human aldolases.

Furthermore, the yeast enolases *Sc*Eno1 and *Sc*Eno2, are involved in vacuolar fusion and protein transport to the vacuole [60]. Whether the human enolases can take over this function in yeast is unknown. While enolase-deficient yeast strains display a fragmented vacuole phenotype and growth defects, this phenotype was not observed for the MG strain expressing *Sc*Eno2 only ([61], (Fig. 4) and was also not observed for complementation strains expressing any of the three human enolases (Fig. 4). This vacuolar moonlighting function seems therefore to be conserved between yeast and human enolases. Additionally the yeast enolase plays a role in mitochondrial import of a tRNA^Lys^, a mechanism important at growth temperature above 37°C, particularly on non-fermentable carbon sources [62, 63]. This mechanism seems well conserved in mammals, as yeast tRK1 is imported *in vitro* and *in vivo* in human mitochondria, in an enolase-dependent manner [64–66]. All three human enolase complementation strains show only minor growth defects at 37°C on both glucose and non-fermentable carbon sources compared to the MG control strain (Fig. 4). The fully humanized strains similarly show no growth defect at 37°C (Fig. S10). This suggests that, in addition to the vacuolar function of *Sc*Eno2, the human enolase enzymes are also able to fully take over its role in mitochondrial import of tRK1.

With the exception of *Hs*HK2, no phenotypic defect could be observed in the humanized strains, suggesting that the moonlighting functions of all glycolytic enzymes could be complemented by their human orthologs.

### Engineering and evolutionary approaches to accelerate the slow growth of humanized glycolysis strains

Several enzymes (*Hs*HK2, *Hs*HK4, *Hs*ALDOA and *Hs*PGAM2) showed a significantly higher degree of saturation in the humanized strains (two to six-fold higher as compared to *Sc*Gly, Fig. 3). The corresponding complementation strains also grew slower than the control strain, suggesting that the capacity of these enzymes might be limiting the glycolytic flux. Indeed simultaneous overexpression of *HsHK2*, *HsALDOA* and *HsPGAM2* in *Hs*Gly-HK2, and of *HsHK4*, *HsALDOA* and *HsPGAM2* in *Hs*Gly-HK4 successfully increased their specific growth rate by 63% and 48% respectively (Fig. 5). These optimized, humanized yeast strains still grew 30% to 40% slower than the control strain with native, minimized yeast glycolysis (Fig. 5). Growth at 37°C, optimal temperature for human enzymes, instead of 30°C did not improve the growth rate of the humanized yeast strains (Fig. 5A, also Fig. S10).

**Figure 5.**
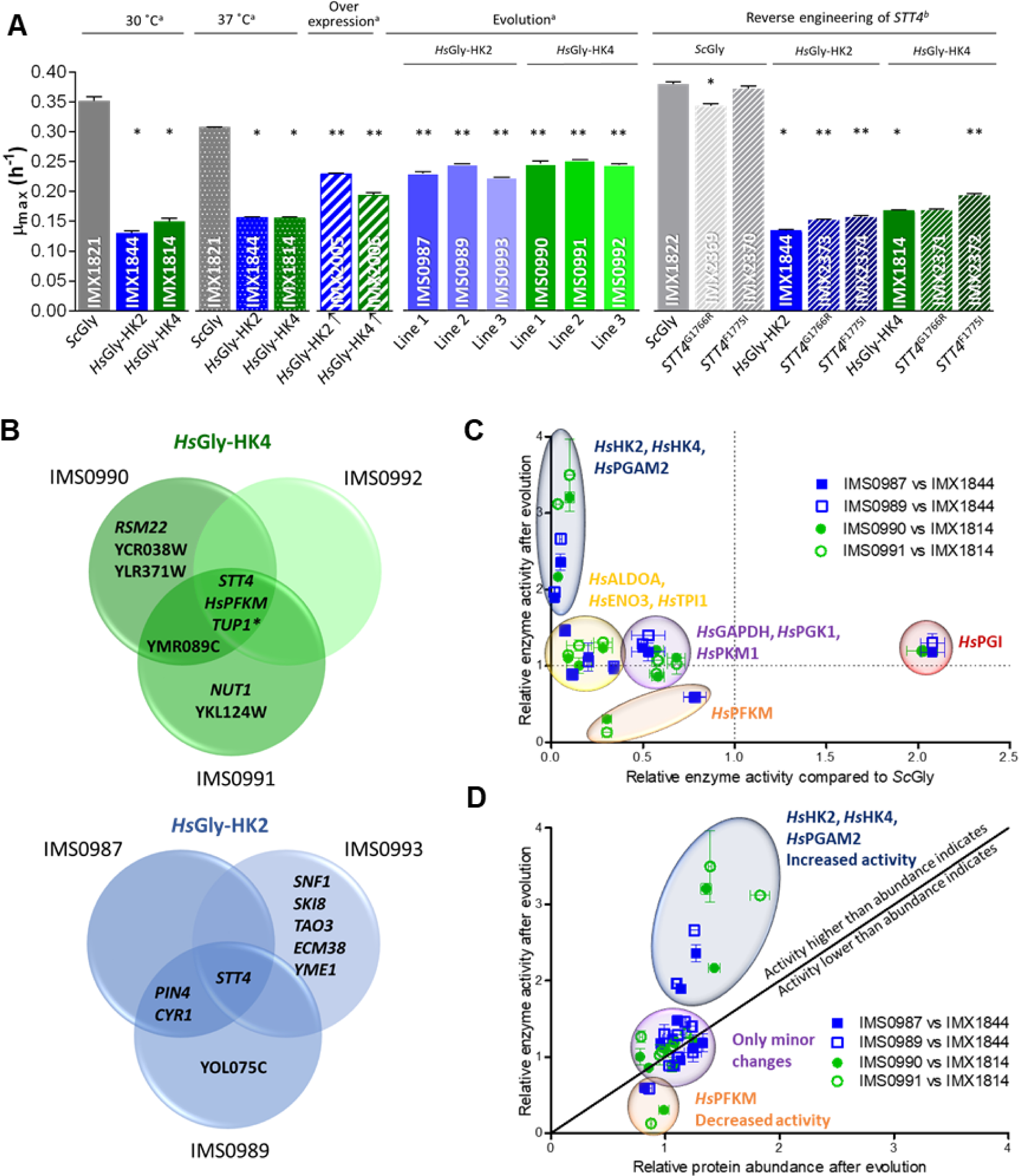
Strategies to improve the growth rates of fully humanized yeast strains. **A)** Specific growth rate of humanized, evolved and reverse-engineered strains in SM glucose. * indicates significant differences between *Hs*Gly-HK2 or *Hs*Gly-HK4 and the control strain IMX1821 and ** between the evolved and reverse-engineered strains and their respective parental strain *Hs*Gly-HK2 or *Hs*Gly-HK4 (Student t-test, two-tailed, homoscedastic, p-value<0.05). ^a^ aerobic shake-flasks, ^b^ Growth Profiler. **B)** Mutations found in single colony isolates from independent evolution lines of humanized yeast strains. **C)** Comparison between the changes in glycolytic enzyme activity caused by humanization of yeast glycolysis and by evolution of the humanized strains. Activities available in Fig. S8 and S13. Error bars represent SEM. Enzymes with a similar response are grouped. **D)** Comparison of changes in enzyme activity and in protein abundance caused by evolution of the humanized yeast strains.

Many other mechanisms could explain the slow growth phenotype of the humanized strains, such as (allosteric) inhibition of the enzymes *in vivo*, incompatibility of substrate and co-factors concentrations with enzyme kinetic requirements, or deleterious effects of moonlighting activities of the human orthologs, more than could reasonably be tested by design-build-test-learn approaches. An adaptive laboratory evolution (ALE) strategy, particularly powerful to elucidate complex phenotypes [67], was therefore used to improve the fitness of the humanized strains. After approximately 630 generations in glucose medium, evolved populations of humanized yeast strains grew ca. twofold faster than their *Hs*Gly-HK2 and *Hs*Gly-HK4 ancestors (Fig. S11). Single colony isolates from six independent evolution lines, three per humanized yeast strain, confirmed the increased growth rate of the evolved humanized yeast strains (strains IMS0987 to IMS0993, Fig. 5, Table S7). These strains evolved towards a higher glycolytic flux and a more fermentative metabolism, producing ethanol, albeit with a lower yield compared to the *Sc*Gly control (Fig. 5 and Fig. S12). This increase in fermentation was in line with the increased specific growth rate and glucose uptake rate [55]. These evolved strains were further characterized in an attempt to elucidate the molecular basis of the slow growth phenotype of the humanized yeast strains.

### Exploring the causes of the slow growth phenotype of humanized glycolysis strains

The activity of several human glycolytic enzymes was affected by evolution (Fig. S13). Across the six evolution lines, the activity of both hexokinases (*Hs*HK2 and *Hs*HK4) and *Hs*PGAM2, for which activity was much lower than their yeast variants, was increased two to three-fold during evolution (Fig. 5C). These enzymes also led to the strongest decrease in growth rate upon complementation (Fig. 1 and Fig. S14). Protein abundance of *Hs*HK4 and *Hs*PGAM2 was accordingly increased, albeit not with the same magnitude, but the change in *in vitro* activity of *Hs*HK2 was not reflected in protein abundance (Fig. 5D, Fig S13). The activity of *Hs*ALDOA, which also led to a large decrease in growth rate upon complementation and for which the activity was strongly reduced in the fully humanized strains, was not markedly altered by evolution. The response of *Hs*PFKM was particularly interesting. While its activity was already lower in the humanized yeast strains than in the *Sc*Gly strain, *Hs*PFKM was the only enzyme for which the activity was substantially decreased during evolution, by a factor of 2 to 8 as compared to their non-evolved humanized ancestors. For *Hs*HK2 and *Hs*PFKM, the changes in *in vitro* activity were not reflected in protein abundance. Global proteomics showed few proteins changed significantly in abundance after evolution (Fig. S15).

With the exception of phosphofructokinase, the genome sequence of the evolved strains offered little insight into the mechanisms leading directly to the above-mentioned alterations in *in vitro* specific activities or abundance of the glycolytic enzymes. However, interesting mutations were identified. The promoter, coding or terminator regions of the human glycolytic genes were exempt of mutations in the evolved strains. Only *Hs*PFKM carried a single mutation in its coding region in all three evolution lines of *Hs*Gly-HK4 (Fig. 5, Table S5), one located in the N-terminal catalytic domain of the protein and the two others in the C-term regulatory domain where several allosteric effectors can bind (F2,6biP, ATP, ADP, citrate, etc. [68, 69]). The impact of these mutations cannot be inferred directly from the location, but they are most likely involved in the strong decrease in *in vitro* activity of PFKM in the strains evolved from *Hs*Gly-HK4. All three evolved strains from the *Hs*Gly-HK4 strain were also mutated in *TUP1* (general repressor of transcription with a role in glucose repression), with a conserved non-synonymous mutation resulting in an amino acid substitution (Fig. 5, Table S5). Other transcription factors involved in the regulation of the activity of the yeast glycolytic promoters (Rap1, Abf1, Gcr1, Gcr2) did not harbour mutations in any strain. Overall few mutations were conserved between the evolution lines of the two humanized strains, but mutations in a single gene, *STT4*, were found in all six evolution lines (Fig 5). Remarkably the six identified mutations were located within 164 amino acids, in the C-terminus of the protein harbouring its catalytic domain (Table S5). *STT4* encodes a phosphatidylinositol-4P (PI4P) kinase that catalyses the phosphorylation of PI4P into PI4,5P2. As Stt4 is essential in yeast [70], the mutations present in the evolved strains could not cause a loss of function. Phosphoinositides are important signalling molecules in eukaryotes, involved in vacuole morphology and cytoskeleton organisation via actin remodelling [71]. PI4P kinases are conserved eukaryotic proteins [72], and Stt4 shares similarities with human PI3 kinases [73]. In mammals, activation of PI3K remodels actin, thereby releasing aldolase A trapped in the actin cytoskeleton in an inactive state and increasing cellular aldolase activity [74, 75]. As yeast and human forms of actin are highly conserved (89% identity at protein level), a similar mechanism could be active in yeast and enable the evolved, humanized yeast strains to increase aldolase activity *in vivo* without increasing its concentration. Reverse engineering of two of the mutations found in the evolved strains IMS0990 and IMS0992 was performed in the non-evolved strain backgrounds with native yeast glycolysis and humanized glycolysis, by mutating the native *STT4* gene (Fig. 5). The increases in specific growth rate did not match the growth rates of the evolved strains, suggesting other parallel mechanisms. Interestingly, in the reverse engineered strains *STT4* mutations resulted in a fragmented vacuole phenotype (Fig. S16), confirming that the mutations interfered with Stt4 activity and PI4P signalling. Such a phenotype was not observed in the evolved strains, however in these strains vacuoles also displayed abnormal morphologies with collapsed structures, indicating that specific mechanisms might have evolved in parallel to mitigate the effect of *STT4* mutations on vacuolar morphology (Fig. S16).

These findings suggest that evolution led to optimization of the human glycolytic pathway function in yeast through several mechanisms. Hexokinase 4 and phosphoglycerate mutase abundance and activity increased, allowing a higher glycolytic flux. For hexokinase 2 and phosphofructokinase, posttranslational mechanisms to modify enzyme activity must be present, counter-intuitively decreasing *Hs*PFKM activity *in vitro*. In all evolution lines, mutations in Stt4 occurred, which could potentially benefit *in-vivo* aldolase activity through modulation of actin structures. These adaptations reveal that the enzymes with the largest impact on growth rate in single complementation models (*Hs*HK2, *Hs*HK4, *Hs*PGAM2 and *Hs*ALDOA), and not those with the lowest activity, are the main targets for evolution. Changes in enzyme abundance, cellular environment and posttranslational modifications, and not direct mutations of the glycolytic genes, appear to be the most effective evolutionary strategy to improve flux of this heterologous pathway.

### Relevance of yeast as model for human glycolysis

The yeast intracellular environment could interfere with folding or posttranslational modifications of human enzymes and thereby alter their catalytic turnover number (k_cat_). To explore this possibility, the k_cat_ of the human glycolytic proteins in yeast and in their native, human environment in myotube was experimentally determined. Enzyme activities (V_max_) were measured in cell extracts using *in vivo*-like assay conditions. In these assays phosphofructokinase and hexokinase activities were too low for detection, although both could be detected at the protein level (Fig. S17). Overall the V_max_ of glycolytic enzymes were of the same order of magnitude in humanized yeast and muscle cells (Fig. 6A). *Hs*GPI1, *Hs*ALDOA, *Hs*GAPDH and *Hs*PGK1 activity was higher in yeast cells than in muscle cells, particularly for *Hs*GPI1 (seven-fold), while the activity of *Hs*PGAM2 was 5.5 fold lower in yeast compared to muscle cells (Fig. 6A). The differences in *in vitro* activity between yeast and human isoenzymes were mirrored in the peptide abundance for these proteins (Fig. S17), suggesting that the turnover rates of the human proteins expressed in human and yeast cells were not substantially different.

**Figure 6.**
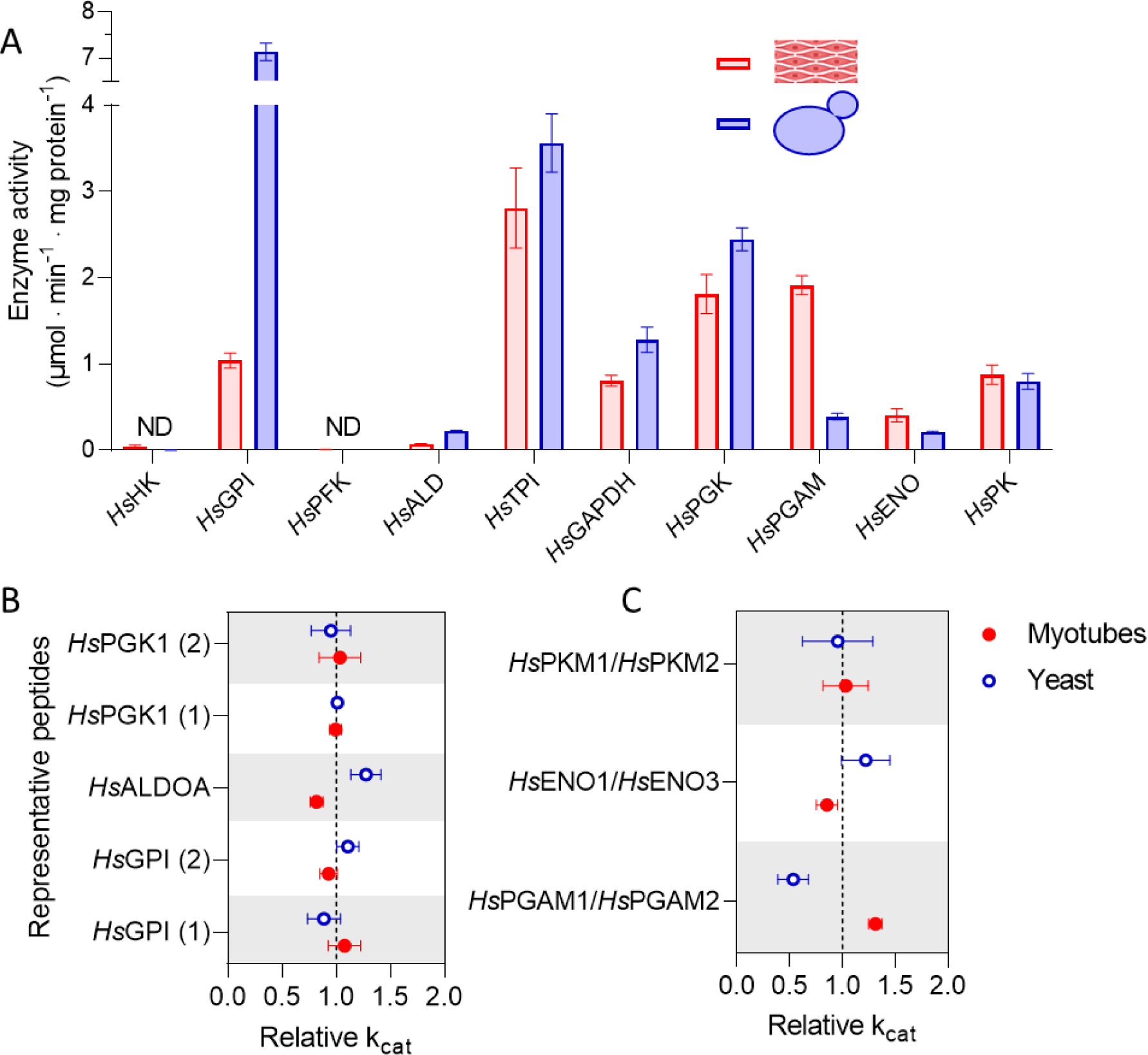
Enzyme activity and k_cat_ of human glycolytic enzymes in human myotubes and humanized yeast. **A)** Specific activity of human glycolytic enzymes measured *in vitro* with *in-vivo* like assaying conditions. Blue, yeast strain *Hs*Gly-HK2, red, muscle myotube cultures. **B)** and **C)** k_cat_ values for enzymes present as single isoform **B)** or multiple isoforms **C)** in the myotubes proteome data (Fig. S17). Data represent the mean values and standard deviation of three and two independent culture replicates for the myotubes and yeast cultures respectively. ND: not detected.

For *Hs*GPI, *Hs*ALDOA and *Hs*PGK1, the k_cat_ values were calculated by dividing the V_max_ values by the respective protein concentrations (Fig. S17C). This revealed no differences in the turnover rate between yeast and myotubes, irrespective of which of the standard peptides was used for protein quantification (Fig. 6B). For the remaining enzymes, calculation of the turnover rate was complicated by the presence of isoenzymes other than the canonical muscle glycolytic enzymes in myotube cultures. In addition to the canonical muscle isoenzymes, the isoenzymes *Hs*PFKL, *Hs*PGAM1, *Hs*ENO1 were present at equivalent or higher concentrations and *Hs*HK1, *Hs*PFKP and *Hs*PKM2 were present in low concentrations (Fig S17). This difference in isoenzyme abundance between tissue and isolated cell lines has been reported before for *in vitro* muscle cultures and, to a lower extent, for muscle biopsies [76]. Therefore, for enolase, phosphoglycerate mutase and pyruvate kinase, the apparent k_cat_ was assumed to be the V_max_ divided by the sum of all detected isoforms catalysing the specific reaction. The k_cat_ values of *Hs*ENO1 and *Hs*ENO3 are reported to be similar [77], but the *Hs*PGAM2 and *Hs*PKM1 have a higher k_cat_ than their respective isozymes [78, 79]. Taking these proportions into account, we found that apparent k_cat_ values for enolase and pyruvate kinase were similar between humanized yeast and myotubes while the k_cat_ of *Hs*PGAM was lower in humanized yeast (2.5 fold lower than the value in myotubes) (Fig. 6). This may suggest that the yeast intracellular environment has a negative impact on posttranslational processing of the enzyme or the influence of the *Hs*PGAM2 isozyme in muscle cells. Taken together, these results demonstrate that out of the six enzymes for which a turnover rate could be determined, five were not catalytically altered by the yeast environment, with *Hs*PGAM2 as potential exception.

## Discussion

The human glycolytic genes showed remarkable levels of complementation in yeast, both individually and as a complete pathway, with conservation of their secondary functions and turnover numbers similar to human muscle glycolysis. The combination of strains presented here can thus serve as new models to study fundamental aspects of human glycolysis in a simplified experimental setting, including moonlighting functions, the effects of PTMs and allosteric regulators, and cross-talk between enzymes. The extensive genetic accessibility and tractability of yeast enables the fast construction and testing of libraries of humanized yeast strains that carry different glycolytic designs.

The 100 kDa human hexokinases 1 to 3 did not show immediate complementation, however for *Hs*HK1 and *Hs*HK2, single amino acid substitutions were sufficient to restore their functionality in the yeast cellular environment. The requirement for mutations illustrates that these hexokinases have evolved to function in a particular metabolic niche. The ease of complementation with the human glycolytic genes is remarkable since glycolytic enzymes are known to be involved in numerous different moonlighting functions in yeast and human cells. In line with earlier work [24], we found that *Hs*HK4 but not *Hs*HK2 is able to complement the yeast Hxk2 function in invertase repression. The ability of *Hs*HK4 to transduce glucose signalling in yeast is surprising since this enzyme is not reported to have a transcriptional regulation function in human cells. *Hs*HK4 also lacks the decapeptide required for the translocation of *Sc*Hxk2 to the nucleus and its binding to the Mig1 transcription factor, although it has previously been shown to localize to the yeast nucleus [54, 80, 81]. The secondary functions of yeast aldolase and enolase in vacuolar ATPase assembly, vacuolar fusion and transport, and mitochondrial tRNA import were complemented by all human aldolase and enolase isozymes. This extraordinary conservation of glycolytic moonlighting functions observed between human and yeast glycolytic enzymes challenges our understanding of the underlying molecular mechanisms and reveals evolutionarily conserved functions.

The successful humanization of the entire glycolytic pathway in yeast and the availability of a library of strains with single complementation offer a unique opportunity to study potential synergetic effects between glycolytic enzymes and the impact of a full pathway on individual enzymes. A good example is the different evolutionary strategies found by fully humanized and single complementation strains to restore functionality of human hexokinases in a yeast context. All tested single complementation strains alleviated G6P inhibition on *Hs*Hk1 and *Hs*Hk2, while a fully humanized strain reduced *Hs*Hk2 activity without altering G6P sensitivity. G6P is a key metabolite at the branchpoint of several pathways in both yeast and human. While the capacity in glycogen synthesis and hexokinase activity, as measured by their V_max_’s, does not largely differ between yeast and skeletal muscle [82–86], several yeast to muscle differences could account for the higher cellular G6P levels found in yeast [36, 37]. Critically, glucose uptake in yeast and muscle cells is very differently regulated. In the skeletal muscle, glycolytic fluxes are very dynamic and respond to the physiological status (such as meal status and exercise), a response largely controlled by glucose transport [87, 88] and ATP demand [89]. Even at their maximum capacity upon stimulation by insulin, glucose uptake is *ca*. two orders of magnitude lower than in yeast cells, and the muscle environment offers a much higher phosphorylation/uptake ratio than yeast. Other differences influencing the G6P concentration could be the presence in yeast of the trehalose cycle [90], and the greater capacity in yeast of glucose 6-phosphate dehydrogenase, first step of the pentose phosphate pathway, compared to human muscle [82, 91–93]. G6P has also been implicated in transcriptional regulation via the ChREBP and MondoA-MlX transcription factor complexes, which in turn modulate glycolytic gene expression in human cells [94, 95]. Altogether these factors contribute to yeast to skeletal cells differences in cellular G6P levels, resulting in supra-inhibitory levels for *Hs*HK1 and *Hs*HK2 in the yeast context. The different evolutionary strategy found in strains with fully humanized glycolysis might originate from different factors. The isomerization of G6P into F6P does most likely not account for large differences in G6P levels in fully humanized and single complementation strains as both human *Hs*PGI and native *Sc*Pgi1 operate near-equilibrium and have similar *in vitro* activity [96]. Conversely, the substantially lower activity of several human glycolytic enzymes as compared to their yeast equivalent (e.g. ALDOA and PGAM activities are ca. 10-fold) and resulting low glycolytic flux might alter the yeast cellular context (i.e. metabolite concentrations), and thereby the selection pressure exerted on hexokinase. Measuring intracellular metabolites in the different humanized strains should shed some light on the impact of partial and full humanization on the yeast cellular context. Another difference between human and yeast hexokinase 2 is the VDAC-dependent mitochondrial binding of the human variant, a binding not likely to be conserved in the humanized strains, unless the human VDAC protein is heterologously expressed [97, 98]. Humanization of glucose transport or mitochondrial VDAC proteins in yeast could be extremely useful to elucidate specific aspects of human hexokinases regulation and function in a human-like context.

A potential crosstalk between *Hs*PFKM and hexokinase was also revealed by comparing single gene and full pathway transplantation. *Hs*PFKM displayed a 2.5-fold higher *in vitro* activity in a strain expressing *Hs*HK2 as sole hexokinase as compared to a strain expressing *Hs*HK4, while protein abundance was identical. In the *Hs*Gly-HK4 strain the low *in vitro* activity of *Hs*PFKM activity does not match the predicted *in vivo* activity based on the observed fluxes. This discrepancy between *in vitro* and *in vivo* was even stronger in evolved isolates of *Hs*Gly-HK4, in which *Hs*PFKM was systematically mutated. Conversely, no mutations were found in *Hs*PFKM in the evolved *Hs*Gly-HK2 strains. *Hs*PFKM is regulated at multiple levels (e.g. post-translational modification, binding to various cytoskeleton components, etc.) and most likely does not operate optimally in yeast [68, 99]. Notably, stabilization of *Hs*PFKM oligomerization promoted by calmodulin in human cells might be impaired in yeast considering the difference between yeast and human calmodulin (ca. 60% identity at the protein level) [100, 101]. The present results suggest the existence of a yet unknown, hexokinase-dependent mechanisms controlling *Hs*PFKM.

Both ALE and overexpression identified hexokinase and *Hs*PGAM2 as critical enzymes for glycolytic flux and growth rate improvement of the fully humanized strains. *Hs*PGAM2 lower activity and k_cat_ as compared to human myotube cells suggested that the yeast cellular environment is not favourable for this enzyme, a problem that both humanized strains solved by increasing *Hs*PGAM2 activity. The suboptimal activity of the hexokinases was similarly solved during evolution. However, for *Hs*PGAM2 and *Hs*HK2, as well as *Hs*PFKM, which decreased in activity during evolution, the changes in *in vitro* activity in evolved strains could not be fully explained by the changes in protein abundance and did not result from mutations in non-coding or coding regions of the corresponding genes. Modulation of enzyme activity through interactions with the cellular environment or direct posttranslational covalent modifications are most likely responsible for this discrepancy between protein level and *in vitro* enzyme activity. The regulation of several human glycolytic proteins occurs via interaction with the cytoskeleton, as mentioned above for *Hs*PFKM and calmodulin. In mammalian cells, phosphoinositide signalling via PI3-kinase regulates aldolase activity by actin remodelling. The systematic mutation in the evolved humanized strains of *STT4*, encoding a PI4-kinase involved in cellular signalling for many cellular processes, including actin organization in yeast, suggests that ALDOA activity might also be modulated in yeast by binding to actin and altered by phosphoinositide-mediated signalling. Overall optimization of human glycolysis in yeast seems to be largely exerted by posttranslational mechanisms, and ALE is a powerful strategy to identify the mechanisms causing suboptimal functionality in yeast.

This first successful humanization of the skeletal muscle glycolysis in yeast offers new possibilities to explore human glycolysis. Since many complex interactions with various organelles and signalling pathways that are present in human cells will be absent in yeast, such model strains can be applied to study the pathway in a ‘clean’ background. Transplantation to the yeast context enables to dissect metabolic from signalling-related mechanisms in the control and regulation of glucose metabolism, mechanisms often debated in the field of diabetes and muscle insulin resistance. As an example, whether glycolytic enzymes themselves could be inhibited by intermediates of lipid metabolism in the muscle and consequently impact enzyme activity and glycolytic fluxes remains an open question to be tested [102, 103]. Beyond muscle tissue, the glycolysis swapping concept can be extended to any glycolytic configuration. Complete pathway transplantation can in the future be used to generate translational microbial models to study fundamental aspects of evolutionary conservation between species and tissues, and to unravel mechanisms of related diseases.

## Material and methods

### Strains, media and laboratory evolution

All strains used in this study are derived from a CEN.PK background [104] and are listed in table S7. Yeast strains were propagated on YP medium containing 10g L^-1^ Bacto Yeast extract, 20 g L^-1^ Bacto Peptone or synthetic medium containing 5 g L^-1^ (NH_4_)_2_SO_4_, 3 g L^-1^ KH_2_PO_4_, 0.5 g L^-1^ MgSO_4_·7·H_2_O, and 1 mL L^-1^ of a trace elements and vitamin solution [105]. Media were supplemented with 20 g L^-1^ glucose or galactose or 2% (v/v) ethanol. For the physiological characterization of the individual hexokinase complementation strains (Fig. 4A) and to test the aldolase moonlighting function (Fig. 4D and S10B), (NH_4_)_2_SO_4_ was replaced with 6.6 g L^-1^ K_2_SO_4_ and 2.3 g L^-1^ urea to reduce acidification of the medium. Urea was filter sterilized and added after heat sterilization of the medium at 121°C. When indicated, 125 mg L^-1^ histidine was added. For solid media 2% (w/v) agar was added to the medium prior to heat sterilization. The pH of SM was adjusted to pH 6 by addition of 2 M KOH. For selection, YP medium was supplemented with 200 mg L^-1^ G418 (KanMX) or 100 mg L^−1^ nourseothricin (Clonat). For removal of the native yeast glycolysis cassette from the *sga1* locus the SM glucose (SMG) medium was supplemented with 2.3 g L^-1^ fluoracetamide to counter select for the *AmdS* marker present in the cassette [106]. For plasmid propagation chemically competent *Escherichia coli* XL1-Blue (Agilent Technologies, Santa Clara, CA) cells were used which were grown in lysogeny broth (LB) supplemented with 100 mg L^-1^ ampicillin, 25 mg L^-1^ chloramphenicol or 50 mg L^-1^ kanamycin when required [107, 108]. Yeast and *E. coli* strains were stored at -80 °C after addition of 30% (v/v) glycerol to an overnight grown culture.

For all growth experiments in shake flask, 100 mL medium in a 500 mL shake flask was used except for the shake flask growth study with individual complementation strain for which 20 mL in a 100 mL volume shake flaks was used. Strains were incubated with constant shaking at 200 rpm and at 30°C unless stated otherwise. Strains were inoculated from glycerol stocks in YPD and grown overnight. This culture was used to inoculate the pre-culture (SMG) from which the exponentially growing cells were transferred to new shake flasks to start a growth study.

Growth studies in microtiter plate were performed at 30 °C and 250 rpm using a Growth Profiler 960 (EnzyScreen BV, Heemstede, The Netherlands). Strains from glycerol freezer stocks were inoculated and grown overnight in 10 mL YPD or YPGal medium in a 50 mL volume shake flask. This culture was used to inoculate a preculture in a 24-wells plate with a 1 mL working volume (EnzyScreen, type CR1424f) or a shakeflask with 15 mL of the medium of interest, which was grown until mid/late-exponential growth. From this culture the growth study was started in a 96-wells microtiter plate (EnzyScreen, type CR1496dl), with final working volumes of 250 µL and starting OD_660_ of 0.1-0.2. Microtiter plates were closed with a sandwich cover (EnzyScreen, type CR1296). Images of cultures were made at 30 min intervals. Green-values for each well were corrected for position in the plate using measurements of a culture of OD_660_ 5.05 of CEN.PK113-7D. Corrected green values were converted to OD-values based on 15-point calibrations, fitted with the following equation: OD-equivalent = a×GV(t) + b×GV(t)^c^ - d in which GV(t) is the corrected green-value measured in a well at time point ‘t’. This resulted in curves with a = 0.0843, b = 5.35×10^-8^, c = 4.40 and d = 0.42 for the data in Fig. 1, 5 and S2 and a = 0.077, b = 1.66×10^-7^, c = 3.62 and d = 1.61 for the data in Fig. 4 and S10. Growth rates were calculated in a time frame where the calculated OD was between 2 and 10 in which OD doubled at least twice.

Adaptive laboratory evolution of IMX1814 and IMX1844 was performed in SMG at 30°C in 100 mL volume shake flasks with a working volume of 20 mL. Initially every 48 hours 200 μL of the culture was transferred to a new shake flask with fresh medium, after 22 transfers (approximately 170 generations) this was done every 24h. For both strains three evolution lines were run in parallel. At the end of the experiment single colony isolates were obtained by restreaking three times on YPD plates (Table S7D).

### Molecular techniques, gene synthesis and Golden Gate plasmid construction

PCR amplification for cloning purposes was performed with Phusion High-Fidelity DNA polymerase (Thermo Fisher Scientific, Waltham, MA) according to the manufacturers recommendations except that the primer concentration was lowered to 0.2 μM. PCR products for cloning and Sanger sequencing were purified using the Zymoclean Gel DNA Recovery kit (Zymo Research, Irvine, CA) or the GeneJET PCR Purification kit (Thermo Fisher Scientific). Sanger sequencing was performed at Baseclear BV (Baseclear, Leiden, The Netherlands) and Macrogen (Macrogen Europe, Amsterdam, The Netherlands). Diagnostic PCR to confirm correct assembly, integration of the constructs and sequence verification by Sanger sequencing was done with DreamTaq mastermix (Thermo Fisher Scientific) according to the manufacturers recommendations. To obtain template DNA, cells of single colonies were suspended in 0.02 M NaOH, boiled for 5 min and spun down to use the supernatant. All primers used in this study are listed in Table S8. Primers for cloning purposes were ordered PAGE purified, the others desalted. To obtain gRNA and repair fragments the designed forward and reverse primers were incubated at 95°C for 5 min to obtain a double stranded piece of DNA. PCR products were separated in gels containing 1% agarose (Sigma) in Tris-acetate buffer (TAE). Genomic DNA from CEN.PK113-7D was extracted using the YeaStar™ Genomic DNA kit (Zymo Research Corporation, Irvine, CA, USA). Cloning of promoters, genes and terminators was done using Golden Gate assembly. Per reaction volume of 10 μl, 1 μl T4 buffer (Thermo Fisher Scientific), 0.5 μl T7 DNA ligase (NEB New England Biolabs, Ipswich, MA) and 0.5 μl BsaI (Eco31I) (Thermo Fisher Scientific) or BsmBI (NEB) was used and DNA parts were added in equimolar amounts of 20 fmol as previously described [109]. First a plasmid backbone was constructed from parts of the yeast toolkit [109] using a kanamycin marker, *URA3* marker, bacterial origin of replication, 3’and 5’ *ura3* integration flanks and a *GFP* marker resulting in pGGKd002 (Table S9C). In a second assembly, the *GFP* gene in this plasmid was replaced by a transcriptional unit containing a *S. cerevisiae* promoter and terminator and a human glycolytic gene. The sequences of the human glycolytic genes were obtained from the Uniprot data base, codon optimized for *S. cerevisiae* and ordered from GeneArt Gene Synthesis (Thermo Fisher Scientific). Genes were synthetized flanked with BsaI restriction sites to use them directly in Golden Gate assembly (Table S9A). The *PKL* gene which is a shorter splicing variant of *PKR* was obtained by amplifying it from the *PKR* plasmid pGGKp024 using primers containing BsaI restriction site flanks (Table S8G)*. S. cerevisiae* promoters and terminators were PCR amplified from genomic DNA using primers flanked with BsaI and BsmBI restriction sites (Table S8A) [110]. The resulting PCR product was directly used for Golden Gate assembly. For long term storage of the fragments, the promoters and terminators were cloned into the pUD565 entry vector using BsmBI Golden Gate cloning resulting in the plasmids pGGKp025-048 listed in Table S9B. For the *HXK2* and *TEF2* promoters and *HXK2* and *ENO2* terminators already existing plasmids were used (Table S9B). For the construction of pUDE750 which was used as PCR template for the amplification of the *HsHK4* fragment used in IMX1814, first a dropout vector (pGGKd003) was constructed from the yeast toolkit parts pYTK002, 47, 67, 74, 82 and 84 (Table S9C). In this backbone, *ScHXK2p, HK4* and *ScHXK2t* were assembled as described above (Table S9E). Plasmid isolation was done with the GenElute™ Plasmid Miniprep Kit (Sigma-Aldrich, St. Louis, MO). Yeast transformations were performed according to the lithium acetate method [111].

### Construction of individual gene complementation strains

To enable CRISPR/Cas9 mediated gene editing, *Cas9* and the *NatNT1* marker were integrated in the *SGA1* locus of the minimal glycolysis strain IMX370 by homologous recombination, resulting in strain IMX1076 [27]. *Cas9* was PCR amplified from p414-*TEF1p-Cas9-CYC1t* and *NatNT1* from pUG-*natNT1* (Tables S8G and S9E) and 750 ng of both fragments were, after gel purification, used for transformation.

For the individual gene complementation study, 400 ng of the constructed plasmids containing the human gene transcriptional units (Table S9D) were linearized by digestion with NotI (FastDigest, Thermo Fisher Scientific) according to the manufacturer’s protocol for 30 min and subsequently the digestion mix was directly transformed to IMX1076. The linearized plasmids were integrated by homologous recombination in the disrupted *ura3-52* locus of strain IMX1076 and the transformants were plated on SMG. After confirmation of correct integration by PCR (Table S8B), in a second transformation the orthologous yeast gene (or genes, in case of *PFK1* and *PFK2*) was removed using CRISPR/Cas9 according to the protocol of Mans *et al.* [112]. Since only the yeast gene and not the human ortholog should be targeted, the gRNAs were designed manually (Table S8D). For deletion of *FBA1*, *GPM1*, and *PFK1* and *PFK2,* the plasmids containing the gRNA were preassembled as previously described [112] using Gibson assembly and a PCR amplified pROS13 backbone containing the KanMX marker (Tables S8D and S9F). For *HXK2* deletion, the double stranded gRNA and a PCR amplified pMEL13 backbone were assembled using Gibson assembly (Table S8D). The constructed plasmids were verified by PCR. The rest of the gRNA plasmids for yeast gene deletion were assembled *in vivo* in yeast and were not stored as individual plasmid afterwards. For the *in vivo* assembly approach the strains were co-transformed with 100 ng of the PCR amplified backbone of pMEL13 (Table S8D and S9F), 300 ng of the double stranded gRNA of interest (Table S8D) and 1 μg repair fragment to repair the double stranded break (Table S8E). For pre-assembled plasmids, strains were co-transformed with 0.6-1 μg of plasmid (Table S9F) and 1 μg repair fragment (Table S8E). Transformants were plated on YPD + G418 and for the *HsHK1*-*HK3* strains on YPGal + G418. Successful gene deletion was confirmed with diagnostic PCR (Table S8B, Fig. S18). gRNA plasmids were afterwards removed by several restreaks on non-selective medium. To test if the complementation was successful, the strains were tested for growth in SMG. For *HsHK2,* three complementation strains were made. IMX1690 (*pScPDC1*-*HsHK2)* and IMX1873 (*pScHXK2-HsHK2*) which were grown on glucose medium and contain a mutation in *HsHK2* and IMX2419 (*pScPDC1*-*HsHK2)* which was never exposed to glucose and does not contain mutations. Similarly for *HsHK1,* complementation strain IMX1689 was not grown on glucose, after growth on glucose mutations occurred and strains IMS1137, IMS1140 and IMS1143 were stocked. *HsHK4* was also expressed both with the *ScHXK2* and *ScPDC1* promoter, resulting in IMX1874 and IMX1334 respectively (Table S7A,B). An overview of the workflow is provided in Fig. S1. To test for the occurrence of mutations, the human gene transcriptional units were PCR amplified using the primers listed in Table S8 and sent for Sanger sequencing.

### Full human glycolysis strain construction

For the construction of the strains containing a full human glycolysis, the transcriptional units of the *HsHK2, HsHK4, HsGPI, HsPFKM, HsALDOA, HsTPI1, HsGAPDH, HsPGAM2, HsENO3,* and *HsPKM1* gene were PCR amplified from the same plasmids as were used for the individual gene complementation using primers with flanks containing synthetic homologous recombination (SHR) sequences (Table S8C and S9D). For the *HsHK2* and *HsHK4* gene for which pUDE750 and pUDI207 were used as template, which contain the Sc*HXK2* promoter and terminator. An overview of the promoters used for the human gene expression is provided in Table S2. The yeast *PDC1* and *ADH1* genes were amplified with their corresponding promoter and terminator regions from genomic DNA from CEN.PK113-7D (Table S7E). The fragments were gel purified and the fragments were assembled in the *CAN1* locus of strain IMX589 by *in vivo* assembly. 160 fmol per fragment and 1 μg of the pMEL13 plasmid targeting *CAN1* was used. Transformation mix was plated on YPD + G418 and correct assembly was checked by PCR and resulted in strain IMX1658. In a second transformation, the cassette in the *SGA1* locus containing the native *S. cerevisiae* glycolytic genes and the *AmdS* marker was removed. To this end IMX1658 was transformed with 1 μg of the gRNA plasmid pUDE342 (Table S9F) and 2 μg repair fragment (counter select oligo) (Table S8E) and plated on SMG medium with fluoracetamide to counter select for the *AmdS* marker. From the resulting strain the pUDE342 plasmid was removed and it was stored as IMX1668. To replace the *HsHK4* gene with *HsHK2,* the *HsHK2* gene was PCR amplified from pGGKp002 using primers flanked with sequences homologous to the *ScHXK2* promoter and terminator to allow for recombination (Table S8G). IMX1668 was co-transformed with this fragment and pUDR387 containing the gRNA targeting the *HsHK4* gene and the cells were plated on YPD + G418 (Table S9F). After confirmation of correct integration by PCR and plasmid removal, the strain was stored as IMX1785. The pUDR387 gRNA plasmid was constructed with Gibson Assembly from a pMEL13 backbone and double stranded *HsHK4* gRNA fragment (Table S8D). To make the constructed yeast strains prototrophic, the *ScURA3* marker was PCR amplified from CEN.PK113-7D genomic DNA using primers with flanks homologous to the *TDH1* region (Table S8G) and integrated in the *tdh1* locus of IMX1785 and IMX1668, by transforming the strains with 500 ng of the fragment and plating on SMG. This resulted in IMX1844 and IMX1814 respectively. These strains were verified by whole genome sequencing (Table S3) and the ploidy was verified (Fig. S19). IMX2418 (*Hs*Gly *HK2* strain without mutation in Hs*HK2*) was constructed by transforming IMX1814 with pUDR387 and the *HsHK2* fragment amplified as described above. The cells were plated on YPGal + G418 and later restreaked on YPGal plates to remove the plasmid. For the overexpression of *HsALDOA, HsPGAM2* and *HsHK2/HsHK4* resulting in IMX2005 and IMX2006, the expression cassettes were PCR amplified from pUDI141, pUDI150, pUDI134 and pUDI136 respectively using primer sets 12446/12650, 12467/14542 and 14540/14541 (Table S8C, S9D). IMX1844 and IMX1814 were transformed with 160 fmol per fragment and 1 μg of the plasmid pUDR376 containing a gRNA targeting the X2 locus [113] and plated on SMG-acetamide plates. To obtain the reference strain IMX1821 which contains a yeast glycolysis cassette integrated in *CAN1,* the pUDE342 plasmid was removed from the previously described strain IMX605 [28] and the *URA3* fragment was integrated in *tdh1* in the manner as described above. An overview of strain construction is provided in Fig. S20.

### *STT4* reverse engineering

The single nucleotide polymorphisms (SNPs) which were found in the *STT4* gene of evolved strains IMS0990 and IMS0992 resulting in amino acid changes G1766R and F1775I respectively, were introduced in the *STT4* genes of the non-evolved strains IMX1814, IMX1844 and IMX1822 using CRISPR/Cas9 editing [112] (Table S7D). Two gRNA plasmids pUDR666 and pUDR667 were constructed using Gibson Assembly of a backbone amplified from pMEL13 (Table S8D, S9F) and a gRNA fragment consisting of oligo 16748+16749 and 16755+16756 respectively (Table S8D). For introduction of the G1766R mutation, strains were transformed with 500 ng of pUDR666 and 1 µg of repair fragment (oligo 16750+16751) and for introduction of F1775I with 500 ng of pUDR667 and 1 µg of repair fragment (oligo 16757+16758) (Table S8E, S9F). Strains were plated on YPD + G418 and introduction of the mutation was verified by Sanger sequencing. The control strain IMX1822 containing the native yeast minimal glycolysis in the *SGA1* locus originates from strain IMX589 [28]. From this strain the *AmdS* marker was removed by transforming the strain with 1 µg repair fragment (oligo 11590+11591) and 300 ng of a gRNA fragment (oligo 11588+11589, Table S8D) targeting *AmdS* and 100 ng of backbone amplified from pMEL10 resulting in a *in vivo* assembled gRNA plasmid. After removal of the plasmid by restreaking on non-selective medium, this strain, IMX1769, was made prototrophic by integrating *ScURA3* in *tdh1* (Table S8G, S7E), resulting in IMX1822.

### Visualization of hexokinase mutants and mathematical modelling

Sequencing of the *HsHK1* and *HsHK2* carrying strains showed the presence of mutations in all strains after growth on glucose. All found mutations were mapped unto the protein sequence and visualized on the structural model with PDB code 1HKB [114] for *Hs*HK1 and 2NZT [33] for *Hs*HK2 using the PyMOL Molecular Graphics System, version 1.8.6 (Schrödinger LLC).

Native human hexokinase complementation strains were simulated with the use of a previously published computational model of yeast glycolysis [40]. The SBML version of the model was downloaded from *jjj.bio.vu.nl/models/vanheerden1* and imported into COPASI (software version 4.23) [115]. All concentrations in the model are expressed as mM and time in minutes. Equilibrium constants (Keq’s) were obtained from Equilibrator [116] at pH 6.8 and ionic strength 360 mM [117]. The V_max_’s from the MG strain from Kuijpers et al., 2016 [28] were initially incorporated to create a control strain. The forward V_max_’s from GAPDH and PGI were calculated from the measured reverse V_max_’s and model Keq’s and Km’s according to the Haldane relationship. For the complementation strains, the kinetic equation of hexokinase was first adapted to include competitive terms from G6P and ADP inhibition and trehalose 6-phosphate inhibition of the human enzymes was disregarded based on our kinetic results. Mammal kinetic parameters were obtained from [44]. *Hs*HK1 and *Hs*HK2 glycolysis models were subsequently obtained by incorporating the hexokinase V_max_ measured from strains IMX1689 and IMX2419, respectively. Steady-state fluxes were calculated with the integration of ordinary differential equations. Flux control coefficients (FCC) and response coefficients (R) were calculated in COPASI under Metabolic Control Analysis and Sensitivities according to equations 1 and 2 below, respectively. J_ss_ represents a steady-state flux, for which the glucose uptake flux was used. V_max i_ represents the V_max_ while P_i_ stands for a kinetic parameter of an enzyme ‘i’ in the pathway. Summation theorem was verified for FCC calculations [118]. A complete overview of model construction and assumptions can be found in Appendix 1.

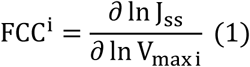

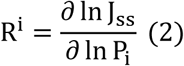

### Construction of *Δhxk1Δhxk2* strain IMX165 and control strain IMX2015

The *Δhxk1Δhxk2* strain IMX165 which was used as control in the invertase assay was constructed in three steps. The *HXK1* and *HXK2* deletion cassettes were PCR amplified from pUG73 and pUG6 respectively using the primers listed in Table S8F. First, *HXK1* was removed from CEN.PK102-12A by transformation with the *HXK1* deletion cassette containing the *Kluyveromyces lactis LEU2* marker flanked with loxP sites and *HXK1* recombination flanks resulting in strain IMX075. To remove the *LEU2* marker from this strain, it was transformed with the plasmid pSH47 containing the galactose inducible Cre recombinase [119]. Transformants were plated on SMG with histidine and were transferred to YPGal for Cre recombinase induction to remove *LEU2,* resulting in strain IMS0336. Subsequently, this strain was transformed with the *HXK2* deletion cassette containing the *KanMX* marker flanked with LoxP sites and *HXK2* recombination flanks, resulting in IMX165. IMX2015 was constructed as control strain for the characterization of the human hexokinase complementation strains. In this strain Sc*HXK2* is expressed with the p*PDC1* promoter instead of the native *HXK2* promoter. *pPDC1* was PCR amplified from genomic DNA from CEN.PK113-7D with primer 14670 and 14671 containing *HXK2* recombination flanks. 500 ng of this fragment was transformed to IMX1076 together with 800 ng of pUDE327 containing a gRNA targeting the *HXK2* promoter (table S9F).

### Illumina whole genome sequencing

Genomic DNA for sequencing was isolated with the the Qiagen 100/G kit according to the manufacturer’s description (Qiagen, Hilden, Germany) and library preparation and sequencing was done as described previously using Illumina Miseq sequencing (Illumina, San Diego, CA) [110]. A list of mutations is provided in Table S3. For the mutation found in the *SBE2* gene which is involved in bud growth, it is unlikely to have an effect since it has a functionally redundant paralog *SBE22* [120]. No abnormalities were observed under the microscope. The sequencing data generated in this project are accessible at NCBI under bioproject PRJNA717746.

### Quantitative aerobic batch cultivations

Quantitative characterization of strain IMX1821, IMX1814 and IMX1844 was done in 2 L bioreactors with a working volume of 1.4 L (Applikon, Schiedam, The Netherlands). The cultivation was done in synthetic medium supplemented with 20 g L^-1^ glucose, 1.4 mL of a vitamin solution [105] and 1.4 mL of 20% (v/v) Antifoam emulsion C (Sigma, St. Louise, USA). During the fermentation 0.5 mL extra antifoam was added when necessary. The salt and antifoam solution were autoclaved separately at 121°C and the glucose solution at 110°C for 20 min. During the fermentation the temperature was kept constant at 30°C and the pH at 5 by automatic addition of 2 M KOH. The stirring speed was set at 800 rpm. The medium was flushed with 700 mL min^-1^ of air (Linde, Gas Benelux, The Netherlands).

For preparation of the inoculum, freezer stocks were inoculated in 100 mL YPD and grown overnight. From this culture the pre-culture was inoculated in 100 mL SMG which was incubated till mid-exponential growth phase. This culture was used to inoculate the inoculum flasks which were incubated till OD 4.5. The cells were centrifuged for 10 min at 3000g and the pellet was suspended in 100 mL demineralized water and added to the fermenter to start the fermentation with an OD of 0.25-0.4.

Biomass dry weight determination was done as previously described [105] by filtering 10 mL of culture on a filter with pore-size 0.45 mm (Whatman/GE Healthcare Life Sciences, Little Chalfont, United Kingdom) in technical duplicate. For extracellular metabolite analysis 1 mL of culture was centrifuged for 3 min at 20000g and the supernatant was analysed using high performance liquid chromatography (HPLC) using an Aminex HPX-87H ion-exchange column operated at 60°C with 5 mM H_2_SO_4_ as the mobile phase with a flow rate of 0.6 mL min^-1^ (Agilent, Santa Clara). The OD_660_ was measured with a Jenway 7200 spectrophotometer (Jenway, Staffordshire, UK) at 660 nm. Per strain at least two independent fermentations were performed. The carbon balances for all reactors closed within 5%.

### Sample preparation and enzymatic assays for comparison of yeast and humanized yeast samples

Yeast samples were prepared as previously described [121], from exponentially growing cultures (62 mg dry weight per sample) from bioreactor and for testing of allosteric effectors and for comparison of the evolved strains from shake flask. Sonication was used for cell-free extract preparation except for the hexokinase measurements (Fig. 2, S4 and S5) where fast-prep was used. All determinations were performed at 30°C and 340 nm (εNAD(P)H at 340 nm/6.33 mM^-1^).

In most cases glycolytic *V_max_* enzyme activities were determined in 1 mL reaction volume (in 2 mL cuvettes), using a Hitachi model 100-60 spectrophotometer, using previously described assays [52], except for phosphofructokinase activity which was determined according to Cruz *et al.*[122]. To increase throughput, the specific activities of the evolved strains and the glucose, ATP and ADP dependency of the hexokinase complementation strains (Fig S4 C-E) were assayed using a TECAN infinite M200 Pro. (Tecan, Männedorf, Switzerland) microtiter plate reader. Samples were prepared manually in microtiter plates (transparent flat-bottom Costar plates; 96 wells) using a reaction volume of 300 µl per well. The assays were the same as for the cuvette-based assays. For determination of glucose-6-phosphate inhibition of hexokinase, cell extracts were prepared in Tris-HCl buffer (50 mM, pH 7.5) to limit phosphate concentrations, for these measurements an alternative enzyme assay coupled by pyruvate kinase and lactate dehydrogenase was used based on [123], buffer and metabolite concentrations were kept the same as the yeast hexokinase assay. The reported data are based on at least two independent biological replicate samples, with at least two analytic replicates per sample per assay, including two different cell free extract concentrations. The protein concentration was determined using the Lowry method with bovine serum albumin as a standard [124]. Enzyme activities are expressed as μmol substrate converted (mg protein)^-1^ h^-1^.

To calculate the degree of saturation of glycolytic enzymes, the specific activity in μmol^-1^.mg_protein_ ^-1^. h was converted into mmol.gDW^-1^.h^-1^ considering that soluble proteins represent 30% of cell dry weight. This value represents the maximal enzyme flux capacity. The *in vivo* flux in the glycolytic reactions were approximated from the glucose specific uptake rate (q_glu_). Reactions in the top of glycolysis (hexokinase to triosephosphate isomerase) were assumed to equal the q_glu_, while reactions in the bottom of glycolysis (glyceraldhyde-3P dehydrogenase to pyruvate kinase) were calculated as the q_glu_ times two. The degree of saturation was calculated as follows:

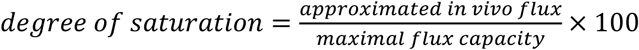

### Invertase enzyme assay

The invertase assay was performed on whole cells previously described [125]. Exponentially growing cells in SMG were washed with sterile dH_2_O, transferred to shake flasks (at OD 3) with 100 mL fresh SMG or SME+0.075% glucose and incubated for 2h at 30°C and shaking at 200 rpm. Afterwards the dry weight of the cultures was determined and the cells were washed in 50mM sodium acetate buffer wit 50mM NaF to block the metabolism and were then suspended till a concentration of 2.5-7.5 mg dry weight per mL. 4 mL of this cell suspension were added to a dedicated vessel thermostated at 30°C, and kept under constant aeration by flushing with air (Linde, Gas Benelux, The Netherlands) and stirring with a magnetic stirrer. The reaction was started by addition of 1 mL 1M sucrose and 1 mL reaction mix was taken at 0, 1, 2, 3 and 5 minutes, directly filtered using 13 mm diameter 0.22 μm pore size nylon syringe filters to remove cells and put on ice. Afterwards the glucose concentration resulting from sucrose hydrolysis by invertase was determined using a D-Glucose assay kit (Megazyme, Bray, Ireland). The glucose production rate was calculated in μMol·min^-1^·g dry weight^-1^.

### Staining of vacuoles

Yeast strains were stained with the red fluorescent dye FM4-64 (excitation/emission, 515/640 nm) (Thermo Fisher Scientific). Exponentially growing cells were incubated at an OD of 0.5-1 in YPD with 2 µM FM4-64 in the dark for 30 minutes at 30°C. Afterwards cells were spun down, washed and incubated for 2-3 h in 5 mL YPD. For analysis, cells were spun down and suspended in SMG medium. Yeast cells and vacuoles were visualized with an Imager-Z1 microscope equipped with an AxioCam MR camera, an EC Plan-Neofluar 100x/1.3 oil Ph3 M27 objective, and the filter set BP 535/25, FT 580, and LP 590 (Carl-Zeiss, Oberkochen, Germany).

### Ploidy determination by flow cytometry

Samples of culture broth (equivalent to circa 10^7^ cells) were taken from mid-exponential shake-flask cultures on YPD and centrifuged (5 min, 4700g). The pellet was washed once with demineralized water, and centrifuged again (5 min, 4700g) and suspended in 800 μL 70% ethanol while vortexing. After addition of another 800 μL 70% ethanol, fixed cells were stored at 4°C until further staining and analysis. Staining of cells with SYTOX^®^ Green Nucleic Acid Stain (Invitrogen S7020) was performed as described [126]. Samples were analysed on a BD Accuri C6 flow cytometer equipped with a 488 nm laser (BD Biosciences, Breda, The Netherlands). The fluorescence intensity (DNA content) was represented using FlowJo (v. 10.6.1, FlowJo, LLC, Ashland, OR, USA), (Fig. S19).

### Transition experiment

For testing transitioning between carbon sources, strains were grown overnight in SMGal medium till mid-exponential phase. These cultures were used to plate single cells on SMG and SMGal plates (96 cells per plate) using a BD FACSAriaII (Franklin Lakes, NJ). After 5 days the percentage of cells which was growing was determined.

### Whole cell lysate proteomics of humanized yeast strains

#### Sampling, cell lysis, protein extraction, in-solution proteolytic digestion and TMT labelling

For proteomics analysis, yeast strains grown to exponential phase in SMG shake flasks were inoculated to fresh shake flasks in biological triplicates. During mid-exponential phase samples of 8 ml were taken, centrifuged for 10 min. at 5000 g at 4°C and the pellet was stored at -80°C. Approx. 50mg of cell pellets (wet weight) were lysed using beads beating in 1% SDS, 100mM TEAB, including protease inhibitor and phosphate inhibitor. The lysed cells were centrifuged and the supernatant was transferred to a new tube. The proteins were reduced with dithiothreitol and alkylated using iodoacetamide (IAA), where the protein content was precipitated and washed using ice cold acetone. The protein pellets were dissolved in 100mM ammonium bicarbonate buffer, and subjected to overnight digestion using proteomics grade Trypsin at 37°C and under gentle shaking using an Eppendorf incubator. The peptides were desalted using solid-phase extraction on a Waters Oasis HLB 96-well µElution plate according to the manufacturers protocol, SpeedVac dried and stored at -20°C until analysed. One aliquot of each sample was dissolved in 3% acetonitrile in H_2_O, containing 0.1% formic acid and subjected to nLC-Orbitrap-MS analysis for digestion quality control purpose. One aliquot of each sample was further dissolved in 100mM TEAB for further labelling using a TMT 10plex labelling kit (Thermo scientific, catalog number: 90110). The peptide content of the samples were estimated using a NanoDrop photospectrometer, and the samples were diluted to achieve an approx. equal concentration. The TMT labelling agents were dissolved by adding 40µL anhydrous acetonitrile, and 5µL of each label was added to the individual samples. The samples were incubated at 25°C under gentle shaking, for 75 minutes. Then the reaction was stopped by adding diluted hydroxylamine solution, and incubated at 25°C under gentle shaking. Equal amounts of sample were combined in a LoBind Eppendorf tube. After dilution with aqueous buffer, solid phase extraction was once more performed according to the manufacturers protocol. The samples were SpeedVac dried, and stored at -20°C until analysed. Before analysis, the samples were solubilised in 3% acetonitrile and 0.01% TFA and subjected to nLC-Orbitrap-MS analysis.

#### Whole cell lysate shotgun proteomics

An aliquot corresponding to approx. 500ng of every TMT 10plex peptide mixture (SET1, SET2 and SET3, where 3 strains were mixed and compared within one SET, and each strain within one SET was present as triplicate) was analysed by duplicate analysis employing one-dimensional shotgun proteomics. The sets were chosen to be able to directly compare the evolved strains with their parental strain and to compare the humanized strains with the *Sc*Gly strain, SET1 contained strains IMX1814, IMS0990 and IMS0991, SET2 IMX1844, IMS0987 and IMS0989, and SET3 IMX1814, IMX1844 and IMX1821. Briefly, TMT labelled peptides were analysed using a nano-liquid-chromatography system consisting of an EASY nano LC 1200, equipped with an Acclaim PepMap RSLC RP C18 separation column (50 µm x 150 mm, 2µm and 100 Å), online coupled to a QE plus Orbitrap mass spectrometer (Thermo Scientific, Germany). The flow rate was maintained at 350nL/min over a linear gradient from 5% to 25% solvent B over 178 minutes, and finally to 55% B over 60 minutes. Solvent A consisted of H_2_O containing 0.1% formic acid, and solvent B consisted of 80% acetonitrile in H_2_O, plus 0.1% formic acid. The Orbitrap was operated in data-dependent acquisition mode acquiring peptide signals form 385-1450 m/z at 70K resolution, 75ms max IT, and an AGC target of 3e6, where the top 10 signals were isolated at a window of 1.6m/z and fragmented at a NCE of 32. The peptide fragments were measured at 35K resolution, using an AGC target of 1e5 and allowing a max IT of 100ms.

#### MS raw data processing and determination of protein expression levels

For protein identification, raw data were processed using PEAKS Studio 10.0 (Bioinformatics Solutions Inc.) allowing 20ppm parent and 0.01Da fragment mass error tolerance, TMT10plex and Carbamidomethylation as fixed, and methionine oxidation and N/Q deamidation as variable modifications. Data were matched against an in-house established yeast protein sequence database, including the GPM crap contaminant database (https://www.thegpm.org/crap/) and a decoy fusion for determining false discovery rates. Peptide spectrum matches were filtered against 1 % false discovery rate (FDR) and protein identifications were accepted as being significant when having 2 unique peptides matches minimum. Quantitative analysis of the global proteome changes between the individual yeast strains was performed using the PEAKS-Q software package (Bioinformatics Solutions Inc.), considering a quantification mass tolerance of 10ppm, a FDR threshold of 1%, using auto normalisation and ANOVA as the significance method. Significance (-10log(p)) vs fold change volcano plots were created using the scatter function in Matlab2019b (Fig. S9 and S15). TIC normalized signal intensity was calculated by dividing the signal intensity by the total intensity of each sample (Fig. S8 and S13). Mass spectrometric raw data have been deposited to the ProteomeXchange Consortium (http://proteomecentral.proteomexchange.org) via the PRIDE repository with the dataset identifier PXD025349.

### Comparison human and humanized yeast glycolytic enzymes

#### Human cell culture and harvest

Human myoblasts were obtained from orbicularis oculi muscle biopsies, as previously described [127]. Briefly, subclone V49 expressed Pax7, MyoD and Myogenin and was used for the assays here described. Cells were maintained in high glucose Dulbecco’s Modified Eagle’s Medium (DMEM, Sigma-Aldrich/Merck) in the presence of L-glutamine, 20% fetal bovine serum (FBS, Life Technologies Gibco/Merck) and 1% penicillin/streptomycin (p/s, Sigma-Aldrich/Merck). For differentiation, cells were seeded on 10 cm dishes covered with polydimethylsiloxane (PDMS) gradients at 5,000 cells/cm^2^ and after reaching confluence, medium was changed to DMEM in the presence of 2% FBS, 1% p/s, 1% Insulin-Transferrin-Selenium (Life Technologies Gibco/Merck) and 1% dexamethasone (Sigma-Aldrich/Merck). The presence of PDMS gradients allows cells to grow aligned, which in turn improves myotube maturity and functionality. Cells were harvested after 5 days in differentiation medium. In short, cells were washed twice with ice-cold Dulbecco’s Phosphate Buffered Saline (DPBS, Gibco) and scraped in DPBS in the presence of Complete Protease Inhibitor Cocktail (Merck, 11836145001, 1:25 v/v after resuspension according to manufacturer’s guidelines). Cells were frozen at -80 °C.

#### Cell-free extract preparation and *V_max_* enzyme assays

Human cells stored at -80°C were thawed, centrifuged at 20000 g for 10 minutes at 4 °C and the pellet was discarded to obtain cell-free extracts.

Yeast samples (IMX1844) were harvested as previously described [121] from exponentially growing cultures (62 mg dry weight per sample) from bioreactor. Cell-free extract preparation for yeast cells was done using YeastBuster^TM^ Protein Extraction Reagent supplemented with 1% of 100x THP solution according to the description (Novagen, San Diego, CA, USA). To a pellet with a wet weight of 0.3 g, 3.5 mL YeastBuster and 35 μl THP solution was added. The pellet was suspended and incubated for 20 min at room temperature. Afterwards the cell debris was removed by centrifugation at 20000 g for 15 min at 4 °C and the supernatant was used for the assays.

Prior to experimentation, YeastBuster^TM^ Protein Extraction Reagent with 1% THP (Novagen) was added to the human cell samples and DPBS supplemented with protease inhibitor was added to the yeast samples (both as 50% of final volume). This strategy was taken in order to equalize the buffer composition of yeast and human culture samples to perform enzyme kinetics assays and proteomics.

*V_max_* assays for comparison of yeast and human cell extracts were carried out with freshly prepared extracts via NAD(P)H-linked assays at 37 °C in a Synergy H4 plate reader (BioTek™). The reported *V_max_* values represent total capacity of all isoenzymes in the cell at saturating concentrations of all substrates and expressed per extracted cell protein. Four different dilutions of extract were used to check for linearity. Unless otherwise stated, at least 2 dilutions were proportional to each other and these were used for further calculation. All enzymes were expressed as µmoles of substrate converted per minute per mg of extracted protein. Protein determination was carried out with the Bicinchoninic Acid kit (BCA™ Protein Assay kit, Pierce) with BSA (2 mg/ml stock solution of bovine serum albumin, Pierce) as standard.

Based on the cytosolic concentrations described in literature, we have designed an assay medium that was as close as possible to the *in vivo* situation, whilst at the same time experimentally feasible. The standardized *in vivo*-like assay medium contained 150 mM potassium[128–131], 5 mM phosphate [128, 132], 15 mM sodium [128, 133], 155 mM chloride [134, 135], 0.5 mM calcium, 0.5 mM free magnesium [128, 136, 137] and 0.5-10.5 mM sulfate. For the addition of magnesium, it was taken into account that ATP and ADP bind magnesium with a high affinity. The amount of magnesium added equalled the concentration of either ATP or ADP plus 0.5 mM, such that the free magnesium concentration was 0.5 mM. Since the sulfate salt of magnesium was used the sulfate concentration in the final assay medium varied in a range between 0.5 and 10.5 mM. The assay medium was buffered at a pH of 7.0 [138–143] by using a final concentration of 100 mM Tris-HCl (pH 7.0). To end up with the above concentrations, an assay mixture containing 100 mM Tris-HCl (pH 7.0), 15 mM NaCl, 0.5 mM CaCl_2_, 140 mM KCl, and 0.5-10.5 mM MgSO_4_ was prepared.

In addition to the assay medium, the concentrations of the coupling enzymes, allosteric activators and substrates for each enzyme were as follows:

**Hexokinase (HK; EC2.7.1.1)** – 1.2 mM NADP^+^, 10 mM Glucose, 1.8 U/mL glucose-6-phosphate dehydrogenase (EC1.1.1.49), and 10 mM ATP as start reagent.

**Phosphoglucose isomerase (GPI; EC5.3.1.9)** – 0.4 mM NADP^+^, 1.8 U/mL glucose-6-phosphate dehydrogenase (EC1.1.1.49), and 2 mM fructose 6-phosphate as start reagent.

**Phosphofructokinase (PFK; EC2.7.1.11)** – 0.15 mM NADH, 1 mM ATP, 0.5 U/mL aldolase (EC4.1.2.13), 0.6 U/mL glycerol-3P-dehydrogenase (EC1.1.1.8), 1.8 U/mL triosephosphate isomerase (EC5.3.1.1), 65 µM fructose 2,6-bisphosphate as activator (synthesized as previously described [144]), and 10 mM fructose 6-phosphate as start reagent.

**Aldolase (ALDO; EC4.1.2.13)** – 0.15 mM NADH, 0.6 U/mL glycerol-3P-dehydrogenase (EC1.1.1.8), 1.8 U/mL triosephosphate isomerase (EC5.3.1.1), and 2 mM fructose 1,6-bisphosphate as start reagent.

**Glyceraldehyde-3-phosphate dehydrogenase (GAPDH; EC1.2.1.12)** – 0.15 mM NADH, 1 mM ATP, 24 U/mL 3-phosphoglycerate kinase (EC2.7.2.3), and 5 mM 3-phosphoglyceric acid as start reagent.

**3-phosphoglycerate kinase (PGK; EC2.7.2.3)** – 0.15 mM NADH, 1 mM ATP, 8 U/mL glyceraldehyde-3-phosphate dehydrogenase (EC1.2.1.12), and 5 mM 3-phosphoglyceric acid as start reagent.

**Phosphoglycerate mutase (PGAM; EC5.4.2.1)** – 0.15 mM NADH, 1 mM ADP, 2.5 mM 2,3-diphospho-glyceric acid, 5 U/mL enolase (EC4.2.1.11), 50 U/mL pyruvate kinase (EC2.7.1.40), 60 U/mL L-lactate dehydrogenase (EC1.1.1.27), and 5 mM 3-phosphoglyceric acid as start reagent.

**Enolase (ENO; EC4.2.1.11)** – 0.15 mM NADH, 1 mM ADP, 50 U/ml pyruvate kinase (EC2.7.1.40), 15 U/mL L-lactate dehydrogenase (EC1.1.1.27), and 1 mM 2-phosphoglyceric acid as start reagent.

**Pyruvate kinase (PK; 2.7.1.40)** – 0.15 mM NADH, 1 mM ADP, 1 mM fructose 1,6-bisphosphate, 60 U/mL L-lactate dehydrogenase (EC1.1.1.27) and 2 mM phosphoenolpyruvate as start reagent.

#### Determination of absolute enzyme concentrations [E]

Absolute concentrations of glycolytic targets was performed by targeted proteomics [145]. Isotopically labelled peptides with ^13^C lysines and arginines were designed for human glucose metabolism and a list of peptides of interest detected in our samples can be found in Table S6.

#### Turnover number (*k_cat_*) calculations

Turnover numbers were estimated based on the equation *V_max_* = *k_cat_*·[*E*], the maximal enzyme activity was divided by the concentration of each individual peptide detected by proteomics when no other isoform was detected. In the human skeletal muscle samples, more than one isoform was detected for certain proteins. In these cases (phosphoglycerate mutase, enolase and pyruvate kinase) the sum of the concentrations of all isoforms was used to estimate the turnover number. In Fig. S17 k_cat_ values were calculated for the specific isoforms based on the ratio between turnover numbers found in the literature for the human isoforms. Protein concentrations [E] were measured as *pmol · mg protein^-1^* and V_max_’s as *µmol · min^-1^ ·mg protein^-1^*. In order to obtain k_cat_ values in *min^-1^*, the following equation was used for each enzymatic reaction in the dataset:

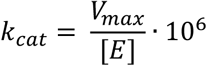

## Acknowledgements

We thank Jordi Geelhoed for his work on reverse engineering and growth characterization, Agnes Hol and Ingeborg van Lakwijk for their valuable work on the human hexokinase strains and Lycka Kamoen, Eveline Vreeburg and Daniel Solis-Escalante for their contribution to construction of reference strains. We thank Carol de Ram for processing proteomics samples. We thank Pilar de la Torre and Marcel van den Broek for whole genome sequencing and analysis, and Philip de Groot for help with bioreactor sampling.

## Funding

This project was funded by the AdLibYeast European Research Council (ERC) consolidator 648141 grant awarded to PD-L., a UMCG-GSMS PhD fellowship and the De Cock-Hadders Foundation.

## Author contributions

F.B., E.K., M.V.-L., designed and performed research and wrote the article, M.W. performed molecular biology and strain construction, M.A.H.L. performed enzyme activity and ploidy assays, K.v.E. set up *in vivo* like enzyme assays, M.d.R., J.C.W. and M.P. performed proteomics analyses, R.B., A.M.A.S. set up and performed human myotube cultures, M.C.H., P.v.R. provided material and input on the *in silico* model development, J.M.D.,B.M.B and P.D.-L designed and supervised the research and wrote the article.

All authors approved the final manuscript

## Competing interests

Authors declare no competing interests

## Supplementary material

Supplementary Figures S1 to S20

Supplementary Tables S1 to S9

Supplementary methods computational modelling: Appendix 1

## Notes

### Competing Interest Statement

The authors have declared no competing interest.

