## Supplementary Material for "A yeast with muscle doesn’t run faster: full humanization of the glycolytic pathway in *Saccharomyces cerevisiae*"

#### Figure S1 - Complementation strategy

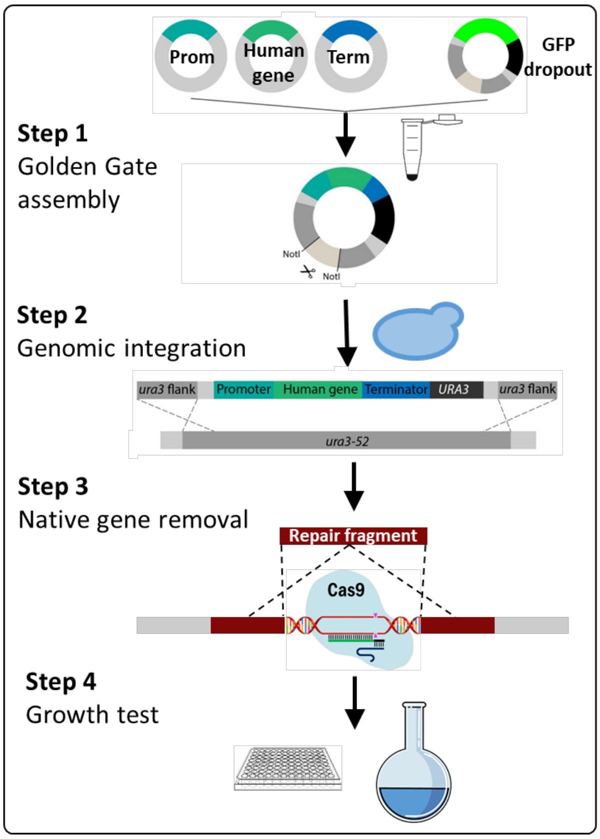


Codon-optimized human glycolytic genes were stitched to yeast promoters and terminators using Golden Gate assembly (step 1). The resulting plasmids were linearized by restriction with NotI and integrated in the *ura3-52* locus of the minimal glycolysis strain IMX1076 which was plated on SMG (Step 2). In a second transformation round, the yeast ortholog was selectively removed using CRISPR/Cas9-mediated DNA editing. Transformed cells were plated on YPD + G418, except for the *HsHK1-3* strains which were plated on YPGal + G418 (step 3). Gene integration and deletion was checked for all strains by PCR and Sanger sequencing. All strains were tested for growth on chemically defined medium with glucose as sole carbon source (SMG).

#### Figure S2 - Growth rate of complementation strains on SMG

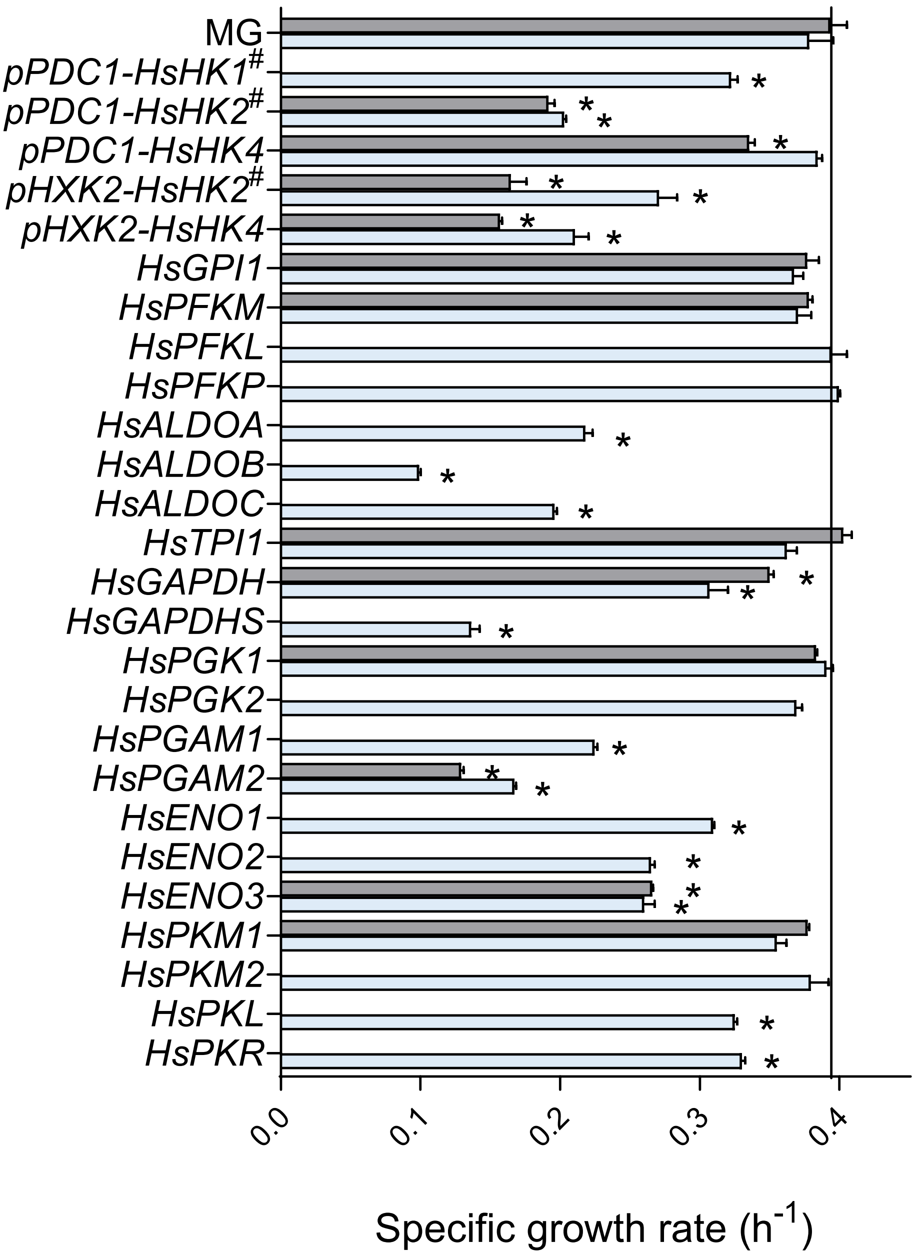


Specific growth rates of complementation strains and minimal glycolysis (MG) control strain determined in shake flask (grey) and growth profiler (light blue) using chemically defined medium with glucose as sole carbon source (SMG). # Indicates strains with mutations after growth on glucose, *pPDC1-HsHK1*: IMS1137, *pPDC1-HsHK2*: IMX1689, *pHXK2-HsHK2*: IMX1873. * indicates significant difference from control strain MG (IMX372) (P < 0.01, Student t-test, two-tailed, homoscedastic).

#### Figure S3 – Comparison of promoter strength, protein identity and growth rate in the complementation strains

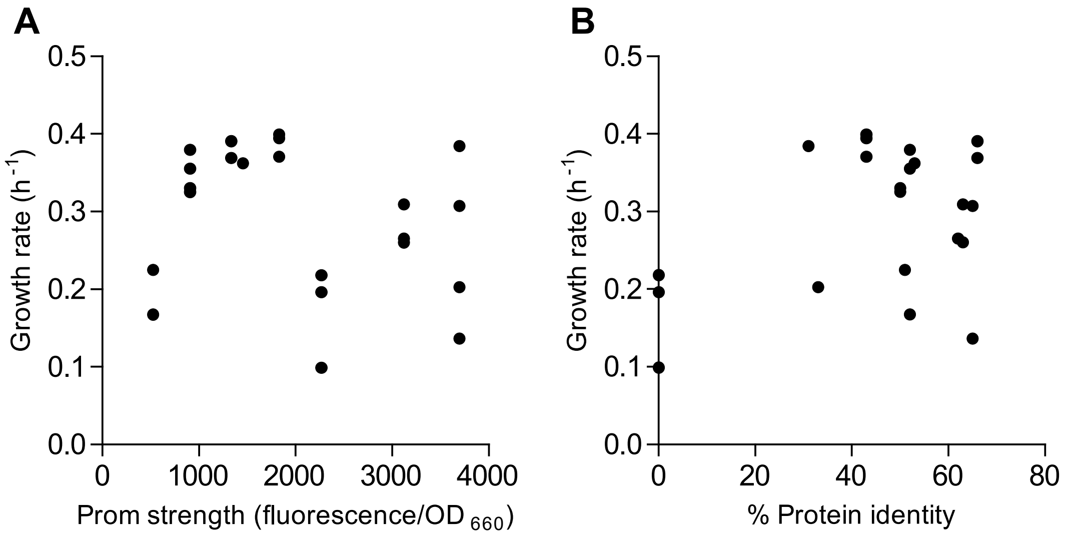


The growth rate of the complementation strains measured in the growth profiler was plotted against: **A)** the strength of the promoter used to express the human gene. Promoter strengths were obtained from Boonekamp *et al.* 2018 [1]. **B)** Percentage protein identity between the human gene and the corresponding yeast ortholog.

### Figure S4 - Human hexokinase 2 complementation strain characterization

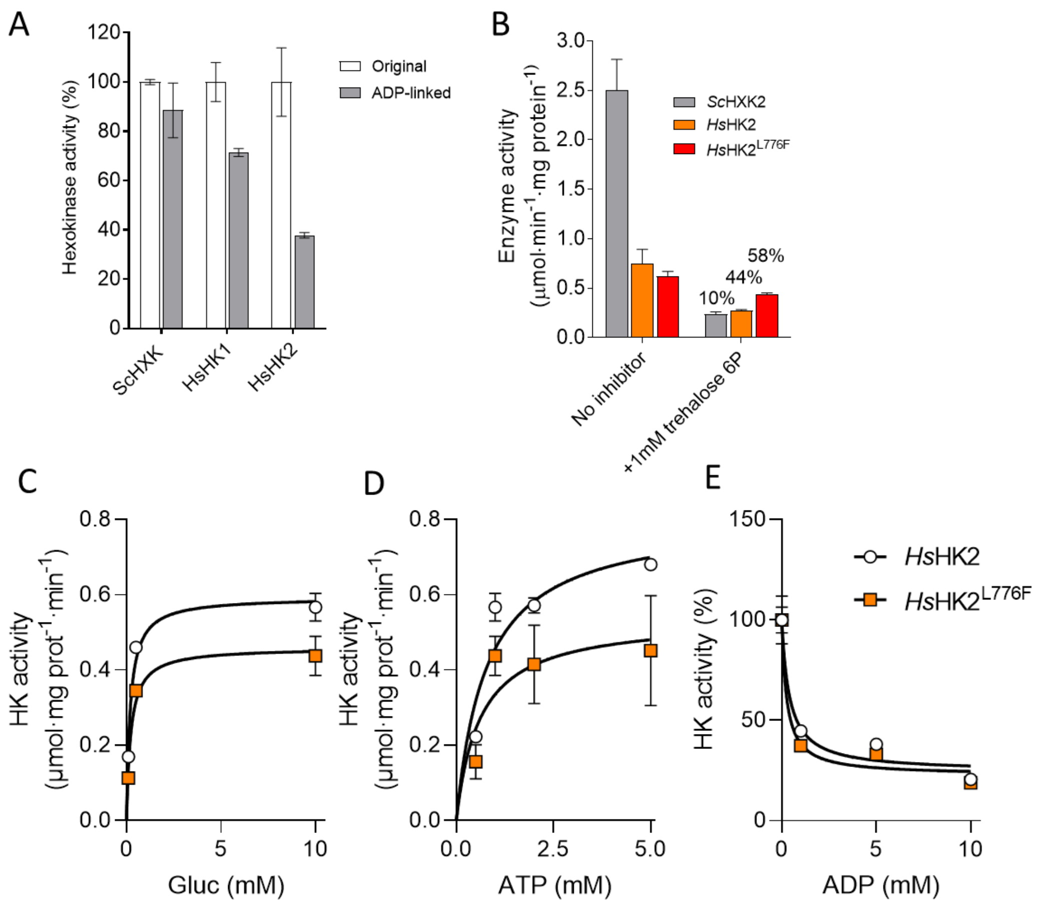


F

| Parameter | *Hs*HK2 Native | *Hs*HK2 L776F |
|---|---|---|
| $K_m$ gluc (mM) | 0.20±0.04 | 0.23±0.06 |
| $K_m$ ATP (mM) | 0.77±0.04 | 0.61±0.44 |
| $IC_{50}$ ADP (mM) | 0.41±0.20 | 0.26±0.16 |

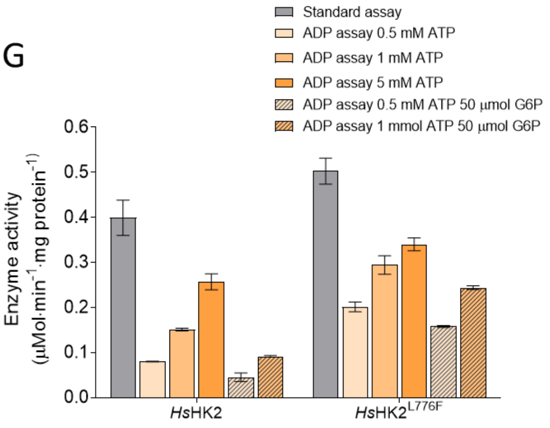


*In vitro* biochemical characterization was performed for the non-mutated *Hs*HK2 and *Hs*HK2$^{L776F}$ variant (strains IMX2419 and IMX1690 respectively). Strains were grown in shake flask on SM-Galactose medium to prevent any selection for mutations. **A)** As the standard hexokinase assay was based on G6P detection (original assay), a different assay based on ADP detection (ADP-linked assay) was implemented to quantify hexokinase sensitivity to G6P. The ADP-linked assay systematically detected a lower hexokinase activity, possibly due to G6P accumulation during the assay. Hexokinase activity expressed as % of the activity measured with the original assay for the same strain. For all other measurements (panel B-F) the original G6PDH-linked assay was used. **B)** Inhibition by trehalose-6-phosphate of the yeast *Sc*Hxk2 (strain IMX2015) and the native and mutated variants of *Hs*HK2 enzymes. 1 mM trehalose-6P resulted in 90% inhibition of the yeast *Sc*Hxk2, while it only marginally affected the human *Hs*HK2. As concentrations of Trehalose-6P above 1 mM are not often encountered in yeast cells, [2-4], trehalose-6P inhibition was probably not responsible for the lack of activity of the native *Hs*HK2 in yeast cells. Inhibition of *Hs*HK2 and *Hs*HK2$^{L776F}$ was not significantly different (t-test, two-tailed, homoscedastic, $p>0.05$). **C-E)** Response to varying concentrations of glucose, ATP and ADP of the two *Hs*HK2 variants. These measurements were performed in a 96-wells reader. **F)** Calculated kinetic parameters of *Hs*HK2 and *Hs*HK2$^{L776F}$ based on the data shown in previous panels. **G)** Hexokinase activity at various concentrations of ATP and G6P concentrations in the ADP-linked assay. Increasing ATP concentrations led to an increase in measured activity in the ADP-linked assay consistent with an effect of the production of competitive inhibitor G6P which is not removed in this assay.

#### Figure S5 – Human hexokinase 2 characterization in the fully humanized strains

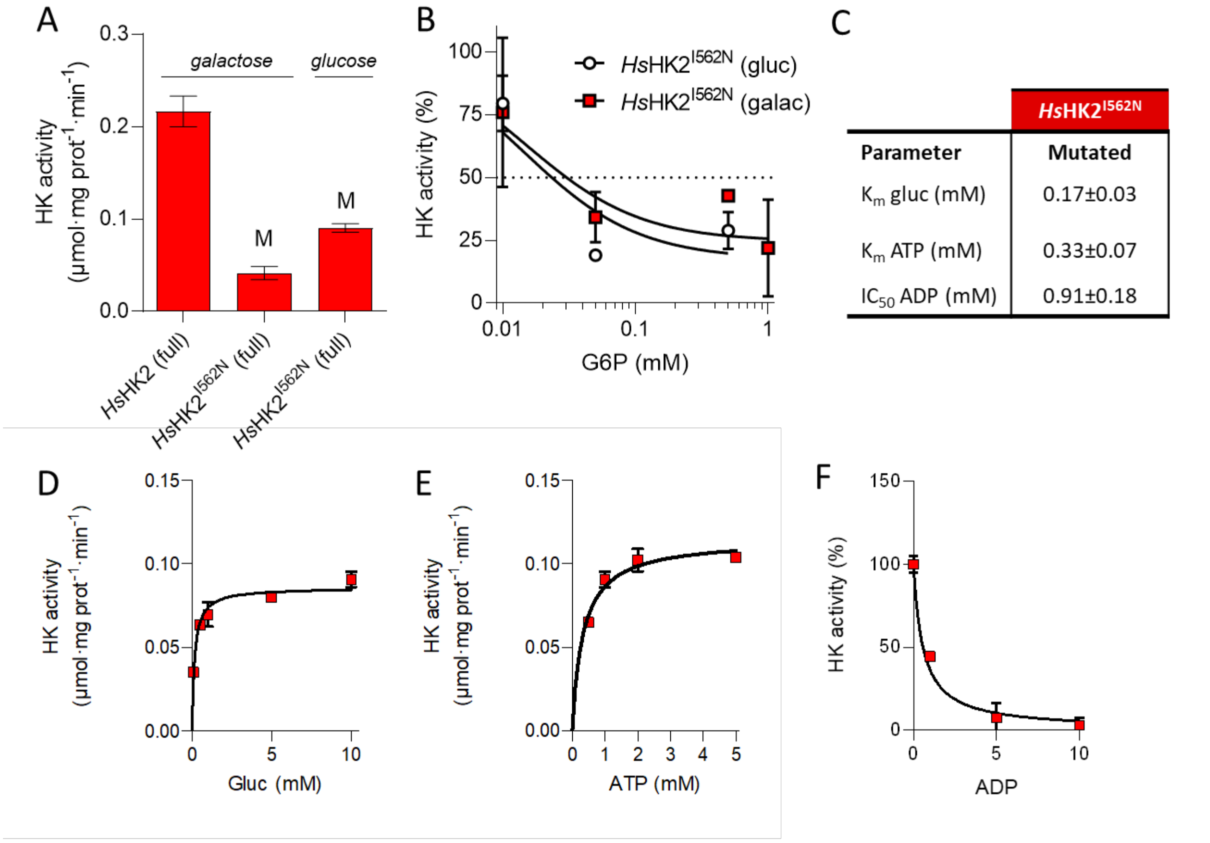


*In vitro* characterization of hexokinase in cell free extracts from the fully humanized *Hs*Gly-HK2 strain IMX1844 (*Hs*HK2$^{I562N}$) and fully humanized control strain IMX2496 (*Hs*Gly-HK2 without mutation in HK2). In both strains, *Hs*HK2 is expressed from the *ScHXK2* promotor, which is repressed in galactose cultures, hence both glucose and galactose grown cultures were compared for strain IMX1844. **A)** comparison of hexokinase activity measured in the IMX2496 and IMX1844 strains shows a strong decrease in activity in the mutant strain. **B)** The sensitivity of the mutated hexokinase to glucose-6-phosphate is independent of the carbon source used to grow the fully humanized strain. **C)** Calculated kinetic parameters of hexokinase *Hs*HK2$^{I562N}$ in strain IMX1844 from the data shown in panels D-F. For these measurements strain IMX1844 was grown on SM glucose (non-repressive condition). All values are in the same order of magnitude as the values measured for the *Hs*HK2 complementation strains shown in Fig. S4 and hence do not explain the lower measured activity. **D-F)** Hexokinase activity of glucose-grown IMX1844 (*Hs*Gly-HK2) at various concentrations of glucose, ATP and ADP.

#### Figure S6 – Sensitivity of human and yeast pyruvate kinase to fructose-1,6bisP

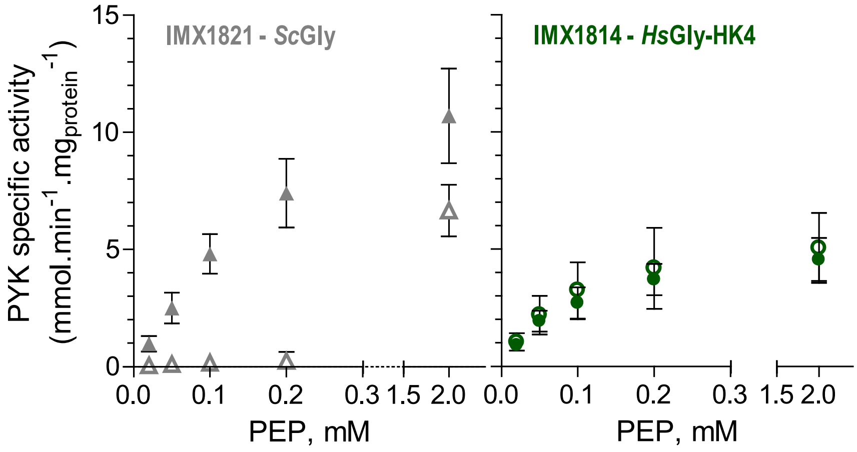


Yeast (left) and human (right) specific pyruvate kinase activity *in vitro* assayed with fructose-1,6bisP (closed symbols) and without fructose-1,6bisP (open symbols) in IMX1821 (*Sc*gly) and IMX1814 (*Hs*gly) with different concentrations of the substrate phosphoenolpyruvate (PEP). Symbols and error bars represent the average and SEM of three biological replicates.

#### Figure S7 – Robustness to transitioning between carbon sources

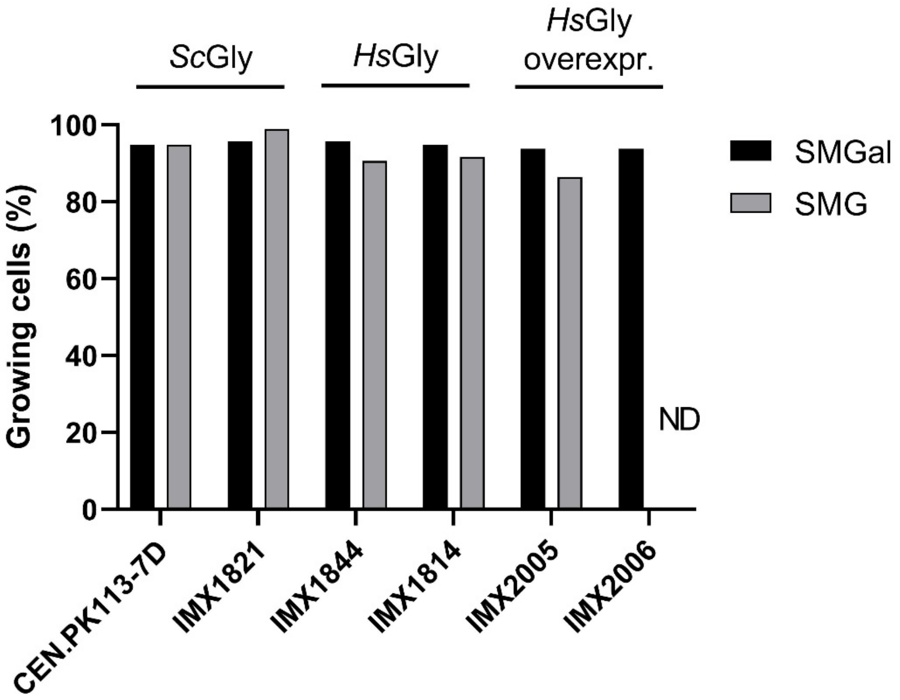


The control *Sc*Gly and humanized glycolysis strains were cultivated in liquid synthetic medium with galactose as sole carbon source (SMGal), from these cultures, single cells were plated on plates with fresh SM medium with glucose (SMG) or galactose as sole carbon source. Per condition 96 single cells were plated. Five days after plating, the colonies growing on SMGal and SMG were counted. These data originate from a single experiment. ND; not determined. Strains with humanized glycolysis displayed no impairment in transitioning between respiratory and fermentative carbon sources.

#### Figure S8 – Specific activity and relative levels of glycolytic enzymes in fully humanized yeast strains

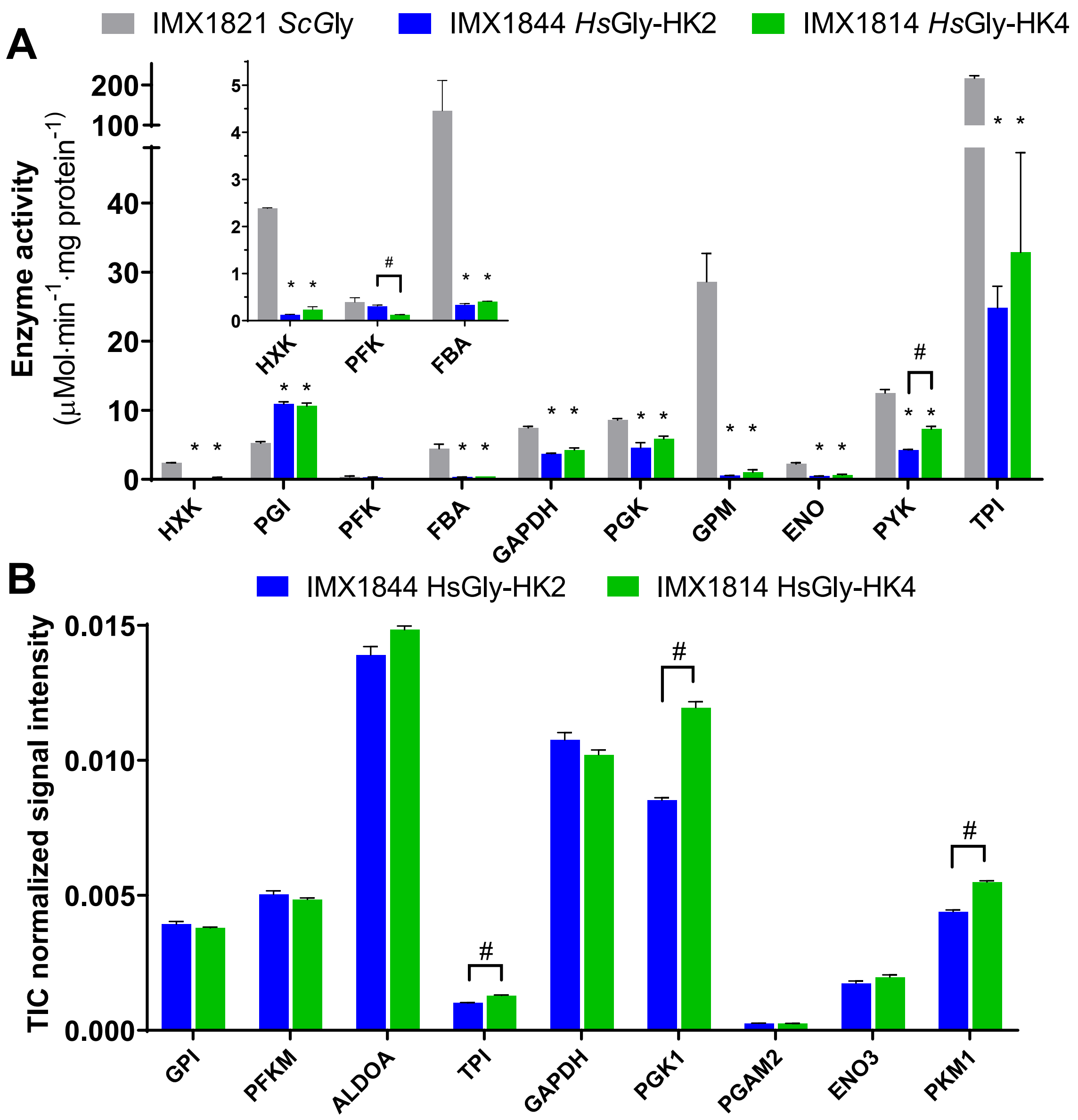


**A)** Specific enzyme activities measured *in vitro* with cell free extracts from batch cultures in bioreactors. Error bars represent the SEM from two biological replicates. * indicate values that are significantly different from the control strain IMX1821 with *S. cerevisiae* glycolysis, # indicate significant difference between the two humanized strains (Student t-test, two-tailed, homoscedastic, P<0.05). **B)** Relative enzyme abundance of the human glycolytic enzymes (normalized to the total signal intensity) in the *Hs*Gly-HK2 (IMX1844) and *Hs*Gly-HK4 (IMX1814) strains. Means with SEM are shown for biological triplicates from one of the duplicate injections. # indicates significant differences between IMX1814 and IMX1844 (p-value <0.05 student t-test, two tailed, homoscedastic).

#### Figure S9 - Global proteome response to humanization of the glycolytic pathway

A

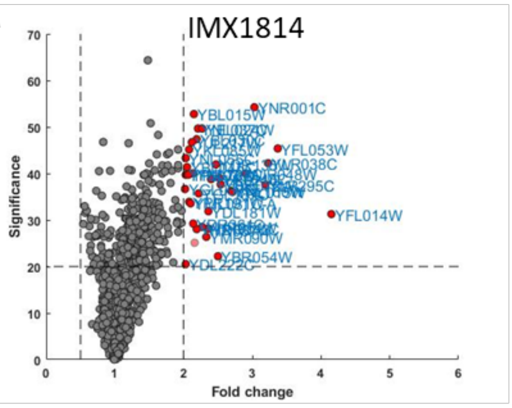


B

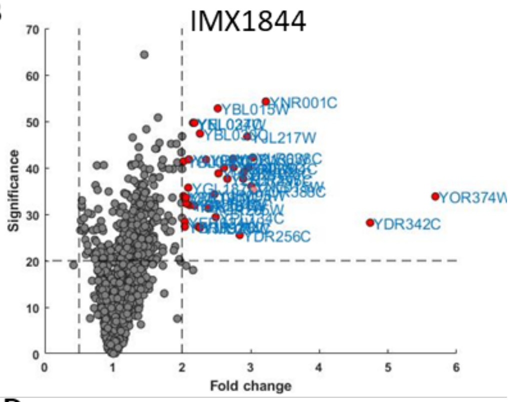


C

| Systematic name | Common name | Fold change IMX1814 vs IMX1821 | P value | TIC normalized abundance IMX1821 |
|---|---|---|---|---|
| YFL014W | Hsp12 | 4.15 | 7.4E-04 | 1.5E-04 |
| YFL053W | Dak2 | 3.37 | 2.9E-05 | 1.5E-05 |
| YLR038C | Cox12 | 3.23 | 6.0E-05 | 2.8E-04 |
| YLR295C | Atp14 | 3.19 | 1.7E-04 | 1.8E-04 |
| YNR001C | Cit2/Cit1 | 3.03 | 3.8E-06 | 2.4E-03 |
| YJR048W | Cyc7/Cyc1 | 2.9 | 1.0E-04 | 1.4E-03 |
| YNL015W | Pbi2 | 2.71 | 2.5E-04 | 2.7E-04 |
| YNL160W | Ygp1/Sps100 | 2.69 | 2.4E-04 | 1.2E-04 |
| YBR072W | Hsp26 | 2.54 | 1.7E-04 | 2.6E-04 |
| YKL148C | Sdh1 | 2.52 | 1.2E-04 | 3.6E-04 |
| YBR054W | Mrh1/Yro2 | 2.5 | 6.0E-03 | 4.5E-05 |
| YOR136W | Idh2 | 2.47 | 6.4E-05 | 8.9E-04 |
| YHL021C | Aim17 | 2.4 | 1.3E-04 | 2.0E-05 |
| YDR342C | hypothetical protein | 2.36 | 1.5E-03 | 2.2E-05 |
| YDL181W | Inh1/Stf1 | 2.36 | 6.5E-04 | 1.5E-04 |
| YMR090W | hypothetical protein | 2.33 | 2.3E-03 | 3.6E-05 |
| YEL024W | Rip1 | 2.27 | 1.1E-05 | 5.0E-04 |
| YHR051W | Cox6 | 2.27 | 1.4E-03 | 7.4E-04 |
| YGL187C | Cox4 | 2.22 | 2.7E-04 | 5.1E-04 |
| YNL037C | Idh1 | 2.21 | 1.1E-05 | 1.2E-03 |
| YBL030C | Aac3/Pet9 | 2.19 | 1.8E-05 | 2.4E-03 |
| YHR087W | Rtc3 | 2.19 | 1.6E-03 | 3.8E-05 |
|  | Ddr48 | 2.16 | 3.1E-03 | 1.0E-05 |
| YBL015W | Ach1 | 2.15 | 5.3E-06 | 1.6E-03 |
| YNL104C | Leu4/Leu9 | 2.14 | 9.9E-05 | 9.2E-04 |
| YDR361C | Bcp1 | 2.14 | 1.2E-03 | 1.0E-05 |
| YJL217W | hypothetical protein | 2.12 | 2.1E-05 | 4.2E-04 |
| YML081C-A | Atp18 | 2.1 | 4.4E-04 | 8.6E-05 |
| YPR191W | Qcr2 | 2.08 | 4.1E-04 | 1.7E-03 |
| YKL085W | Mdh1 | 2.08 | 3.1E-05 | 5.2E-04 |
| YPR020W | Atp20 | 2.07 | 1.1E-04 | 1.8E-04 |
| YBL045C | Cor1 | 2.05 | 7.3E-05 | 1.4E-03 |
| YNL055C | Por2/Por1 | 2.03 | 4.6E-05 | 2.0E-03 |
| YPL078C | Atp4 | 2.03 | 1.1E-04 | 6.8E-04 |
| YDL222C | Fmp45 | 2.03 | 8.9E-03 | 6.0E-06 |
| YGL037C | Pnc1 | 2.02 | 2.2E-04 | 1.2E-04 |

D

| Systematic name | Common name | Fold change IMX1844 vs IMX1821 | P value | TIC normalized abundance IMX1821 |
|---|---|---|---|---|
| YOR374W | Ald4 | 5.69 | 4.1E-04 | 1.5E-03 |
| YDR342C | hypothetical protein | 4.74 | 1.5E-03 | 2.2E-05 |
| YNR001C | Cit2/Cit1 | 3.22 | 3.8E-06 | 2.4E-03 |
| YOR388C | Fdh1 | 3.04 | 3.1E-04 | 2.0E-04 |
|  | hypothetical protein | 3.04 | 3.1E-04 | 2.0E-04 |
| YLR038C | Cox12 | 3.03 | 6.0E-05 | 2.8E-04 |
| YNL015W | Pbi2 | 3.01 | 2.5E-04 | 2.7E-04 |
| YNL104C | Leu4/Leu9 | 2.97 | 9.9E-05 | 9.2E-04 |
| YJL217W | hypothetical protein | 2.95 | 2.1E-05 | 4.2E-04 |
| YKL148C | Sdh1 | 2.9 | 1.2E-04 | 3.6E-04 |
| YLR295C | Atp14 | 2.89 | 1.7E-04 | 1.8E-04 |
| YDR256C | Cta1 | 2.84 | 2.8E-03 | 4.3E-05 |
| YER065C | Icl1 | 2.75 | 9.9E-05 | 1.3E-04 |
| YOR136W | Idh2 | 2.74 | 6.4E-05 | 8.9E-04 |
| YLR174W | Idp3/Idp2 | 2.66 | 1.7E-04 | 4.2E-04 |
| YJR048W | Cyc7/Cyc1 | 2.61 | 1.0E-04 | 1.4E-03 |
| YHL021C | Aim17 | 2.53 | 1.3E-04 | 2.0E-05 |
| YBL015W | Ach1 | 2.52 | 5.3E-06 | 1.6E-03 |
| YPL134C | Odc2/Odc1 | 2.49 | 1.2E-03 | 5.7E-06 |
| YJR095W | Sfc1 | 2.47 | 3.7E-04 | 1.2E-04 |
| YOR285W | Rdl1 | 2.38 | 7.4E-04 | 3.5E-04 |
| YPR002W | Pdh1 | 2.35 | 6.7E-05 | 8.2E-04 |
| YBL030C | Aac3/Pet9 | 2.26 | 1.8E-05 | 2.4E-03 |
| YML054C | Cyb2 | 2.26 | 2.0E-03 | 5.9E-05 |
| YJR148W | Bat2/Bat1 | 2.23 | 1.8E-03 | 7.3E-05 |
| YEL024W | Rip1 | 2.18 | 1.1E-05 | 5.0E-04 |
| YDL181W | Inh1/Stf1 | 2.18 | 6.5E-04 | 1.5E-04 |
| YNL037C | Idh1 | 2.16 | 1.1E-05 | 1.2E-03 |
| YGR234W | Yhb1 | 2.11 | 6.4E-04 | 4.5E-03 |
| YLL041C | Sdh2 | 2.1 | 6.6E-05 | 2.2E-04 |
| YGL187C | Cox4 | 2.09 | 2.7E-04 | 5.1E-04 |
| YER057C | Mmf1/Hmf1 | 2.06 | 5.8E-04 | 9.2E-04 |
| YML081C-A | Atp18 | 2.06 | 4.4E-04 | 8.6E-05 |
| YER024W | Yat2 | 2.04 | 1.4E-03 | 1.9E-05 |
| YBR092C | Pho3 | 2.04 | 1.8E-03 | 1.4E-04 |
| YPR191W | Qcr2 | 2.02 | 4.1E-04 | 1.7E-03 |
| YBL045C | Cor1 | 2.02 | 7.3E-05 | 1.4E-03 |

A-B) Volcano plots showing fold change and significance of the complete set of identified proteins of the humanized strains relative to the *Sc*Gly strain IMX1821. Proteins that changed more than 2-fold with a significance above 20 (corresponding to a p value below 0.01) are highlighted red. C-D) Proteins that changed >2-fold are listed with common name, fold change, p-value and normalized abundance in reference strain (*Sc*Gly, IMX1821).

#### Figure S10 – Moonlighting functions in fully humanized strains.

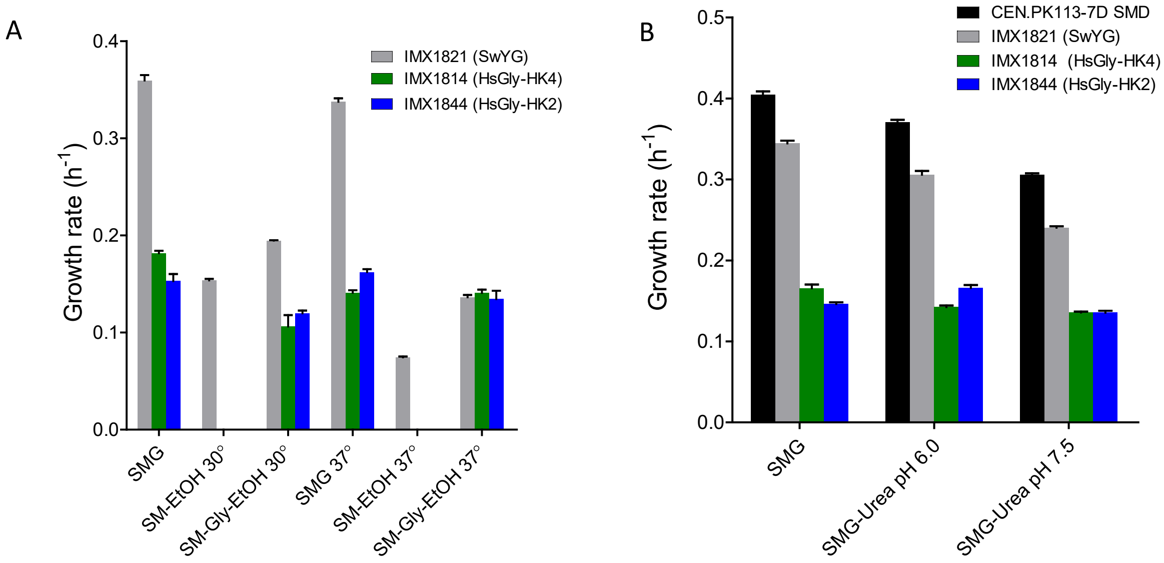


Growth rates of the fully humanized strains IMX1844 and IMX1814 were determined in conditions where moonlighting functions of enolase and aldolase are known to play a role. **A)** Growth rates at 30 and 37 °C. Strain IMX1814 and IMX1844 (*Hs*Gly-HK4 and *Hs*Gly-HK2) did not show growth in synthetic medium with only ethanol as carbon source. **B)** Growth rates at different media pH, acidification was prevented by use of urea instead of ammonium as nitrogen source. In both high temperature and high pH conditions no decrease in growth rate specific to the *Hs*Gly strains was observed. Specific growth rates were determined in 96-well plates using the growth profiler, data represents average and standard deviation of independent culture triplicates.

#### Figure S11 – Specific growth rate during evolution of IMX1844 and IMX1814

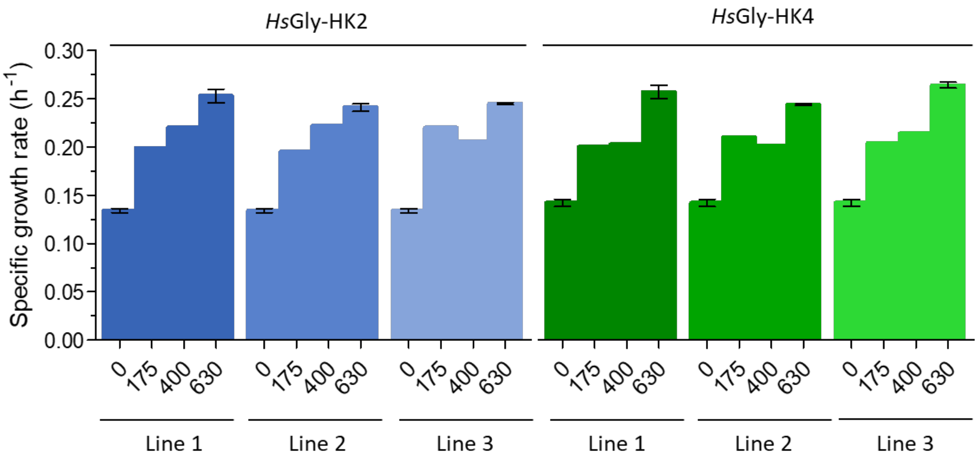


The specific growth rates were measured in shake-flask with SMG. Independent duplicates were only done for 0 and 630 generations. Measurements at 400 and 630 generations are from single colony isolates, while at 175 generations they were done with the whole evolved population.

### Figure S12 - Physiology of evolved, humanized glycolysis strains in shake flask

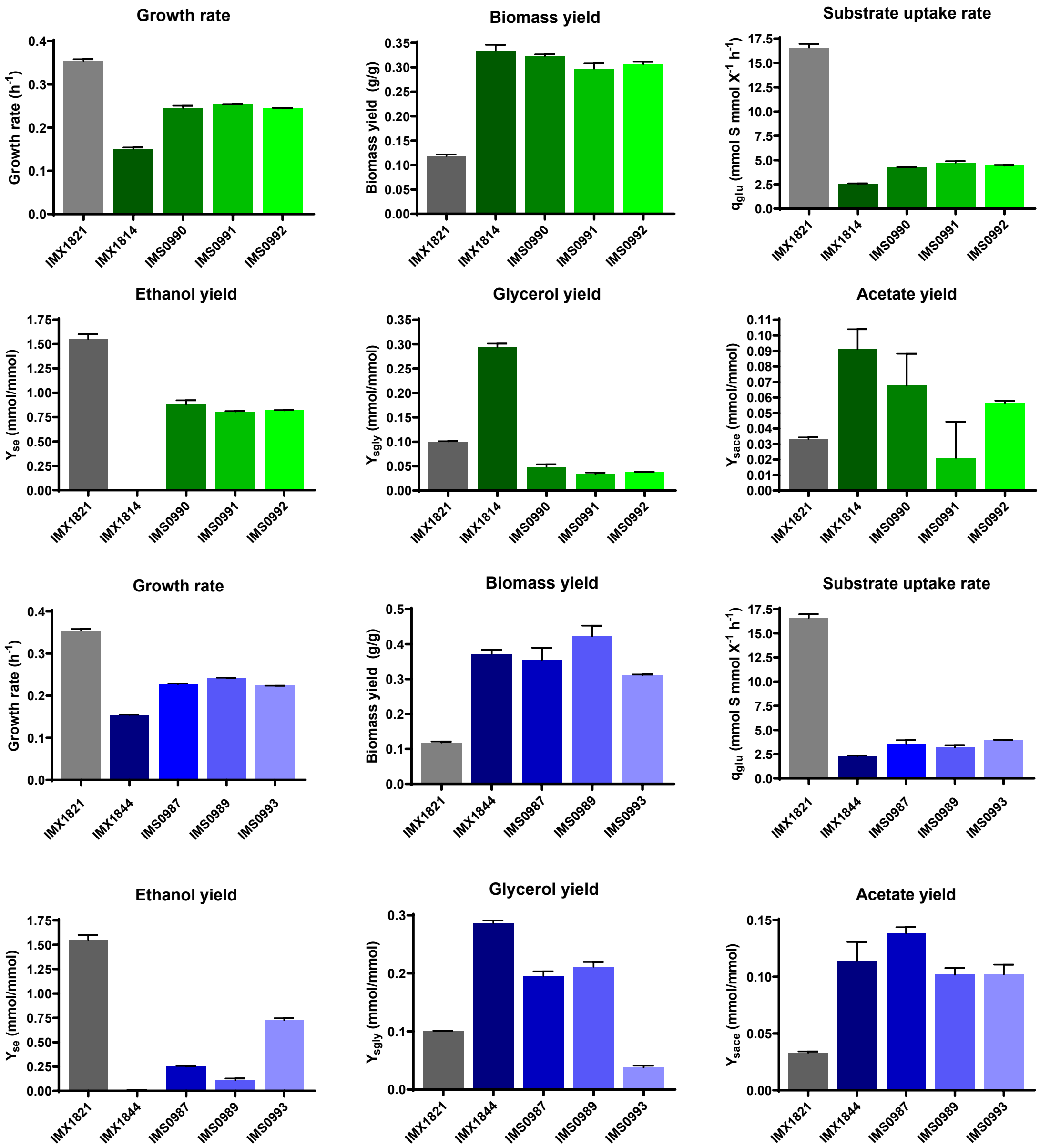


Strains were grown in SMG at 30 °C in shake flask cultures. Data represent the average and SEM of two independent culture replicates.

#### Figure S13 – Relative levels and activity of the glycolytic enzymes in evolved humanized yeast strains

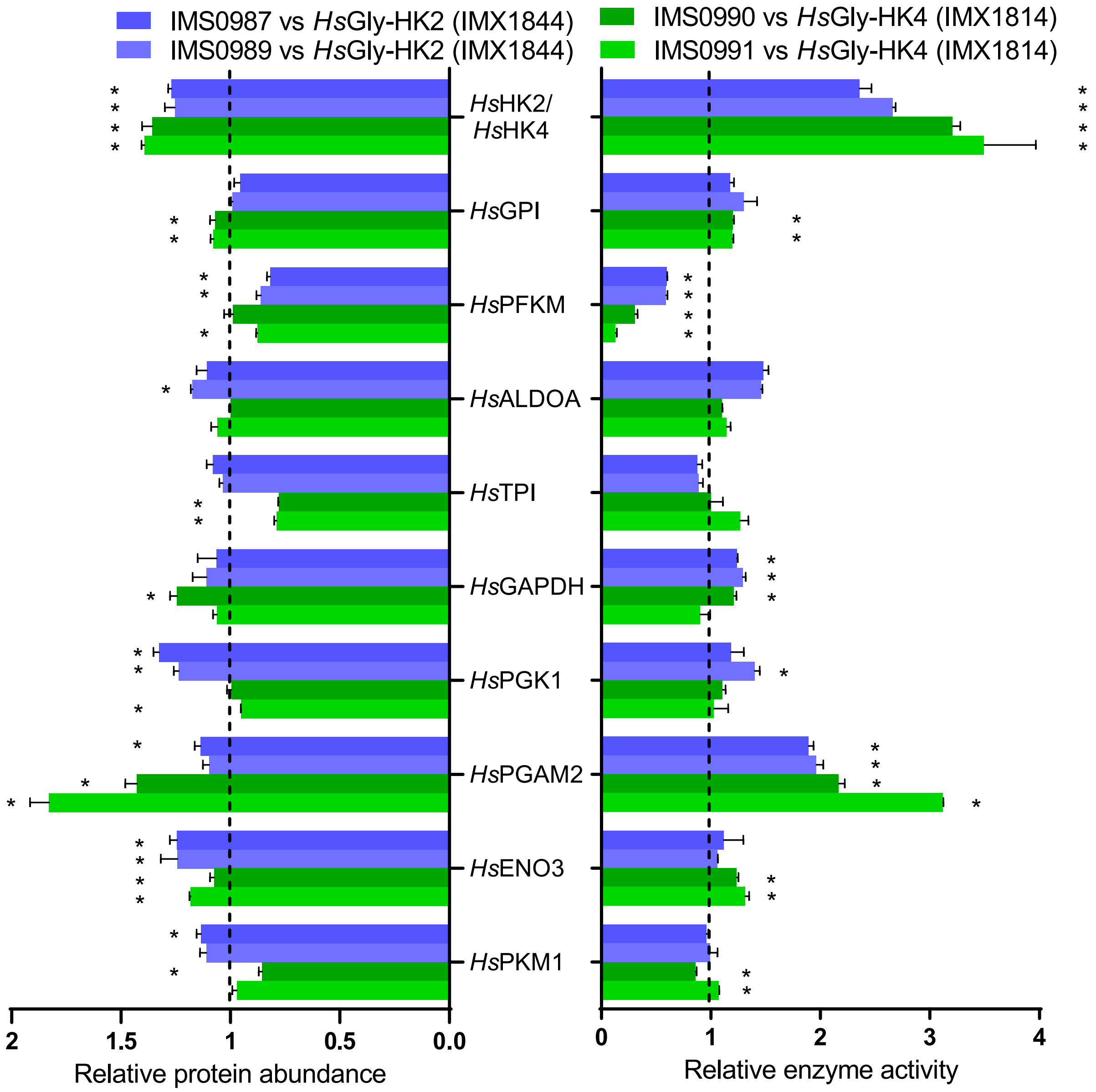


Left side shows relative abundances of the human glycolytic enzymes in the evolved strains relative to unevolved strains IMX1814 and IMX1844. Means with SEM are shown for biological triplicates from one of the duplicate injections. * indicates significant difference with IMX1814/IMX1844 (p-value <0.05 student t-test, two tailed, homoscedastic). Right side shows relative enzyme activities compared to IMX1814/IMX1844. Glycolytic enzymes were assayed from shake flask cultures on SMG at 30˚C. Activities were assayed in 96-well plates. The data represent the average and mean deviation of biological culture duplicates. The asterisks indicate statistical significance (Student t-test, two-tailed, homoscedastic, P<0.05).

#### Figure S14 – Correlation between the change in enzyme activity after evolution of the fully humanized strains and growth rate of single complementation strains

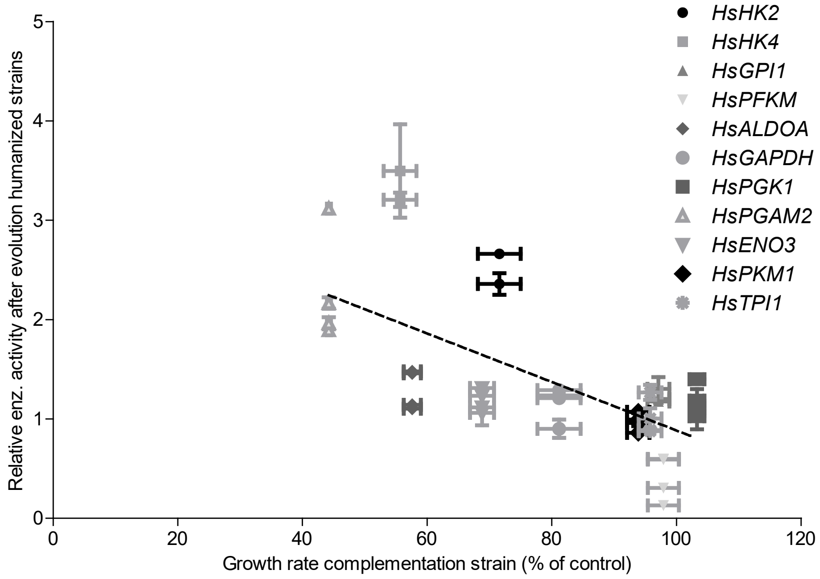


The change in enzyme activity after evolution is plotted against the growth rate of the corresponding complementation strains. Since a low growth rate in the single complementation strains suggests a suboptimal expression and activity, this is expected to correlate with increased activity after evolution. This is the case for several enzymes, especially *PGAM2* and the hexokinases. A linear decreasing trendline is shown ($R^2$ = 0.44). For the hexokinases the values of the complementation strains with *ScHXK2* promoters were used since the same promotor was used in the fully humanized strains

### Figure S15: Global proteome response to evolution of humanized strains

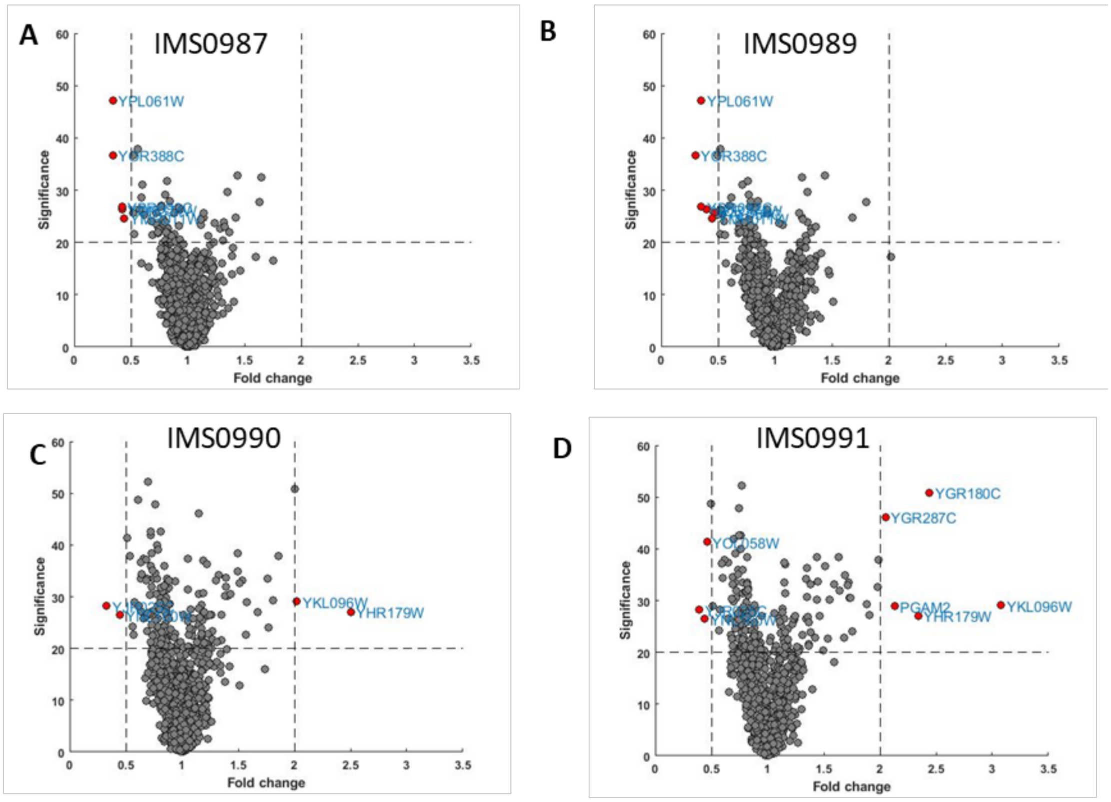


E

| Systematic name | Common name | Fold change vs IMX1844 | P value | TIC normalized abundance IMX1844 |
|---|---|---|---|---|
| IMS0987 | | | | |
| YMR011W | hypothetical protein | 0.44 | 3.5E-03 | 5.5E-04 |
| YOR374W | Ald4 | 0.42 | 2.4E-03 | 1.8E-02 |
| YBR092C | Pho3 | 0.42 | 2.1E-03 | 4.3E-04 |
| YPL061W | Ald6 | 0.34 | 1.9E-05 | 1.9E-02 |
| YOR388C | Fdh1 | 0.34 | 2.2E-04 | 1.2E-03 |
| IMS0989 | | | | |
| YAL054C | Acs1 | 0.46 | 2.8E-03 | 1.3E-03 |
| YMR011W | hypothetical protein | 0.45 | 3.5E-03 | 5.5E-04 |
| YOR374W | Ald4 | 0.4 | 2.4E-03 | 1.8E-02 |
| YPL061W | Ald6 | 0.35 | 1.9E-05 | 1.9E-02 |
| YBR092C | Pho3 | 0.35 | 2.1E-03 | 4.3E-04 |
| YOR388C | Fdh1 | 0.3 | 2.2E-04 | 1.2E-03 |

F

| Systematic name | Common name | Fold change vs IMX1814 | P value | TIC normalized abundance IMX1814 |
|---|---|---|---|---|
| IMS0990 | | | | |
| YHR179W | Oye2 | 2.5 | 2.0E-03 | 2.6E-03 |
| YKL096W | Cwp2 | 2.02 | 1.2E-03 | 2.7E-05 |
| YNL160W | Ygp1/Sps 100 | 0.45 | 2.2E-03 | 4.8E-04 |
| YJR025C | Bna1 | 0.33 | 1.5E-03 | 5.8E-05 |
| IMS0991 | | | | |
| YKL096W | Cwp2 | 3.08 | 1.2E-03 | 2.7E-05 |
| YGR180C | Rnr4/Rnr 2 | 2.44 | 8.3E-06 | 1.0E-03 |
| YHR179W | Oye2 | 2.34 | 2.0E-03 | 2.6E-03 |
| | PGAM2 | 2.13 | 1.3E-03 | 4.0E-04 |
| YGR287C | Ima1 | 2.05 | 2.4E-05 | 4.0E-04 |
| YOL058W | Arg1 | 0.46 | 7.4E-05 | 2.1E-03 |
| YNL160W | Ygp1/Sps 100 | 0.44 | 2.2E-03 | 4.8E-04 |
| YJR025C | Bna1 | 0.39 | 1.5E-03 | 5.8E-05 |

A) - D) Volcano plots showing fold change and significance of the complete set of identified proteins of the evolved strains relative to their parental strain. Proteins that changed more than 2-fold with a significance above 20 (corresponding to a p value of 0.01) are indicated. E-F) Proteins changed > 2-fold in abundance after evolution are indicated with common name, fold change, p-value and normalized abundance in parental strain

#### Figure S16 - Vacuole morphology in evolved humanized yeast strain

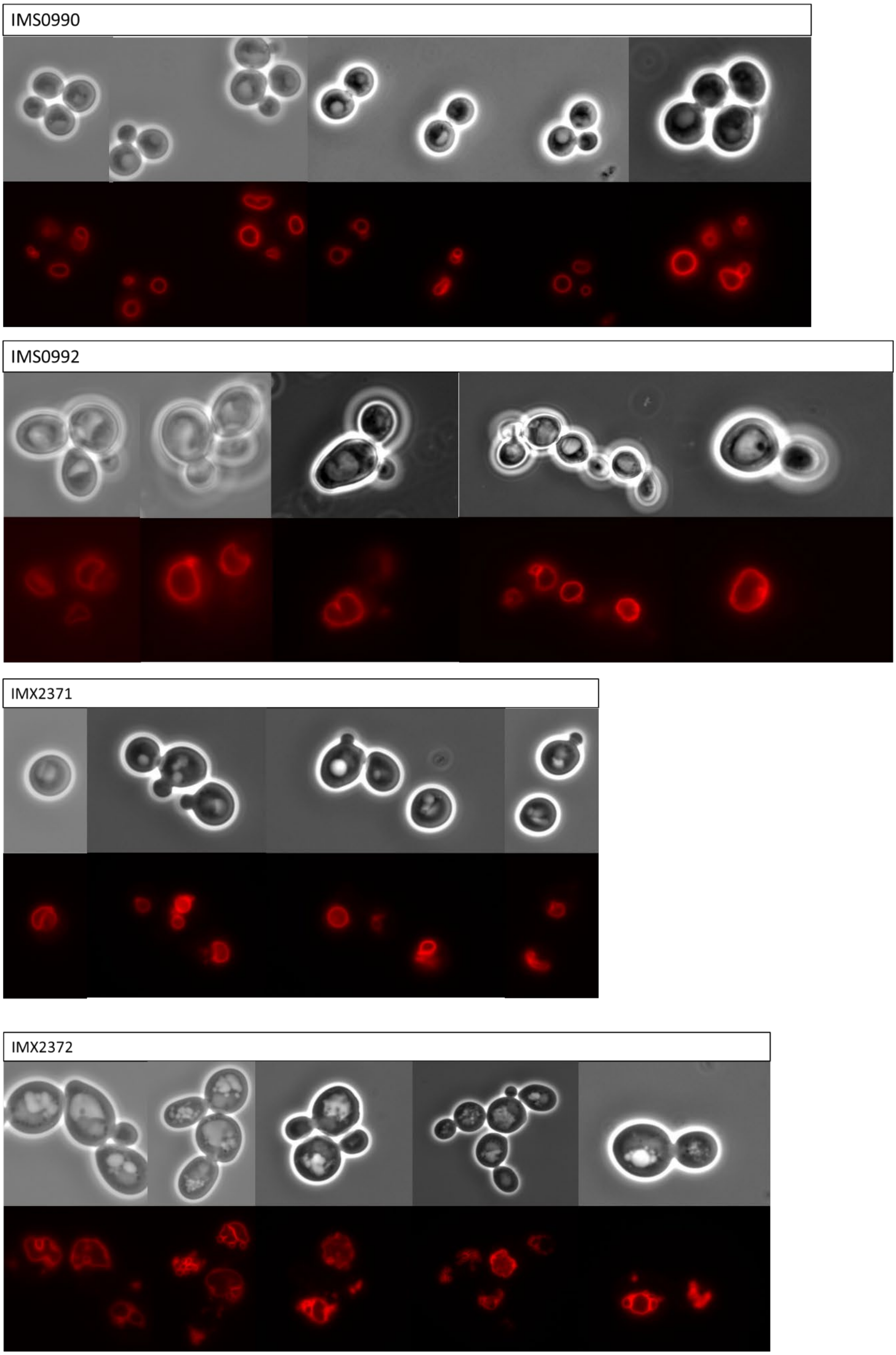

Cells were stained with the red fluorescent dye FM4-64 and visualized with an Imager-Z1 microscope (Carl-Zeiss). Strain IMS0990 and IMS0992 are *Hs*Gly evolved strains and IMX2371 and IMX2372 are the non-evolved, genetically engineered variants in which the *STT4* gene was mutated.

### Figure S17 – Peptide abundance

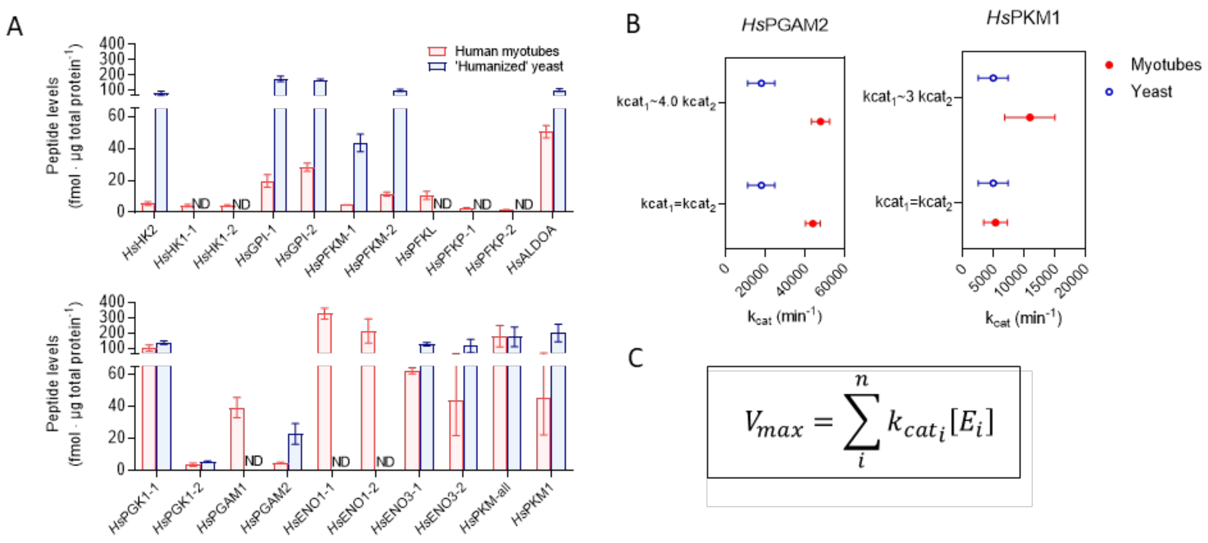


**A)** Peptide abundance in fmol·µg prot$^{-1}$ for the human glycolytic enzymes in yeast (*Hs*Gly-HK2 strain IMX1844, blue) and myotube (red) cell extracts. Peptides identified in the targeted proteomics analysis are represented on the x-axis. When more than one peptide was quantified per protein, they are indicated by -1 or -2 after the protein name (see Table S6 for the peptide sequences). For the myotubes, three independent cultures were averaged, for yeast, two independent cultures were averaged. No absolute quantification could be made for TPI and GAPDH due to the lack of standard peptides. **B)** Estimated $k_{cat}$'s for *Hs*PKM1 and *Hs*PGAM2 with different assumptions, $k_{cat}$ ratios for isozymes derived from the literature. Data show means ± SD. **C)** Equation used to estimate $k_{cat}$, $[E_i]$ represents the concentration of each isoform, number of subunits per enzyme complex not included.

### Figure S18 - Diagnostic PCR of single complementation strains

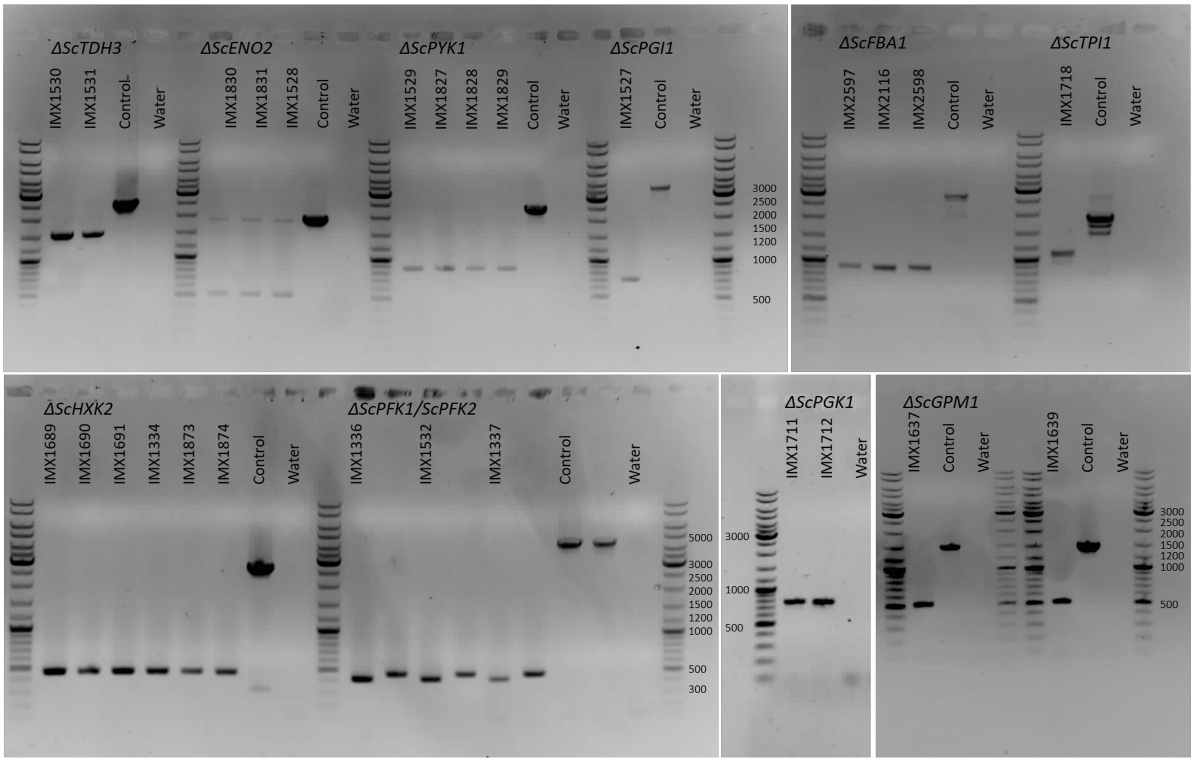


| Yeast gene | Deletion (bp) | No deletion (bp) |
|---|---|---|
| ENO2 | 520 | 1839 |
| PYK1 | 860 | 2373 |
| TDH3 | 1530 | 2534 |
| PGI1 | 710 | 3453 |
| PGK1 | 755 | 2362 |
| GPM1 | 512 | 1481 |
| FBA1 | 865 | 2714 |
| HXK2 | 460 | 2653 |
| TPI1 | 1090 | 1840 |
| PFK1 | 415 | 4418 |
| PFK2 | 375 | 4302 |

PCR confirmation of yeast gene deletions resulting in the individual human gene complementation strains which are indicated per well. The forward and reverse primer were chosen upstream and downstream of the region of gene deletion. The control shows the PCR product resulting from a strain (IMX1076) in which the gene is not deleted. In case of the *ENO2* deletion the primers were binding in the promoter and terminator which resulted in amplification of the human gene as well.

#### Figure S19 – Ploidy analysis of the strains used in this study

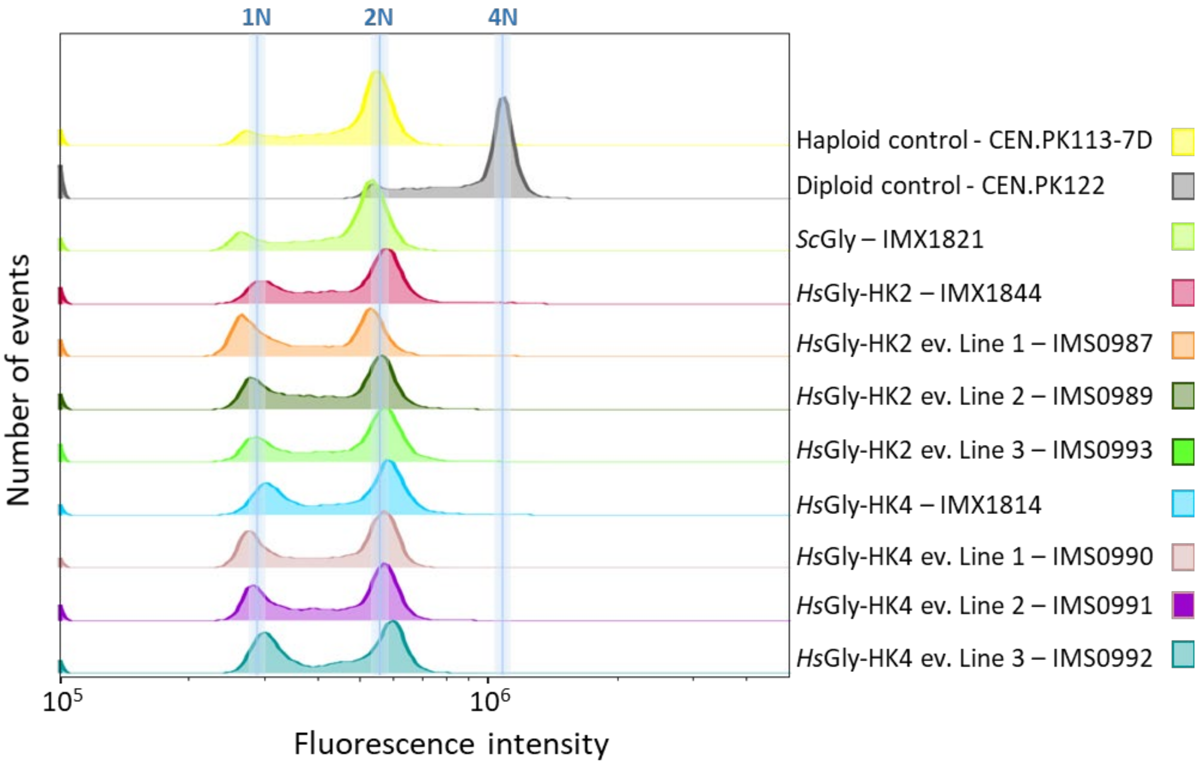


Ploidy measurement based on DNA content determination by flow cytometry. The two top plots represent the haploid and diploid controls. All humanized strains are haploids as expected.

#### Figure S20 - Overview of single locus glycolysis strain construction.

Construction of IMX605 and IMX589 is described in Kuijpers *et al.* 2016.

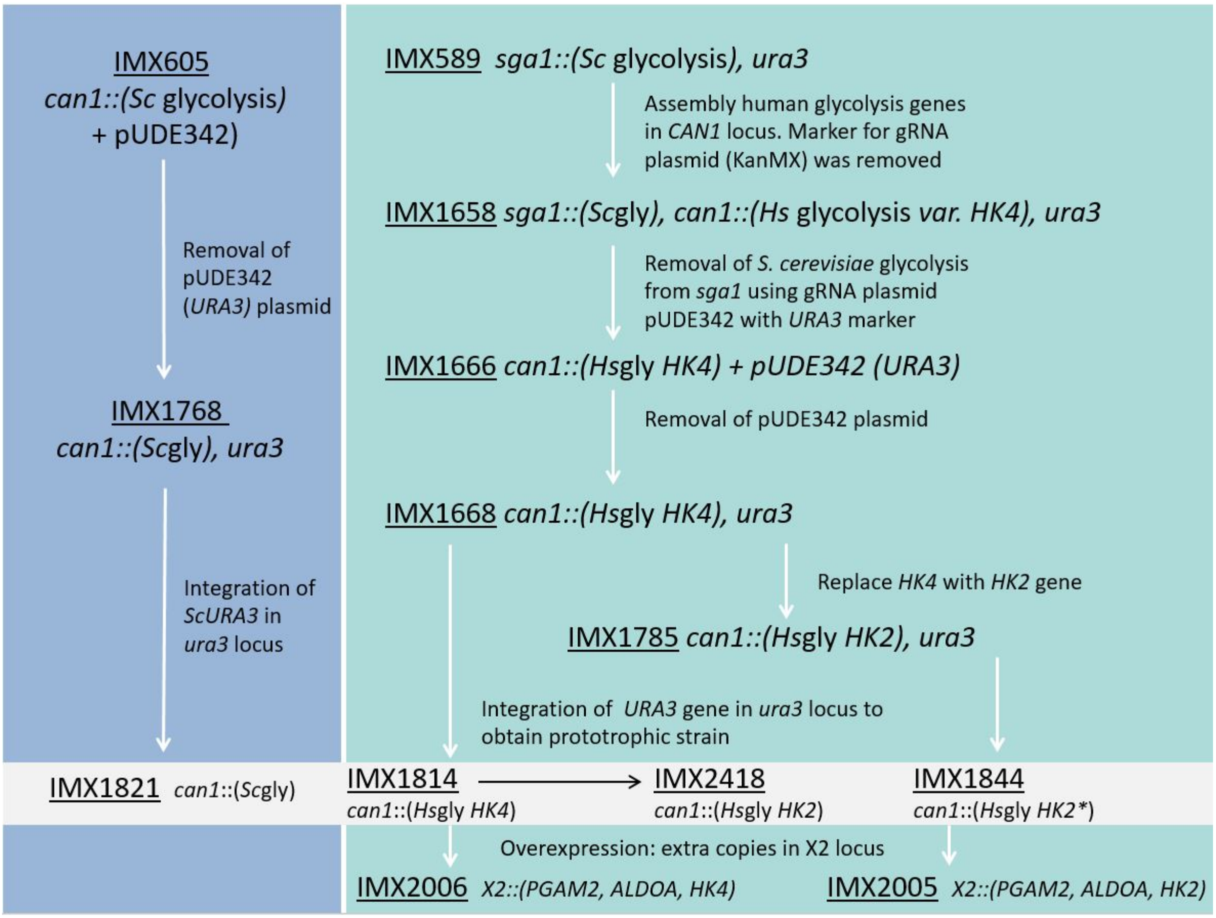

Table S1: Comparison of the human glycolytic proteins to their yeast orthologues

| Human enzyme | Size (aa) | Closest yeast enzyme | Size (aa) | % Identity at protein level | Complementation | Shown previously? |
|---|---|---|---|---|---|---|
| **HK1** | 917 | HXK2 | 486 | 30% to subunit 1, 35% to subunit 2 | Yes | No |
| **HK2** | 917 | HXK2 | 486 | 33% to both subunits | Yes | No |
| **HK3** | 923 | HXK2 | 486 | 28% to subunit 1 33% to subunit 2 | No | No |
| **HK4** | 465 | HXK2 | 486 | 31% | Yes | Yes, [5] |
| **GPI1** | 558 | PGI1 | 554 | 58% | Yes | No |
| **PFKM** | 780 | PFK1, PFK2 | 987 /959 | 43% to PFK1, 43% to PFK2 | Yes | Yes, [6] |
| **PFKP** | 784 | PFK1, PFK2 | 987 /959 | 43% to PFK1, 43% to PFK2 | Yes | No |
| **PFKL** | 780 | PFK1, PFK2 | 987 /959 | 43% to PFK1, 45% to PFK2 | Yes | No |
| **ALDOA** | 364 | FBA1 | 359 | - | Yes | No |
| **ALDOB** | 364 | FBA1 | 359 | - | Yes | Yes, [7] |
| **ALDOC** | 364 | FBA1 | 359 | - | Yes | No |
| **TPI** | 286 | TPI1 | 248 | 53% | Yes | Yes |
| **GAPDH** | 335 | TDH3 | 332 | 65% | Yes | No |
| **GAPDHS** | 408 | TDH3 | 332 | 65% | Yes | No |
| **PGK1** | 417 | PGK1 | 416 | 66% | Yes | Yes, [8] |
| **PGK2** | 417 | PGK1 | 416 | 66% | Yes | Yes, [8] |
| **PGAM1** | 253 | GPM1 | 247 | 51% | Yes | Yes, [8] |
| **PGAM2** | 253 | GPM1 | 247 | 52% | Yes | Tested, but negative, [8] |
| **ENO1** | 434 | ENO2 | 437 | 63% | Yes | No |
| **ENO2** | 434 | ENO2 | 437 | 62% | Yes | No |
| **ENO3** | 434 | ENO2 | 437 | 63% | Yes | No |
| **PKM1*** | 531 | PYK1 | 500 | 52% | Yes | Yes[8] |
| **PKM2*** | 531 | PYK1 | 500 | 52% | Yes | No |
| **PKR#** | 574 | PYK1 | 500 | 50% | Yes | Yes[8] |
| **PKL#** | 553 | PYK1 | 500 | 50% | Yes | Yes[8] |

*,# Splicing variants

#### Table S2: Genetic composition of the glycolytic transcriptional units used for the single complementation strains and strains with fully humanized glycolysis

| Human gene | Yeast prom | Yeast term | Yeast gene | Yeast prom | Yeast term |
|---|---|---|---|---|---|
| *HsHK1* | *PDC1* | *PDC1* | *ScHXK2* | *HXK2* | *HXK2* |
| *HsHK2*[#] | *PDC1* | *PDC1* | | | |
| *HsHK2*[#] | *HXK2* | *HXK2* | | | |
| *HsHK3* | *PDC1* | *PDC1* | | | |
| *HsHK4*[#] | *PDC1* | *PDC1* | | | |
| *HsHK4*[#] | *HXK2* | *HXK2* | | | |
| *HsGPI* | *TEF2* | *TEF2* | *ScPGI1* | *PGI1* | *PGI1* |
| *HsPFKM* | *TEF1* | *TEF1* | *ScPFK1* | *ScPFK1* | *ScPFK1* |
| *HsPFKP* | *TEF1* | *TEF1* | *ScPFK2* | *ScPFK2* | *ScPFK2* |
| *HsPFKL* | *TEF1* | *TEF1* | | | |
| *HsALDOA* | *FBA1* | *FBA1* | *ScFBA1* | *FBA1* | *FBA1* |
| *HsALDOB* | *FBA1* | *FBA1* | | | |
| *HsALDOC* | *FBA1* | *FBA1* | | | |
| *HsTPI* | *TPI1* | *TPI1* | *ScTPI1* | *TPI1* | *TPI1* |
| *HsGAPDH* | *TDH3* | *TDH3* | *ScTDH3* | *TDH3* | *TDH3* |
| *HsGAPDHS* | *TDH3* | *TDH3* | | | |
| *HsPGK1* | *PGK1* | *PGK1* | *ScPGK1* | *PGK1* | *PGK1* |
| *HsPGK2* | *PGK1* | *PGK1* | | | |
| *HsPGAM1* | *GPM1* | *GPM1* | *ScGPM1* | *GPM1* | *GPM1* |
| *HsPGAM2* | *GPM1* | *GPM1* | | | |
| *HsENO1* | *ENO2* | *ENO2* | *ScENO2* | *ENO2* | *ENO2* |
| *HsENO2* | *ENO2* | *ENO2* | | | |
| *HsENO3* | *ENO2* | *ENO2* | | | |
| *HsPKM1* | *PYK1* | *PYK1* | *ScPYK1* | *PYK1* | *PYK1* |
| *HsPKM2* | *PYK1* | *PYK1* | | | |
| *HsPKR* | *PYK1* | *PYK1* | | | |
| *HsPKL* | *PYK1* | *PYK1* | | | |

### For hexokinase expression in full human glycolysis strains *HXK2* promoter and terminator was used and corresponding complementation strains with the same promotor and terminator were constructed.

#### Table S3. Whole genome sequence analysis of the strains with fully humanized glycolysis

| Systematic name | Name | Type | Amino acid change |
|---|---|---|---|
| **Mutations in IMX1666 (human glycolysis var. *HK4*) compared to IMX589 (SwYG)** | | | |
| YDR351W | SBE2 | NSY | Ser-412-Thr |
| YFL064C | uncharacterized protein | NSY | Asn-66-Asp |
| YHR219W | uncharacterized protein | NSY | Val-383-Leu |
| YJL223C | PAU1 | SYN | Ala-13-Ala |
| | *Hs*PFKM | SYN | Val-529-Val |
| - | tPDC1/SHR BC | 4 bp gap | Last 2 bp missing and 2 bp from SHR BC |
| **Mutations in IMX1844 (human glycolysis var. HK2) relative to IMX1666** | | | |
| YJL225C | uncharacterized protein | NSY | Arg-267-Ser |
| - | *Hs*HK2 | NSY | Ile-562-Asn |
| **Mutations in IMX2005 relative to IMX1844** | | | |
| YEL074W | Putative protein | NSY | His-66-Pro |
| **No mutations in IMX2006 relative to IMX1666** | | | |

#### Table S4: Physiology of humanized glycolysis strains in bioreactors

Yields and biomass specific conversion rates of IMX1844 (*Hs*Gly-*HK2*), IMX1814 (*Hs*Gly-*HK4*) and the control strain IMX1821 (*Sc*Gly). Strains were grown in bioreactor at 30°C under aerobic conditions in SMG. Maximum specific growth rate ($\mu_{max}$), biomass specific rates of glucose and oxygen consumption ($q_{glu}$ and $q_{O2}$, respectively), biomass specific rates of production of ethanol ($q_{etoh}$), glycerol ($q_{gly}$), acetate ($q_{ace}$) and carbon dioxide ($q_{CO2}$) and RQ: respiration quotient ($q_{CO2}/q_{O2}$). Yields on glucose for biomass ($Y_{sx}$), ethanol ($Y_{setoh}$), glycerol ($Y_{sgly}$), and acetate ($Y_{sace}$). Values show the average and standard error of the mean from at least two independent bioreactors per strain. DW: biomass dry weight. BDL: below detection level.

| | *Sc*Gly IMX1821 | HsGly-HK2 IMX1844 | HsGly-HK4 IMX1814 |
|---|---|---|---|
| Biomass specific rates | | | |
| $\mu_{max}$ ($h^{-1}$) | 0.32 ± 0.01 | 0.15 ± 0.00 | 0.15 ± 0.00 |
| $-q_{glu}$ ($mmol.g_{DW}^{-1}.h^{-1}$) | 16.1 ± 0.8 | 2.2 ± 0.03 | 3.0 ± 0.44 |
| $q_{etoh}$ ($mmol.g_{DW}^{-1}.h^{-1}$) | 22.3 ± 0.8 | 0.03 ± 0.00 | 0.25 ± 0.07 |
| $q_{gly}$ ($mmol.g_{DW}^{-1}.h^{-1}$) | 1.6 ± 0.1 | 0.11 ± 0.00 | 0.77 ± 0.15 |
| $q_{ace}$ ($mmol.g_{DW}^{-1}.h^{-1}$) | 0.59 ± 0.06 | BDL | 0.58 ± 0.05 |
| Respiration | | | |
| $q_{CO2}$ ($mmol.g_{DW}^{-1}.h^{-1}$) | 29.9 ± 1.6 | 6.8 ± 0.21 | 6.2 ± 0.27 |
| $q_{O2}$ ($mmol.g_{DW}^{-1}.h^{-1}$) | 7.9 ± 0.4 | 6.7 ± 0.31 | 5.4 ± 0.25 |
| RQ | 3.8 ± 0.01 | 1.0 ± 0.01 | 0.87 ± 0.00 |
| Yields | | | |
| $Y_{sx}$ ($g_{DW}/g_{glu}$) | 0.11 ± 0.01 | 0.37 ± 0.01 | 0.28 ± 0.04 |
| $Y_{setoh}$ ($mol_{ethanol}/mol_{glu}$) | 1.4 ± 0.0 | 0.01 ± 0.00 | 0.08 ± 0.01 |
| $Y_{sgly}$ ($mol_{glycerol}/mol_{glu}$) | 0.10 ± 0.00 | 0.05 ± 0.00 | 0.25 ± 0.01 |
| $Y_{sace}$ ($mol_{acetate}/mol_{glu}$) | 0.04 ± 0.00 | BDL | 0.19 ± 0.01 |

#### Table S5 - Mutations in the coding regions of evolved strains

Text in grey italics indicates synonymous mutation. The pink background indicates the mutations common to all six evolved strains and the blue background the mutations common to all evolution lines from a single humanized strain.

| | | IMS0990 | | IMS0991 | | IMS0992 | |
|---|---|---|---|---|---|---|---|
| Evolved strains with IMX1814 (HsGly-HK4) background | TUP1 | G to A | His-489-Tyr | G to A | His-489-Tyr | G to A | His-489-Tyr |
| | STT4 | C to T | Gly-1766-Arg | C to G | Arg-1707-Pro | A to T | Phe-1775-Ile |
| | PFKM | A to G | Thr-81-Ala | G to C | Arg-623-Ser | G to T | Arg-673-Ile |
| | NUT1 | | | T to G | Tyr-432-stop | | |
| | RSM22 | G to A | Arg-120-Cys | | | | |
| | YMR089C | C to A | Pro-217-Thr | G to C | Asp-430-His | | |
| | YCR038W | G to A | Leu-123-Leu | | | | |
| | YLR371W | A to T | Ala-74-Ala | | | | |
| | YKL124W | | | G to A | Thr-346-Thr | | |
| | NMa111 | G to A | Glu-40-Glu | G to A | Glu-40-Glu | G to A | Glu-40-Glu |

| | | IMS0987 | | IMS0989 | | IMS0993 | |
|---|---|---|---|---|---|---|---|
| Evolved strains with IMX1844 (Hx-Gly-HK2) background | STT4 | C to A | Asp-1650-Tyr | A to G | Ile-1771-Thr | G to C | Ser-1611-Cys |
| | PIN4 | A to T | Phe-130-Leu | G to T | Gln-482-stop | | |
| | SNF1 | | | | | T to G | Phe-261-Cys |
| | SKI8 | | | | | G to A | Pro-219-Ser |
| | TAO3 | | | | | G to T | His-167-Asn |
| | CYR1 | G to T | Gly-1612-Val | G to A | Gly-1768-Ser | | |
| | ECM38 | | | | | T to G | Tyr-650-Asp |
| | YOL075C | | | G to C | Pro-246-Ala | | |
| | YME1 | | | | | G to T | STOP-748-Leu |
| | YOR114W | C to A | Pro-146-Pro | | | | |

#### Table S6 – Peptides standards for absolute quantification by targeted proteome analysis

| Protein | Peptide abbreviation | Identified peptide |
|---|---|---|
| *Hs*HK2 | HK2 | LDESFLVSWTK |
| *Hs*HK1 | HK1-1 | FLLSESGSGK |
| | HK1-2 | HIDLVEGDEGR |
| *Hs*GPI | GPI-1 | TFTTQETITNAETAK |
| | GPI-2 | VFEGNRPTNSIVFTK |
| *Hs*PFKM | PFKM-1 | LLAHVRPPVSK |
| | PFKM-2 | VLVVHDGFEGLAK |
| *Hs*PFKL | PFKL | GQVQEVGWHDVAGWLGR |
| *Hs*PFKP | PFKP-1 | VTILGHVQR |
| | PFKP-2 | YLEHLSGDGK |
| *Hs*ALDOA | ALDOA | ALQASALK |
| *Hs*PGK1 | PGK1 | ALESPERPFLAILGGAK |
| | PGK1 | ITLPVDFVTADK |
| *Hs*PGAM1 | PGAM1 | FSGWYDADLSPAGHEEAK |
| *Hs*PGAM2 | PGAM2 | FCGWFDAELSEK |
| *Hs*ENO1 | ENO1-1 | IGAEVYHNLK |
| | ENO1-2 | VNQIGSVTESLQACK |
| *Hs*ENO3 | ENO3-1 | IGAEVYHHLK |
| | ENO3-2 | VNQIGSVTESIQACK |
| *Hs*PKM | PKM | CDENILWLDYK |
| | PKM | CLAAALIVLTESGR |

#### Table S7 – List of strains used in this study

Table S7A: Strains with integrated human gene expression cassette (native yeast ortholog still present)

| Strain | Genotype | Integration plasmid |
|---|---|---|
| IMX1316 | MATa ura3-52 his3-1 leu2-3,112 MAL2-8c SUC2 glk1::SpHis5, hxk1::KlLEU2, tdh1, tdh2, gpm2, gpm3, eno1, pyk2, pdc5, pdc6, adh2, adh5, adh4, sga1::(CAS9, NatNT), *ura3::(pPDC1-HK1(hs)-tPDC1, URA3)* | pUDI133 |
| IMX1317 | MATa ura3-52 his3-1 leu2-3,112 MAL2-8c SUC2 glk1::SpHis5, hxk1::KlLEU2, tdh1, tdh2, gpm2, gpm3, eno1, pyk2, pdc5, pdc6, adh2, adh5, adh4, sga1::(CAS9, NatNT), ura3::(pPDC1-HK2(hs)-tPDC1, URA3) | pUDI134 |
| IMX1318 | MATa ura3-52 his3-1 leu2-3,112 MAL2-8c SUC2 glk1::SpHis5, hxk1::KlLEU2, tdh1, tdh2, gpm2, gpm3, eno1, pyk2, pdc5, pdc6, adh2, adh5, adh4, sga1::(CAS9, NatNT), ura3::(pPDC1-HK3(hs)-tPDC1, URA3) | pUDI135 |
| IMX1319 | MATa ura3-52 his3-1 leu2-3,112 MAL2-8c SUC2 glk1::SpHis5, hxk1::KlLEU2, tdh1, tdh2, gpm2, gpm3, eno1, pyk2, pdc5, pdc6, adh2, adh5, adh4, sga1::(CAS9, NatNT) , ura3::(pPDC1-HK4(hs)-tPDC1, URA3) | pUDI136 |
| IMX1838 | MATa ura3-52 his3-1 leu2-3,112 MAL2-8c SUC2 glk1::SpHis5, hxk1::KlLEU2, tdh1, tdh2, gpm2, gpm3, eno1, pyk2, pdc5, pdc6, adh2, adh5, adh4, sga1::(CAS9, NatNT), ura3::(pHXK2-HK2(hs)-tHXK2, URA3) | pUDI206 |
| IMX1839 | MATa ura3-52 his3-1 leu2-3,112 MAL2-8c SUC2 glk1::SpHis5, hxk1::KlLEU2, tdh1, tdh2, gpm2, gpm3, eno1, pyk2, pdc5, pdc6, adh2, adh5, adh4, sga1::(CAS9, NatNT) , ura3::(pHXK2-HK4(hs)-tHXK2, URA3) | pUDI207 |

| | | |
|---|---|---|
| IMX1320 | MATa ura3-52 his3-1 leu2-3,112 MAL2-8c SUC2 glk1::SpHis5, hxk1::KlLEU2, tdh1, tdh2, gpm2, gpm3, eno1, pyk2, pdc5, pdc6, adh2, adh5, adh4, sga1::(CAS9, NatNT) , ura3::(pTEF2-GPI1(hs)-tTEF2, URA3) | pUDI137 |
| IMX1321 | MATa ura3-52 his3-1 leu2-3,112 MAL2-8c SUC2 glk1::SpHis5, hxk1::KlLEU2, tdh1, tdh2, gpm2, gpm3, eno1, pyk2, pdc5, pdc6, adh2, adh5, adh4, sga1::(CAS9, NatNT) , ura3::(pTEF1-PFKM(hs)-tTEF1, URA3) | pUDI138 |
| IMX1322 | MATa ura3-52 his3-1 leu2-3,112 MAL2-8c SUC2 glk1::SpHis5, hxk1::KlLEU2, tdh1, tdh2, gpm2, gpm3, eno1, pyk2, pdc5, pdc6, adh2, adh5, adh4, sga1::(CAS9, NatNT) , ura3::(pTEF1-PFKP(hs)-tTEF1, URA3) | pUDI139 |
| IMX1323 | MATa ura3-52 his3-1 leu2-3,112 MAL2-8c SUC2 glk1::SpHis5, hxk1::KlLEU2, tdh1, tdh2, gpm2, gpm3, eno1, pyk2, pdc5, pdc6, adh2, adh5, adh4, sga1::(CAS9, NatNT) , ura3::(pTEF1-PFKL(hs)-tTEF1, URA3) | pUDI140 |
| IMX2585 | MATa ura3-52 his3-1 leu2-3,112 MAL2-8c SUC2 glk1::SpHis5, hxk1::KlLEU2, tdh1, tdh2, gpm2, gpm3, eno1, pyk2, pdc5, pdc6, adh2, adh5, adh4, sga1::(CAS9, NatNT), ura3::pFBA1-ALDOA(hs)-tFBA1, URA3) | pUDI141 |
| IMX1354 | MATa ura3-52 his3-1 leu2-3,112 MAL2-8c SUC2 glk1::SpHis5, hxk1::KlLEU2, tdh1, tdh2, gpm2, gpm3, eno1, pyk2, pdc5, pdc6, adh2, adh5, adh4, sga1::(CAS9, NatNT), ura3::pFBA1-ALDOB(hs)-tFBA1, URA3) | pUDI142 |
| IMX2586 | MATa ura3-52 his3-1 leu2-3,112 MAL2-8c SUC2 glk1::SpHis5, hxk1::KlLEU2, tdh1, tdh2, gpm2, gpm3, eno1, pyk2, pdc5, pdc6, adh2, adh5, adh4, sga1::(CAS9, NatNT), ura3::pFBA1-ALDOC(hs)-tFBA1, URA3) | pUDI143 |
| IMX1692 | MATa ura3-52 his3-1 leu2-3,112 MAL2-8c SUC2 glk1::SpHis5, hxk1::KlLEU2, tdh1, tdh2, gpm2, gpm3, eno1, pyk2, pdc5, pdc6, adh2, adh5, adh4, sga1::(CAS9, NatNT), ura3::(pTPI1-TPI1(hs)-tTPI1, URA3) | pUDI144 |

| | | |
|---|---|---|
| IMX1356 | MATa ura3-52 his3-1 leu2-3,112 MAL2-8c SUC2 glk1::SpHis5, hxk1::KlLEU2, tdh1, tdh2, gpm2, gpm3, eno1, pyk2, pdc5, pdc6, adh2, adh5, adh4, sga1::(CAS9, NatNT), ura3::pTDH3-GAPDH(hs)-tTDH3, URA3) | pUDI145 |
| IMX1300 | MATa ura3-52 his3-1 leu2-3,112 MAL2-8c SUC2 glk1::SpHis5, hxk1::KlLEU2, tdh1, tdh2, gpm2, gpm3, eno1, pyk2, pdc5, pdc6, adh2, adh5, adh4, sga1::(CAS9, NatNT), ura3::pTDH3-GAPDHS(hs)-tTDH3, URA3) | pUDI146 |
| IMX1301 | MATa ura3-52 his3-1 leu2-3,112 MAL2-8c SUC2 glk1::SpHis5, hxk1::KlLEU2, tdh1, tdh2, gpm2, gpm3, eno1, pyk2, pdc5, pdc6, adh2, adh5, adh4, sga1::(CAS9, NatNT), ura3::pPGK1-PGK1(hs)-tPGK1, URA3) | pUDI147 |
| IMX1302 | MATa ura3-52 his3-1 leu2-3,112 MAL2-8c SUC2 glk1::SpHis5, hxk1::KlLEU2, tdh1, tdh2, gpm2, gpm3, eno1, pyk2, pdc5, pdc6, adh2, adh5, adh4, sga1::(CAS9, NatNT), ura3::pPGK1-PGK2(hs)-tPGK1, URA3) | pUDI148 |
| IMX1303 | MATa ura3-52 his3-1 leu2-3,112 MAL2-8c SUC2 glk1::SpHis5, hxk1::KlLEU2, tdh1, tdh2, gpm2, gpm3, eno1, pyk2, pdc5, pdc6, adh2, adh5, adh4, sga1::(CAS9, NatNT), ura3::pGPM1-PGAM1(hs)-tGPM1, URA3) | pUDI149 |
| IMX1304 | MATa ura3-52 his3-1 leu2-3,112 MAL2-8c SUC2 glk1::SpHis5, hxk1::KlLEU2, tdh1, tdh2, gpm2, gpm3, eno1, pyk2, pdc5, pdc6, adh2, adh5, adh4, sga1::(CAS9, NatNT), ura3::pGPM1-PGAM2(hs)-tGPM1, URA3) | pUDI150 |
| IMX1305 | MATa ura3-52 his3-1 leu2-3,112 MAL2-8c SUC2 glk1::SpHis5, hxk1::KlLEU2, tdh1, tdh2, gpm2, gpm3, eno1, pyk2, pdc5, pdc6, adh2, adh5, adh4, sga1::(CAS9, NatNT), ura3::pENO2-ENO1(hs)-tENO2, URA3) | pUDI151 |
| IMX1306 | MATa ura3-52 his3-1 leu2-3,112 MAL2-8c SUC2 glk1::SpHis5, hxk1::KlLEU2, tdh1, tdh2, gpm2, gpm3, eno1, pyk2, pdc5, pdc6, adh2, adh5, adh4, sga1::(CAS9, NatNT), ura3::pENO2-ENO2(hs)-tENO2, URA3) | pUDI152 |

| | | |
|---|---|---|
| IMX1307 | MATa ura3-52 his3-1 leu2-3,112 MAL2-8c SUC2 glk1::SpHis5, hxk1::KlLEU2, tdh1, tdh2, gpm2, gpm3, eno1, pyk2, pdc5, pdc6, adh2, adh5, adh4, sga1::(CAS9, NatNT), ura3::pENO2-ENO3(hs)-tENO2, URA3) | pUDI153 |
| IMX1387 | MATa ura3-52 his3-1 leu2-3,112 MAL2-8c SUC2 glk1::SpHis5, hxk1::KlLEU2, tdh1, tdh2, gpm2, gpm3, eno1, pyk2, pdc5, pdc6, adh2, adh5, adh4, sga1::(CAS9, NatNT), ura3::pPYK1-PKM1(hs)-tPYK1, URA3) | pUDI154 |
| IMX1388 | MATa ura3-52 his3-1 leu2-3,112 MAL2-8c SUC2 glk1::SpHis5, hxk1::KlLEU2, tdh1, tdh2, gpm2, gpm3, eno1, pyk2, pdc5, pdc6, adh2, adh5, adh4, sga1::(CAS9, NatNT), ura3::pPYK1-PKM2(hs)-tPYK1, URA3) | pUDI155 |
| IMX1389 | MATa ura3-52 his3-1 leu2-3,112 MAL2-8c SUC2 glk1::SpHis5, hxk1::KlLEU2, tdh1, tdh2, gpm2, gpm3, eno1, pyk2, pdc5, pdc6, adh2, adh5, adh4, sga1::(CAS9, NatNT), ura3::pPYK1-PKL(hs)-tPYK1, URA3) | pUDI156 |
| IMX1693 | MATa ura3-52 his3-1 leu2-3,112 MAL2-8c SUC2 glk1::SpHis5, hxk1::KlLEU2, tdh1, tdh2, gpm2, gpm3, eno1, pyk2, pdc5, pdc6, adh2, adh5, adh4, sga1::(CAS9, NatNT), ura3::(pPGK1-PKR(hs)-tPGK1, URA3) | pUDI157 |

Table S7B: Individual complementation strains

| Strain | Genotype | Modification |
| --- | --- | --- |
| IMX1689 | MATa ura3-52 his3-1 leu2-3112 MAL2-8c SUC2 glk1::SpHis5 hxk1::KlLEU2 tdh1 tdh2 gpm2 gpm3 eno1 pyk2 pdc5 pdc6 adh2 adh5 adh4 sga1::(CAS9 NatNT) ura3::(pPDC1-HK1(hs)-tPDC1 URA3) hxk2 | ScHXK2 deletion |
| IMS1137 | MATa ura3-52 his3-1 leu2-3112 MAL2-8c SUC2 glk1::SpHis5 hxk1::KlLEU2 tdh1 tdh2 gpm2 gpm3 eno1 pyk2 pdc5 pdc6 adh2 adh5 adh4 sga1::(CAS9 NatNT) ura3::(pPDC1-HK1*(hs)-tPDC1 URA3) hxk2 | Growth on glucose |
| IMS1140 | MATa ura3-52 his3-1 leu2-3112 MAL2-8c SUC2 glk1::SpHis5 hxk1::KlLEU2 tdh1 tdh2 gpm2 gpm3 eno1 pyk2 pdc5 pdc6 adh2 adh5 adh4 sga1::(CAS9 NatNT) ura3::(pPDC1-HK1*(hs)-tPDC1 URA3) hxk2 | Growth on glucose |
| IMS1143 | MATa ura3-52 his3-1 leu2-3112 MAL2-8c SUC2 glk1::SpHis5 hxk1::KlLEU2 tdh1 tdh2 gpm2 gpm3 eno1 pyk2 pdc5 pdc6 adh2 adh5 adh4 sga1::(CAS9 NatNT) ura3::(pPDC1-HK1*(hs)-tPDC1 URA3) hxk2 | Growth on glucose |
| IMX1690 | MATa ura3-52 his3-1 leu2-3112 MAL2-8c SUC2 glk1::SpHis5 hxk1::KlLEU2 tdh1 tdh2 gpm2 gpm3 eno1 pyk2 pdc5 pdc6 adh2 adh5 adh4 sga1::(CAS9 NatNT) ura3::(pPDC1-HK2*(hs)-tPDC1 URA3) hxk2 | ScHXK2 deletion growth on glucose |
| IMX2419 | MATa ura3-52 his3-1 leu2-3112 MAL2-8c SUC2 glk1::SpHis5 hxk1::KlLEU2 tdh1 tdh2 gpm2 gpm3 eno1 pyk2 pdc5 pdc6 adh2 adh5 adh4 sga1::(CAS9 NatNT) ura3::(pPDC1-HK2(hs)-tPDC1 URA3) hxk2 | ScHXK2 deletion |

| | | |
|---|---|---|
| IMX1691 | MATa ura3-52 his3-1 leu2-3112 MAL2-8c SUC2 glk1::SpHis5 hxk1::KlLEU2 tdh1 tdh2 gpm2 gpm3 eno1 pyk2 pdc5 pdc6 adh2 adh5 adh4 sga1::(CAS9 NatNT) ura3::(pPDC1-HK3(hs)-tPDC1 URA3) hxk2 | ScHXK2 deletion |
| IMX1334 | MATa ura3-52 his3-1 leu2-3112 MAL2-8c SUC2 glk1::SpHis5 hxk1::KlLEU2 tdh1 tdh2 gpm2 gpm3 eno1 pyk2 pdc5 pdc6 adh2 adh5 adh4 sga1::(CAS9 NatNT) ura3::(pPDC1-HK4(hs)-tPDC1 URA3) hxk2 | ScHXK2 deletion |
| IMX1873 | MATa ura3-52 his3-1 leu2-3112 MAL2-8c SUC2 glk1::SpHis5 hxk1::KlLEU2 tdh1 tdh2 gpm2 gpm3 eno1 pyk2 pdc5 pdc6 adh2 adh5 adh4 sga1::(CAS9 NatNT) ura3::(pHXK2-HK2(hs)-tHXK2 URA3) hxk2 | ScHXK2 deletion |
| IMX1874 | MATa ura3-52 his3-1 leu2-3112 MAL2-8c SUC2 glk1::SpHis5 hxk1::KlLEU2 tdh1 tdh2 gpm2 gpm3 eno1 pyk2 pdc5 pdc6 adh2 adh5 adh4 sga1::(CAS9 NatNT) ura3::(pHXK2-HK4(hs)-tHXK2 URA3) hxk2 | ScHXK2 deletion |
| IMX1527 | MATa ura3-52 his3-1 leu2-3112 MAL2-8c SUC2 glk1::SpHis5 hxk1::KlLEU2 tdh1 tdh2 gpm2 gpm3 eno1 pyk2 pdc5 pdc6 adh2 adh5 adh4 sga1::(CAS9 NatNT) ura3::(pTEF2-GPI1(hs)-tTEF2 URA3) pgi1 | ScPGI1 deletion |
| IMX1336 | MATa ura3-52 his3-1 leu2-3112 MAL2-8c SUC2 glk1::SpHis5 hxk1::KlLEU2 tdh1 tdh2 gpm2 gpm3 eno1 pyk2 pdc5 pdc6 adh2 adh5 adh4 sga1::(CAS9 NatNT) ura3::(pTEF1-PFKM(hs)-tTEF1 URA3) pfk1 pfk2 | ScPFK1 and PFK2 deletion |
| IMX1532 | MATa ura3-52 his3-1 leu2-3112 MAL2-8c SUC2 glk1::SpHis5 hxk1::KlLEU2 tdh1 tdh2 gpm2 gpm3 eno1 pyk2 pdc5 pdc6 adh2 adh5 adh4 sga1::(CAS9 NatNT) ura3::(pTEF1-PFKL(hs)-tTEF1 URA3) pfk1 pfk2 | ScPFK1 and PFK2 deletion |
| IMX1337 | MATa ura3-52 his3-1 leu2-3112 MAL2-8c SUC2 glk1::SpHis5 hxk1::KlLEU2 tdh1 tdh2 gpm2 gpm3 eno1 pyk2 pdc5 pdc6 adh2 adh5 adh4 sga1::(CAS9 NatNT) ura3::(pTEF1-PFKP(hs)-tTEF1 URA3) pfk1 pfk2 | ScPFK1 and PFK2 deletion |

| | | |
|---|---|---|
| IMX2597 | MATa ura3-52 his3-1 leu2-3112 MAL2-8c SUC2 glk1::SpHis5 hxk1::KlLEU2 tdh1 tdh2 gpm2 gpm3 eno1 pyk2 pdc5 pdc6 adh2 adh5 adh4 sga1::(CAS9 NatNT) ura3::(pFBA1-ALDOA(hs)-tFBA1 URA3) fba1 | ScFBA1 deletion |
| IMX2116 | MATa ura3-52 his3-1 leu2-3112 MAL2-8c SUC2 glk1::SpHis5 hxk1::KlLEU2 tdh1 tdh2 gpm2 gpm3 eno1 pyk2 pdc5 pdc6 adh2 adh5 adh4 sga1::(CAS9 NatNT) ura3::(pFBA1-ALDOB(hs)-tFBA1 URA3) fba1 | ScFBA1 deletion |
| IMX2598 | MATa ura3-52 his3-1 leu2-3112 MAL2-8c SUC2 glk1::SpHis5 hxk1::KlLEU2 tdh1 tdh2 gpm2 gpm3 eno1 pyk2 pdc5 pdc6 adh2 adh5 adh4 sga1::(CAS9 NatNT) ura3::(pFBA1-ALDOC(hs)-tFBA1 URA3) fba1 | ScFBA1 deletion |
| IMX1718 | MATa ura3-52 his3-1 leu2-3112 MAL2-8c SUC2 glk1::SpHis5 hxk1::KlLEU2 tdh1 tdh2 gpm2 gpm3 eno1 pyk2 pdc5 pdc6 adh2 adh5 adh4 sga1::(CAS9 NatNT) ura3::(pTPI1-TPI1(hs)-tTPI1 URA3) tpi1 | ScTPI1 deletion |
| IMX1530 | MATa ura3-52 his3-1 leu2-3112 MAL2-8c SUC2 glk1::SpHis5 hxk1::KlLEU2 tdh1 tdh2 gpm2 gpm3 eno1 pyk2 pdc5 pdc6 adh2 adh5 adh4 sga1::(CAS9 NatNT) ura3::pTDH3-GAPDH(hs)-tTDH3 URA3) tdh3 | ScTDH3 deletion |
| IMX1531 | MATa ura3-52 his3-1 leu2-3112 MAL2-8c SUC2 glk1::SpHis5 hxk1::KlLEU2 tdh1 tdh2 gpm2 gpm3 eno1 pyk2 pdc5 pdc6 adh2 adh5 adh4 sga1::(CAS9 NatNT) ura3::pTDH3-GAPDHS(hs)-tTDH3 URA3) tdh3 | ScTDH3 deletion |
| IMX1711 | MATa ura3-52 his3-1 leu2-3112 MAL2-8c SUC2 glk1::SpHis5 hxk1::KlLEU2 tdh1 tdh2 gpm2 gpm3 eno1 pyk2 pdc5 pdc6 adh2 adh5 adh4 sga1::(CAS9 NatNT) ura3::(pPGK1-PGK1(hs)-tPGK1 URA3) pgk1 | ScPGK1 deletion |
| IMX1712 | MATa ura3-52 his3-1 leu2-3112 MAL2-8c SUC2 glk1::SpHis5 hxk1::KlLEU2 tdh1 tdh2 gpm2 gpm3 eno1 pyk2 pdc5 pdc6 adh2 adh5 adh4 sga1::(CAS9 NatNT) ura3::(pPGK1-PGK2(hs)-tPGK1 URA3) pgk1 | ScPGK1 deletion |

| | | |
|---|---|---|
| IMX1637 | MATa ura3-52 his3-1 leu2-3112 MAL2-8c SUC2 glk1::SpHis5 hxk1::KlLEU2 tdh1 tdh2 gpm2 gpm3 eno1 pyk2 pdc5 pdc6 adh2 adh5 adh4 sga1::(CAS9 NatNT) ura3::(pGPM1-PGAM1(hs)-tGPM1 URA3) gpm1 | ScGPM1 deletion |
| IMX1639 | MATa ura3-52 his3-1 leu2-3112 MAL2-8c SUC2 glk1::SpHis5 hxk1::KlLEU2 tdh1 tdh2 gpm2 gpm3 eno1 pyk2 pdc5 pdc6 adh2 adh5 adh4 sga1::(CAS9 NatNT) ura3::(pGPM1-PGAM2(hs)-tGPM1 URA3) gpm1 | ScGPM1 deletion |
| IMX1830 | MATa ura3-52 his3-1 leu2-3112 MAL2-8c SUC2 glk1::SpHis5 hxk1::KlLEU2 tdh1 tdh2 gpm2 gpm3 eno1 pyk2 pdc5 pdc6 adh2 adh5 adh4 sga1::(CAS9 NatNT) ura3::pENO2-ENO1(hs)-tENO2 URA3) eno2 | ScENO2 deletion |
| IMX1831 | MATa ura3-52 his3-1 leu2-3112 MAL2-8c SUC2 glk1::SpHis5 hxk1::KlLEU2 tdh1 tdh2 gpm2 gpm3 eno1 pyk2 pdc5 pdc6 adh2 adh5 adh4 sga1::(CAS9 NatNT) ura3::pENO2-PENO2(hs)-tENO2 URA3) eno2 | ScENO2 deletion |
| IMX1528 | MATa ura3-52 his3-1 leu2-3112 MAL2-8c SUC2 glk1::SpHis5 hxk1::KlLEU2 tdh1 tdh2 gpm2 gpm3 eno1 pyk2 pdc5 pdc6 adh2 adh5 adh4 sga1::(CAS9 NatNT) ura3::pENO2-ENO3(hs)-tENO2 URA3) eno2 | ScENO2 deletion |
| IMX1529 | MATa ura3-52 his3-1 leu2-3112 MAL2-8c SUC2 glk1::SpHis5 hxk1::KlLEU2 tdh1 tdh2 gpm2 gpm3 eno1 pyk2 pdc5 pdc6 adh2 adh5 adh4 sga1::(CAS9 NatNT) ura3::pPYK1-PKM1(hs)-tPYK1 URA3) pyk1 | ScPYK1 deletion |
| IMX1827 | MATa ura3-52 his3-1 leu2-3112 MAL2-8c SUC2 glk1::SpHis5 hxk1::KlLEU2 tdh1 tdh2 gpm2 gpm3 eno1 pyk2 pdc5 pdc6 adh2 adh5 adh4 sga1::(CAS9 NatNT) ura3::pPYK1-PKM2(hs)-tPYK1 URA3) pyk1 | ScPYK1 deletion |
| IMX1828 | MATa ura3-52 his3-1 leu2-3112 MAL2-8c SUC2 glk1::SpHis5 hxk1::KlLEU2 tdh1 tdh2 gpm2 gpm3 eno1 pyk2 pdc5 pdc6 adh2 adh5 adh4 sga1::(CAS9 NatNT) ura3::pPYK1-PKL(hs)-tPYK1 URA3) pyk1 | ScPYK1 deletion |

| | | |
|---|---|---|
| IMX1829 | MATa ura3-52 his3-1 leu2-3112 MAL2-8c SUC2 glk1::SpHis5 hxk1::KlLEU2 tdh1 tdh2 gpm2 gpm3 eno1 pyk2 pdc5 pdc6 adh2 adh5 adh4 sga1::(CAS9 NatNT) ura3::pPYK1-PKR(hs)-tPYK1 URA3) pyk1 | ScPYK1 deletion |

Table S7C: Single locus glycolysis strains

| Strain | Genotype | Description/ Modification |
|---|---|---|
| IMX605 | MATa ura3-52 his3-1 leu2-3,112 MAL2-8c SUC2 glk1::Sphis5 hxk1::KlLEU2 tdh1 tdh2 gpm2 gpm3 eno1 pyk2 pdc5 pdc6 adh2 adh5 adh4 sga1 pyk1 pgi1 tpi1 tdh3 pfk2::(pTEF-cas9-tCYC1 natNT1) pgk1 gpm1 fba1 hxk2 pfk1 adh1 pdc1 eno2 can1::(FBA1$_{H}$TPI1$_{P}$PGK1$_{Q}$ADH1$_{N}$PYK1$_{O}$ TDH3$_{A}$ENO2$_{B}$HXK2$_{C}$PGI1$_{D}$PFK1$_{J}$PFK2$_{K}$KanMX$_{L}$GPM1$_{M}$PDC1) pUDE342 | Yeast glycolysis in *can1 + pUDE342* [9] |
| IMX1768 | MATa ura3-52 his3-1 leu2-3,112 MAL2-8c SUC2 glk1::Sphis5 hxk1::KlLEU2 tdh1 tdh2 gpm2 gpm3 eno1 pyk2 pdc5 pdc6 adh2 adh5 adh4 sga1 pyk1 pgi1 tpi1 tdh3 pfk2::(pTEF-cas9-tCYC1 natNT1) pgk1 gpm1 fba1 hxk2 pfk1 adh1 pdc1 eno2 can1::( FBA1$_{H}$TPI1$_{P}$PGK1$_{Q}$ADH1$_{N}$PYK1$_{O}$ TDH3$_{A}$ENO2$_{B}$HXK2$_{C}$PGI1$_{D}$PFK1$_{J}$PFK2$_{K}$KanMX$_{L}$GPM1$_{M}$PDC1) | Yeast glycolysis in *can1* |
| IMX1821 | MATa ura3-52 his3-1 leu2-3,112 MAL2-8c SUC2 glk1::Sphis5 hxk1::KlLEU2 tdh1::ScURA3 tdh2 gpm2 gpm3 eno1 pyk2 pdc5 pdc6 adh2 adh5 adh4 sga1 pyk1 pgi1 tpi1 tdh3 pfk2::(pTEF-cas9-tCYC1 natNT1) pgk1 gpm1 fba1 hxk2 pfk1 adh1 pdc1 eno2 can1::( FBA1$_{H}$TPI1$_{P}$PGK1$_{Q}$ADH1$_{N}$PYK1$_{O}$ TDH3$_{A}$ENO2$_{B}$HXK2$_{C}$PGI1$_{D}$PFK1$_{J}$PFK2$_{K}$KanMX$_{L}$GPM1$_{M}$PDC1) | Yeast glycolysis in *can1 + ScURA3* in *tdh1* |
| IMX589 | MATa ura3-52 his3-1 leu2-3,112 MAL2-8c SUC2 glk1::HIS5 hxk1::LEU2 tdh1::URA3 tdh2 gpm2::LoxP gpm3 eno1 pyk2 pdc5 pdc6 adh2 adh5 adh4 sga1::( FBA1$_{H}$TPI1$_{P}$PGK1$_{Q}$ADH1$_{N}$PYK1$_{O}$ | Yeast glycolysis in *sga1* (SwYG) [9] |

| | | |
|---|---|---|
| | TDH3$_{A}$ENO2$_{B}$HXK2$_{C}$PGI1$_{D}$PFK1$_{J}$PFK2$_{K}$AmdSYM$_{L}$GPM1$_{M}$PDC1) pyk1 pgi1 tpi1 tdh3 pfk2::(pTEFCAS9 nat) pgk1 gpm1 fba1 hxk2 pfk1 adh1 pdc1 eno2 | |
| IMX1658 | MATa ura3-52 his3-1 leu2-3,112 MAL2-8c SUC2 glk1::HIS5 hxk1::LEU2 tdh1::URA3 tdh2 gpm2::LoxP gpm3 eno1 pyk2 pdc5 pdc6 adh2 adh5 adh4 sga1::(ENO2(long promoter) FBA1 PGI1 TPI1 PGK1 PFK1 PFK2 HXK2 TDH3 PGK1 GPM1 AmdSYM PYK1 ADH1) pyk1 pgi1 tpi1 tdh3 pfk2::(pTEFCAS9 nat) pgk1 gpm1 fba1 hxk2 pfk1 adh1 pdc1 eno2 can1::(HsALDOA$_{BA}$HsGPI$_{BI}$HsPGK1$_{BG}$HsPFKM$_{BH}$HsPKM1$_{BJ}$HsGAPDH$_{BE}$HsENO3$_{BF}$ScADH1$_{BB}$HsHK4 $_{BC}$ScPDC1$_{BD}$HsPGAM2$_{BK}$HsTPI1) | Integration human glycolysis |
| IMX1666 | MATa ura3-52 his3-1 leu2-3,112 MAL2-8c SUC2 glk1::HIS5 hxk1::LEU2 tdh1 tdh2 gpm2::LoxP gpm3 eno1 pyk2 pdc5 pdc6 adh2 adh5 adh4 sga1 pyk1 pgi1 tpi1 tdh3 pfk2::(pTEFCAS9 nat) pgk1 gpm1 fba1 hxk2 pfk1 adh1 pdc1 eno2 can1::( HsALDOA$_{BA}$HsGPI$_{BI}$HsPGK1$_{BG}$HsPFKM$_{BH}$ HsPKM1$_{BJ}$ HsGAPDH$_{BE}$HsENO3$_{BF}$ScADH1$_{BB}$ HsHK4 $_{BC}$ScPDC1$_{BD}$HsPGAM2$_{BK}$HsTPI1) pUDE342 | Removal yeast glycolysis |
| IMX1668 | MATa ura3-52 his3-1 leu2-3,112 MAL2-8c SUC2 glk1::HIS5 hxk1::LEU2 tdh1 tdh2 gpm2::LoxP gpm3 eno1 pyk2 pdc5 pdc6 adh2 adh5 adh4 sga1 pyk1 pgi1 tpi1 tdh3 pfk2::(pTEFCAS9 nat) pgk1 gpm1 fba1 hxk2 pfk1 adh1 pdc1 eno2 can1::( HsALDOA$_{BA}$HsGPI$_{BI}$HsPGK1$_{BG}$ HsPFKM$_{BH}$HsPKM1$_{BJ}$ HsGAPDH$_{BE}$HsENO3$_{BF}$ScADH1$_{BB}$HsHK4 $_{BC}$ScPDC1$_{BD}$HsPGAM2$_{BK}$HsTPI1) | Removal plasmid |
| IMX1785 | MATa ura3-52 his3-1 leu2-3,112 MAL2-8c SUC2 glk1::HIS5 hxk1::LEU2 tdh1 tdh2 gpm2::LoxP gpm3 eno1 pyk2 pdc5 pdc6 adh2 adh5 adh4 sga1 pyk1 pgi1 tpi1 tdh3 pfk2::(pTEFCAS9 nat) pgk1 gpm1 fba1 hxk2 pfk1 adh1 pdc1 eno2 can1::( HsALDOA$_{BA}$HsGPI$_{BI}$HsPGK1$_{BG}$ HsPFKM$_{BH}$HsPKM1$_{BJ}$ HsGAPDH$_{BE}$HsENO3$_{BF}$ScADH1$_{BB}$HsHK2*$_{BC}$ScPDC1$_{BD}$HsPGAM2$_{BK}$HsTPI1) | Replace *HK4* with *HK2* |
| IMX1814 | MATa ura3-52 his3-1 leu2-3,112 MAL2-8c SUC2 glk1::HIS5 hxk1::LEU2 tdh1::URA3 tdh2 gpm2::LoxP gpm3 eno1 pyk2 pdc5 pdc6 adh2 adh5 adh4 sga1 pyk1 pgi1 tpi1 tdh3 pfk2::(pTEFCAS9 nat) pgk1 gpm1 fba1 hxk2 pfk1 adh1 pdc1 eno2 | Integration *ScURA3* in *tdh1* |

| | | |
|---|---|---|
| | can1::( HsALDOA$_{BA}$HsGPI$_{BI}$HsPGK1$_{BG}$ HsPFKM$_{BH}$ HsPKM1$_{BJ}$ HsGAPDH$_{BE}$HsENO3$_{BF}$ScADH1$_{BB}$HsHK4$_{BC}$ScPDC1$_{BD}$HsPGAM2$_{BK}$HsTPI1) | |
| IMX1844 | MATa ura3-52 his3-1 leu2-3,112 MAL2-8c SUC2 glk1::HIS5 hxk1::LEU2 tdh1::URA3 tdh2 gpm2::LoxP gpm3 eno1 pyk2 pdc5 pdc6 adh2 adh5 adh4 sga1 pyk1 pgi1 tpi1 tdh3 pfk2::(pTEFCAS9 nat) pgk1 gpm1 fba1 hxk2 pfk1 adh1 pdc1 eno2 can1::( HsALDOA$_{BA}$HsGPI$_{BI}$HsPGK1$_{BG}$ HsPFKM$_{BH}$ HsPKM1$_{BJ}$ HsGAPDH$_{BE}$HsENO3$_{BF}$ScADH1$_{BB}$HsHK2*$_{BC}$ScPDC1$_{BD}$HsPGAM2$_{BK}$HsTPI1) | Integration *ScURA3* in *tdh1* |
| IMX2418 | MATa ura3-52 his3-1 leu2-3,112 MAL2-8c SUC2 glk1::HIS5 hxk1::LEU2 tdh1::URA3 tdh2 gpm2::LoxP gpm3 eno1 pyk2 pdc5 pdc6 adh2 adh5 adh4 sga1 pyk1 pgi1 tpi1 tdh3 pfk2::(pTEFCAS9 nat) pgk1 gpm1 fba1 hxk2 pfk1 adh1 pdc1 eno2 can1::( HsALDOA$_{BA}$HsGPI$_{BI}$HsPGK1$_{BG}$ HsPFKM$_{BH}$ HsPKM1$_{BJ}$ HsGAPDH$_{BE}$HsENO3$_{BF}$ScADH1$_{BB}$HsHK2* $_{BC}$ScPDC1$_{BD}$ HsPGAM2$_{BK}$ HsTPI1) | Replace *HK4* from IMX1814 with *HK2* selection on glucose |
| IMX2496 | MATa ura3-52 his3-1 leu2-3,112 MAL2-8c SUC2 glk1::HIS5 hxk1::LEU2 tdh1::URA3 tdh2 gpm2::LoxP gpm3 eno1 pyk2 pdc5 pdc6 adh2 adh5 adh4 sga1 pyk1 pgi1 tpi1 tdh3 pfk2::(pTEFCAS9 nat) pgk1 gpm1 fba1 hxk2 pfk1 adh1 pdc1 eno2 can1::( HsALDOA$_{BA}$HsGPI$_{BI}$HsPGK1$_{BG}$ HsPFKM$_{BH}$ HsPKM1$_{BJ}$ HsGAPDH$_{BE}$HsENO3$_{BF}$ScADH1$_{BB}$HsHK2$_{BC}$ScPDC1$_{BD}$HsPGAM2$_{BK}$HsTPI1) | Replace *HK4* from IMX1814 with *HK2* |
| IMX2005 | MATa ura3-52 his3-1 leu2-3,112 MAL2-8c SUC2 glk1::HIS5 hxk1::LEU2 tdh1::URA3 tdh2 gpm2::LoxP gpm3 eno1 pyk2 pdc5 pdc6 adh2 adh5 adh4 sga1 pyk1 pgi1 tpi1 tdh3 pfk2::(pTEFCAS9 nat) pgk1 gpm1 fba1 hxk2 pfk1 adh1 pdc1 eno2 can1::(HsALDOA$_{BA}$HsGPI$_{BI}$HsPGK1$_{BG}$ HsPFKM$_{BH}$ HsPKM1$_{BJ}$ HsGAPDH$_{BE}$ HsENO3$_{BF}$ScADH1$_{BB}$HsHK2*$_{BC}$ScPDC1$_{BD}$HsPGAM2$_{BK}$HsTPI1), X2::(ALDOA$_{BA}$HK2$_{BD}$PGAM2) | Integration extra copies in X2 locus |

| | | |
|---|---|---|
| IMX2006 | MATa ura3-52 his3-1 leu2-3,112 MAL2-8c SUC2 glk1::HIS5 hxk1::LEU2 tdh1::URA3 tdh2 gpm2::LoxP gpm3 eno1 pyk2 pdc5 pdc6 adh2 adh5 adh4 sga1 pyk1 pgi1 tpi1 tdh3 pfk2::(pTEFCAS9 nat) pgk1 gpm1 fba1 hxk2 pfk1 adh1 pdc1 eno2 can1::(HsALDOA$_{BA}$HsGPI$_{BI}$HsPGK1$_{BG}$ HsPFKM$_{BH}$ HsPKM1$_{BJ}$ HsGAPDH$_{BE}$ HsENO3$_{BF}$ScADH1$_{BB}$HsHK4$_{BC}$ScPDC1$_{BD}$HsPGAM2$_{BK}$HsTPI1), X2::(ALDOA$_{BA}$HK4$_{BD}$PGAM2) | Integration extra copies in X2 locus |

Table S7D: Evolved strains and reverse engineered *STT4* strains

| Strain | Genotype | Description |
|---|---|---|
| IMS0987 | Single colony isolate of evolved IMX1844 (Hsgly var. HK2) | |
| IMS0989 | Single colony isolate of evolved IMX1844 (Hsgly var. HK2) | |
| IMS0993 | Single colony isolate of evolved IMX1844 (Hsgly var. HK2) | |
| IMS0990 | Single colony isolate of evolved IMX1814 (Hsgly var. HK4) | |
| IMS0991 | Single colony isolate of evolved IMX1814 (Hsgly var. HK4) | |
| IMS0992 | Single colony isolate of evolved IMX1814 (Hsgly var. HK4) | |

| | | |
|---|---|---|
| IMX2369 | MATa ura3-52 his3-1 leu2-3,112 MAL2-8c SUC2 glk1::Sphis5 hxk1::KlLEU2 tdh1:URA3 tdh2 gpm2 gpm3 eno1 pyk2 pdc5 pdc6 adh2 adh5 adh4 sga1::( FBA1$_{H}$TPI1$_{P}$ PGK1$_{Q}$ ADH1$_{N}$ PYK1$_{O}$ TDH3$_{A}$ ENO2$_{B}$ HXK2$_{C}$ PGI1$_{D}$ PFK1J PFK2$_{JK}$ GPM1$_{L}$ PDC1) pyk1 pgi1 tpi1 tdh3 pfk2::(pTEF-cas9-tCYC1 natNT1) pgk1 gpm1 fba1 hxk2 pfk1 adh1 pdc1 eno2, STT4 (G1766R) | Host strain: IMX1822 |
| IMX2370 | MATa ura3-52 his3-1 leu2-3,112 MAL2-8c SUC2 glk1::Sphis5 hxk1::KlLEU2 tdh1:URA3 tdh2 gpm2 gpm3 eno1 pyk2 pdc5 pdc6 adh2 adh5 adh4 sga1::(FBA1$_{H}$TPI1$_{P}$ PGK1$_{Q}$ ADH1$_{N}$ PYK1$_{O}$ TDH3$_{A}$ ENO2$_{B}$ HXK2$_{C}$ PGI1$_{D}$ PFK1J PFK2$_{JK}$ GPM1$_{L}$ PDC1) pyk1 pgi1 tpi1 tdh3 pfk2::(pTEF-cas9-tCYC1 natNT1) pgk1 gpm1 fba1 hxk2 pfk1 adh1 pdc1 eno2, STT4 (F1775I) | Host strain: IMX1822 |
| IMX2371 | MATa ura3-52 his3-1 leu2-3,112 MAL2-8c SUC2 glk1::HIS5, hxk1::LEU2, tdh1, tdh2::AB, gpm2::LoxP, gpm3, eno1, pyk2, pdc5, pdc6, adh2, adh5, adh4, sga1, pyk1 pgi1 tpi1 tdh3 pfk2::(pTEFCAS9 nat) pgk1 gpm1 fba1 hxk2 pfk1 adh, pdc1, eno2, can1::(HsALDOA$_{BA}$HsGPI$_{BI}$HsPGK1$_{BG}$ HsPFKM$_{BH}$ HsPKM1$_{BJ}$ HsGAPDH$_{BE}$HsENO3$_{BF}$ScADH1$_{BB}$HsHK4$_{BC}$ScPDC1$_{BD}$HsPGAM2$_{BK}$HsTPI1), STT4 (G1766R) | Host strain: IMX1814 |
| IMX2372 | MATa ura3-52 his3-1 leu2-3,112 MAL2-8c SUC2 glk1::HIS5, hxk1::LEU2, tdh1, tdh2::AB, gpm2::LoxP, gpm3, eno1, pyk2, pdc5, pdc6, adh2, adh5, adh4, sga1, pyk1 pgi1 tpi1 tdh3 pfk2::(pTEFCAS9 nat) pgk1 gpm1 fba1 hxk2 pfk1 adh, pdc1, eno2, can1::(HsALDOA$_{BA}$HsGPI$_{BI}$HsPGK1$_{BG}$ HsPFKM$_{BH}$ HsPKM1$_{BJ}$ HsGAPDH$_{BE}$HsENO3$_{BF}$ScADH1$_{BB}$HsHK4$_{BC}$ScPDC1$_{BD}$HsPGAM2$_{BK}$HsTPI1), STT4 (F1775I) | Host strain: IMX1814 |
| IMX2373 | MATa ura3-52 his3-1 leu2-3,112 MAL2-8c SUC2 glk1::HIS5, hxk1::LEU2, tdh1:URA3sc, tdh2::AB, gpm2::LoxP, gpm3, eno1, pyk2, pdc5, pdc6, adh2, adh5, adh4, sga1, pyk1 pgi1 tpi1 tdh3 pfk2::(pTEFCAS9 nat) pgk1 gpm1 fba1 hxk2 pfk1 adh, pdc1, eno2, can1::(HsALDOA$_{BA}$HsGPI$_{BI}$HsPGK1$_{BG}$ HsPFKM$_{BH}$ HsPKM1$_{BJ}$ HsGAPDH$_{BE}$HsENO3$_{BF}$ScADH1$_{BB}$HsHK2*$_{BC}$ScPDC1$_{BD}$HsPGAM2$_{BK}$HsTPI1), STT4 (G1766R) | Host strain: IMX1844 |

| | | |
|---|---|---|
| IMX2374 | MATa ura3-52 his3-1 leu2-3,112 MAL2-8c SUC2 glk1::HIS5, hxk1::LEU2, tdh1:URA3sc, tdh2::AB, gpm2::LoxP, gpm3, eno1, pyk2, pdc5, pdc6, adh2, adh5, adh4, sga1, pyk1 pgi1 tpi1 tdh3 pfk2::(pTEFCAS9 nat) pgk1 gpm1 fba1 hxk2 pfk1 adh, pdc1, eno2, can1::( $HsALDOA_{BA}HsGPI_{BI}HsPGK1_{BG}$ $HsPFKM_{BH}$ $HsPKM1_{BJ}$ $HsGAPDH_{BE}HsENO3_{BF}ScADH1_{BB}HsHK2*_{BC}ScPDC1_{BD}HsPGAM2_{BK}$HsTPI1), STT4 (F1775I) | Host strain: IMX1844 |

Table S7E: Control strains and intermediate strains

| Strain | Genotype | Source |
|---|---|---|
| CEN.PK113-7D | MATa URA3 HIS3 LEU2 TRP1 MAL2-8c SUC2 | [10, 11] |
| CEN.PK122 | MATa/Matα | [10, 11] |
| CEN.PK102-12A | MATa ura3-52 his3-D1 leu2-3,112 TRP1 MAL2-8c SUC2 | [10, 11] |
| IMX370 | MATa ura3-52 his3-1 leu2-3,112 MAL2-8c SUC2 glk1::SpHis5, hxk1::KlLEU2, tdh1, tdh2, gpm2, gpm3, eno1, pyk2, pdc5, pdc6, adh2, adh5, adh4 | [12] |
| IMX372 | MATa ura3-52 his3-1 leu2-3,112 MAL2-8c SUC2 glk1::SpHIS5, hxk1::KlLEU2, tdh1::URA3, tdh2, gpm2, gpm3, eno1, pyk2, pdc5, pdc6, adh2, adh5, adh4 | [12] |

| | | |
|---|---|---|
| IMX1076 | MATa ura3-52 his3-1 leu2-3,112 MAL2-8c SUC2 glk1::SpHis5, hxk1::KlLEU2, tdh1, tdh2, gpm2, gpm3, eno1, pyk2, pdc5, pdc6, adh2, adh5, adh4, sga1::(CAS9, NatNT) | This study |
| IMX589 | MATa ura3-52 his3-1 leu2-3,112 MAL2-8c SUC2 glk1::Sphis5 hxk1::KlLEU2 tdh1 tdh2 gpm2 gpm3 eno1 pyk2 pdc5 pdc6 adh2 adh5 adh4 sga1::( $FBA1_{H}TPI1_{P}PGK1_{Q}ADH1_{N}PYK1_{O}$ $TDH3_{A}ENO2_{B}HXK2_{C}PGI1_{D}PFK1_{J}PFK2_{K}AmdSYM_{L}GPM1_{M}$PDC1) pyk1 pgi1 tpi1 tdh3 pfk2::(pTEF-cas9-tCYC1 natNT1) pgk1 gpm1 fba1 hxk2 pfk1 adh1 pdc1 eno2 | [9] |
| IMX1769 | MATa ura3-52 his3-1 leu2-3,112 MAL2-8c SUC2 glk1::Sphis5 hxk1::KlLEU2 tdh1 tdh2 gpm2 gpm3 eno1 pyk2 pdc5 pdc6 adh2 adh5 adh4 sga1::( $FBA1_{H}TPI1_{P}$ $PGK1_{Q}$ $ADH1_{N}$ $PYK1_{O}$ $TDH3_{A}$ $ENO2_{B}$ $HXK2_{C}$ $PGI1_{D}$ PFK1J $PFK2_{JK}$ $GPM1_{L}$ PDC1) pyk1 pgi1 tpi1 tdh3 pfk2::(pTEF-cas9-tCYC1 natNT1) pgk1 gpm1 fba1 hxk2 pfk1 adh1 pdc1 eno2 | This study |
| IMX1822 | MATa ura3-52 his3-1 leu2-3,112 MAL2-8c SUC2 glk1::Sphis5 hxk1::KlLEU2 tdh1::ScURA3 tdh2 gpm2 gpm3 eno1 pyk2 pdc5 pdc6 adh2 adh5 adh4 sga1::( $FBA1_{H}TPI1_{P}$ $PGK1_{Q}$ $ADH1_{N}$ $PYK1_{O}$ $TDH3_{A}$ $ENO2_{B}$ $HXK2_{C}$ $PGI1_{D}$ PFK1J $PFK2_{JK}$ $GPM1_{L}$ PDC1) pyk1 pgi1 tpi1 tdh3 pfk2::(pTEF-cas9-tCYC1 natNT1) pgk1 gpm1 fba1 hxk2 pfk1 adh1 pdc1 eno2 | This study |
| IMX075 | MATa ura3-52 his3-1 leu2-3,112 MAL2-8c SUC2 hxk1::LEU2 | This study |
| IMS0336 | MATa ura3-52 his3-1 leu2-3,112 MAL2-8c SUC2 hxk1::LoxP | This study |
| IMX165 | MATa ura3-52 his3-1 leu2-3,112 MAL2-8c SUC2 hxk1::LoxP hxk2::KanMX | This study |
| IMX2014 | MATa ura3-52 his3-1 leu2-3,112 MAL2-8c SUC2 glk1::SpHis5, hxk1::KlLEU2, tdh1, tdh2, gpm2, gpm3, eno1, pyk2, pdc5, pdc6, adh2, adh5, adh4, sga1::(CAS9, NatNT), ScPDC1p-HXK2 | This study |

| | | |
|---|---|---|
| IMX2015 | MATa URA3, his3-1 leu2-3,112 MAL2-8c SUC2 glk1::SpHis5, hxk1::KlLEU2, tdh1, tdh2, gpm2, gpm3, eno1, pyk2, pdc5, pdc6, adh2, adh5, adh4, sga1::(CAS9, NatNT), ScPDC1p-HXK2 | This study |

2

#### Table S8 – List of primers used in this study

Table S8A: Primers amplification of yeast promoters and terminators with yeast toolkit flanks

| Name | Sequence |
|---|---|
| 9755 pPDC1 fw Ytk | AAGCATCGTCTCATCGGTCTCAAACGCATGCGACTGGGTGAGCATATG |
| 9756 pPDC1 rev Ytk | TTATGCCGTCTCAGGTCTCACATATTTGATTGATTTGACTGTGTTATTTTGCG |
| 10755 pTEF1 fw Ytk | AAGCATCGTCTCATCGGTCTCAAACGCGGGATAATTAAGACGACAAGAAG |
| 10756 pTEF1 rv Ytk | TTATGCCGTCTCAGGTCTCACATATTTGTAATTAAAACTTAGATTAGATTGCTATG |
| 9419 pFBA1 fw Ytk | AAGCATCGTCTCATCGGTCTCAAACGCAATACCAGCCTTCCAACTTC |
| 9420 pFBA1 rv Ytk | TTATGCCGTCTCAGGTCTCACATATTTGAATATGTATTACTTGGTTATGG |
| 9423 pTPI1 fw Ytk | AAGCATCGTCTCATCGGTCTCAAACGACCCAGAGATGTTGTTGTCC |
| 9424 pTPI1 rv Ytk | TTATGCCGTCTCAGGTCTCACATATTTTAGTTTATGTATGTGTTTTTTGTAG |
| 10753 pTDH3 fw Ytk | AAGCATCGTCTCATCGGTCTCAAACGCGAATATATACTAGCGTTGAATGTTAG |
| 10754 pTDH3 rv Ytk | TTATGCCGTCTCAGGTCTCACATATTTGTTTGTTTATGTGTGTTTATTCG |
| 9421 pPGK1 fw Ytk | AAGCATCGTCTCATCGGTCTCAAACGTATTTTAGATTCCTGACTTCAACTC |

| | |
|---|---|
| 9422 pPGK1 rv Ytk | TTATGCCGTCTCAGGTCTCACATATGTTTTATATTTGTTGTAAAAAGTAGATAATTAC |
| 9757 pGPM1 fw Ytk | AAGCATCGTCTCATCGGTCTCAAACGGTGATACTTTGACAGGAGC |
| 9758 pGPM1 rv Ytk | TTATGCCGTCTCAGGTCTCACATATATTGTAATATGTGTGTTTGTTTGG |
| 9739 pENO2 fw Ytk | AAGCATCGTCTCATCGGTCTCAAACGGGATGATGAAAACACTAAACGAAG |
| 9740 pENO2 rv Ytk | TTATGCCGTCTCAGGTCTCACATATATTATTGTATGTTATAGTATTAGTTGCTTGG |
| 10608 pPYK1 fw Ytk | AAGCATCGTCTCATCGGTCTCAAACGCCCTGGTCAAACTTCAGAAC |
| 10609 pPYK1 rv Ytk | TTATGCCGTCTCAGGTCTCACATATGTGATGATGTTTTATTTGTTTTGATTG |
| 10773 tPDC1 fw Ytk | AAGCATCGTCTCATCGGTCTCAATCCGCGATTTAATCTCTAATTATTAGTTAAAG |
| 10774 tPDC1 rv Ytk | TTATGCCGTCTCAGGTCTCACAGCCAGTGTTCCTTAATCAAGGATACC |
| 10884 tTEF2 fw Ytk | AAGCATCGTCTCATCGGTCTCAATCCGAGTAATAATTATTGCTTCCATATAATATTTTTATATAC |
| 10885 tTEF2 fw Ytk | TTATGCCGTCTCAGGTCTCACAGCAGGAAACGTAAATTACAAGGTATATAC |
| 10767 tTEF1 fw Ytk | AAGCATCGTCTCATCGGTCTCAATCCGGAGATTGATAAGACTTTTCTAGTTG |
| 10768 tTEF1 rv Ytk | TTATGCCGTCTCAGGTCTCACAGCGGTATCACCATAGATTTCGAAAC |

| | |
|---|---|
| 10757 tFBA1 fw Ytk | AAGCATCGTCTCATCGGTCTCAATCCGTTAATTCAAATTAATTGATATAGTTTTTTAATG |
| 10758 tFBA1 rv Ytk | TTATGCCGTCTCAGGTCTCACAGCCGCGAACTCCAAAATGAGC |
| 10765 tTPI1 fw Ytk | AAGCATCGTCTCATCGGTCTCAATCCGATTAATATAATTATATAAAAATATTATCTTCTTTTC |
| 10766 tTPI1 rv Ytk | TTATGCCGTCTCAGGTCTCACAGCCGGTACACTTCTGAGTAAC |
| 10761 tTDH3 fw Ytk | AAGCATCGTCTCATCGGTCTCAATCCGTGAATTTACTTTAAATCTTGCATTTAAATAAATTTTC |
| 10762 tTDH3 rv Ytk | TTATGCCGTCTCAGGTCTCACAGCGTAACTTCAGAATCGTTATCCTGG |
| 10763 tPGK1 fw Ytk | AAGCATCGTCTCATCGGTCTCAATCCATTGAATTGAATTGAAATCGATAG |
| 10764 tPGK1 rv Ytk | TTATGCCGTCTCAGGTCTCACAGCCGAAATAATATCCTTCTCGAAAG |
| 10759 tGPM1 fw Ytk | AAGCATCGTCTCATCGGTCTCAATCCGTCTGAAGAATGAATGATTTGATG |
| 10760 tGPM1 rv Ytk | TTATGCCGTCTCAGGTCTCACAGCCATTAAACTACGATGTAAACATC |
| 10886 tPYK1 fw Ytk | AAGCATCGTCTCATCGGTCTCAATCCAAAAAGAATCATGATTGAATGAAGATATT |
| 10887 tPYK1 rev Ytk | TTATGCCGTCTCAGGTCTCACAGCGTATCCTTTCGCCATCCTG |

Table S8B: Diagnostic primers

| Name | Sequence |
|---|---|
| Primers to confirm integration human genes | |
| 9442 URA3 5' barcode Ytkit | GTAATGTTATCCATGTGGGC |
| 7653_ URA3 upstream | ATTCCAACTAATGAGATGGAATCG |
| 4728_ URA3 downstream | CCAGCCCATATCCAACTTCC |
| 9441 URA3 3' barcode Ytkit | AGAGCACTTGAATCCACTGC |
| Primers to confirm yeast gene deletion | |
| 2798 ENO2 fw | TGAAGTGTGATACCAAGTCAGC |
| 5237 ENO2 rev | AATAGACAGCACGAGTCTTTG |
| 1152 PYK1 fw | TGGCGTGTGATGTCTGTATCTG |
| 4667 PYK1 rev | CCTTGAGGGAAGATTATCTTGCG |
| 5134 TDH3 fw | CCAAAAATAGCCGAGCAAGCTC |
| 4788 TDH3 rev | AACGCTAAGAGTAACTTCAGAATCG |
| 7414 PGI1 fw | AATGTAGCGACACCACTTCC |
| 5004 PGI1 rev | GTAGATTGCACCATCTGAAGAGGC |
| 10590 PGK1 fw | ACTGTAATTGCTTTTAGTTGTG |
| 4698 PGK1 rev | TACGCTGAACCCGAACATAG |
| 12330 GPM1 fw | GCAGACGACAGATCTAAATGAC |
| 12331 GPM1 rev | GCCACCGTACATTTAATATGTC |

| | |
|---|---|
| 11067 FBA1 fw | AACTACACGGAAGCTCTAAAGATG |
| 5024 FBA1 rev | CCCTCTTATTTATTAGCATTGTCTTCCG |
| 3481 HXK2 fw | GCCTAGCGTCTGGGATTTATTC |
| 3070 HXK2 rev | AGTGCTTCCGTTCGTTCCAG |
| 3514 TPI1 fw | CTGACAGGTGGTTTGTTACG |
| 8726 TPI1 rev | TCAGCCATTGAGCAGAGAAC |
| 4925 PFK1 fw | AATTTTTACCCTGATCTAACTAACTTTGG |
| 4924 PFK1 rev | GTAGACCGATGACAATACGACTAC |
| 4777 PFK2 fw | CGTGAGCCTTAACCAATGAG |
| 4776 PFK2 rev | CTCCGTTCTTCGTGATAAGTTC |

Table S8C: Primers to amplify glycoblocks

| Fragment | Name | Sequence |
|---|---|---|
| HsALDOA$_{can1/BA}$ | 12952 tFBA1 + can1 | GTTTTTAATCTGTCGTCAATCGAAAGTTTATTTCAGAGTTCTTCAGACTTCTTAACTCCTGTGCATGACAAAAGATGAGCTAGG |
| | 12446 pFBA+ BA | TAAGTCTCTTGACATCTCGGAACATATCCACTCAGCGGTGTATCATTCTGTGGTCGGCGCCATGCCTCCAACGGCTACTATC |
| HsGPI$_{BA/BI}$ | 12447 pTEF2 + BA | GCGCCGACCACAGAATGATACACCGCTGAGTGGATATGTTCCGAGATGTCAAGAGACTTAAACGTTGATAGGTCAAGATCAATG |
| | 12448 tTEF2 + BI | TCTGTCAGTTGGTTAAGCGCCGCTACGATTACTACACATGCCACAGACTGATCTACAATGAATTACAAGGTATATACATACGCTGACATG |
| HsPGK1$_{BI/BG}$ | 12474 tPGK1 + BI | CATTGTAGATCAGTCTGTGGCATGTGTAGTAATCGTAGCGGCGCTTAACCAACTGACAGATGGCAGCCGAAATAATATCCTTC |
| | 12475 pPGK1 + BG | GAGGCTTCACAGTGCTTTATTAGTATGATTGCCTAGCTGGTATATGTGTTCCTGGAGCGCTTCCTGACTTCAACTCAAGACGC |
| HsPFKM$_{BG/BH}$ | 12453 pTEF1 + BG | GCGCTCCAGGAACACATATACCAGCTAGGCAATCATACTAATAAAGCACTGTGAAGCCTCCGGGATAATTAAGACGACAAGAAG |
| | 12454 tTEF1 + BH | AGGATCGCTCGCGTACTCATGCATTCTCCCACATATTGAGGCCCTGATTCCATGCAATGTGGCAGCGGTATCACCATAG |
| HsPKM1$_{BH/BJ}$ | 12455 pPYK1 + BH | ACATTGCATGGAATCAGGGCCTCAATATGTGGGAGAATGCATGAGTACGCGAGCGATCCTCCTGGTCAAACTTCAGAACTAAG |
| | 12456 tPYK1 + BJ | GGCGCACATGGTATATTATGATCGGAGATGCGGCAACATAGCTGGGTGTGATCCTCTCTACGTATCCTTTCGCCATCCTG |

| | | |
|---|---|---|
| HsGAPDH$_{BJ/BE}$ | 12457 tTDH3 + BJ | TAGAGAGGATCACACCCAGCTATGTTGCCGCATCTCCGATCA TAATATACCATGTGCGCCCAGAATCGTTATCCTGGCGG |
| | 12458 pTDH3 + BE | TCAATCATTCGTTCTCGCAGATCTACAATCGTCCTGAGCTCTG TGAGTGATGTACGCTCCTACTAGCGTTGAATGTTAGCGTC |
| HsENO3$_{BE/BF}$ | 12459 pENO2 + BE | GGAGCGTACATCACTCACAGAGCTCAGGACGATTGTAGATC TGCGAGAACGAATGATTGATGATGAAAACACTAAACGAAGG |
| | 12460 tENO2 + BF | GCGCGACGTGTCTCGTATATTAGTGAAGTTGGATCTGTCCAT GAATCCTCGGCTCTGGTGTATTTTTCAAACTGCAAATTCAAG |
| ScADH1$_{BF/BB}$ | 12461 tADH1 + BF | CACCAGAGCCGAGGATTCATGGACAGATCCAACTTCACTAAT ATACGAGACACGTCGCGCATGCCGGTAGAGGTGTGGTC |
| | 12462 pADH1 + BB | GCAACGCATTCCATACATGATGCGTTGCTTGGTGTCCACAGC CGTACTTGAGAAGCTCTGAGTCCAATGCTAGTAGAGAAGGG |
| HsHK4$_{BB/BC}$ | 12463 pHXK2 + BB | CAGAGCTTCTCAAGTACGGCTGTGGACACCAAGCAACGCAT CATGTATGGAATGCGTTGCGCTGGTAAAGTACAGCTACATTC |
| | 12464 tHXK2 + BC | CTAGGCTCTGCTGCATGTCAGTGATTTCTATTAGGCAGCGCT TACCCATGATTAGCGCAGACTTGAACAATAAATACGAAATCC |
| ScPDC1$_{BC/BD}$ | 12465 tPDC1 + BC | CTGCGCTAATCATGGGTAAGCGCTGCCTAATAGAAATCACTG ACATGCAGCAGAGCCTAGTGTTCCTTAATCAAGGATACCTC |
| | 12466 pPDC1 + BD | AGTCACGCTGAGTCCATGCTGACCATGATTCACACTCAGTGC CGATAATTCCATAGTCTGCGACTGGGTGAGCATATGTTC |
| HsPGAM2$_{BD/BK}$ | 12467 pGPM1 + BD | CAGACTATGGAATTATCGGCACTGAGTGTGAATCATGGTCA GCATGGACTCAGCGTGACTGATACTTTGACAGGAGCTATATC |
| | 12468 tGPM1 + BK | GAGCATACTGTCCTATCATGTCGACTCTTGTCACATCTGACG CCTCTCTGCGATAGGATTTGCTATAACATGTCATGTCACC |
| HsTPI$_{BK/can1}$ | 12469 tTPI1 + BK | AATCCTATCGCAGAGAGGCGTCAGATGTGACAAGAGTCGAC ATGATAGGACAGTATGCTCTGAGTAACCCATATAGAGATCG |

| | | |
|---|---|---|
| | 12470 pTPI + can1 | GTGTATGACTTATGAGGGTGAGAATGCGAAATGGCGTGGA AATGTGATCAAAGGTAATAACCAGAGATGTTGTTGTCCTAG |
| HsHK2 | 13506 HK2 + pHXK2 flank | CTTTGAAAAGATTGTAGGAATATAATTCTCCACACATAATAA GTACGTTAATTAAATAAAATGATCGCCTCTCATTTGTTG |
| | 13507 HK2 + tHXK2 flank | GTTCACATAATTAAAAAAAGGGCACCTTCTTGTTGTTCAAAC TTAATTTACAAATTAAGTTCATCTTTGACCAGCTTCTCT |
| Overexpression | | |
| HsALDOA$_{X2\ flank/BA}$ | 12446 pFBA1+ BA | TAAGTCTCTTGACATCTCGGAACATATCCACTCAGCGGTGTA TCATTCTGTGGTCGGCGCCATGCCTCCAACGGCTACTATC |
| | 12650 tFBA1 + X2 flank | GCTGAAGATTTATCATACTATTCCTCCGCTCGTTTCTTTTTTCA GTGAGGTGTGTCGTGAGTGCATGACAAAAGATGAGCTAGG |
| HsHK2 and HsHK4 $_{BA/BD}$ | 14540 pPDC1 + BA | GCGCCGACCACAGAATGATACACCGCTGAGTGGATATGTTC CGAGATGTCAAGAGACTTAGCGACTGGGTGAGCATATG |
| | 14541 tPDC1 + BD | AGTCACGCTGAGTCCATGCTGACCATGATTCACACTCAGTGC CGATAATTCCATAGTCTGCTCATTGGCAGCCAGTGTTC |
| HsPGAM2$_{BD/X2\ flank}$ | 12467 pGPM1 + BD | CAGACTATGGAATTATCGGCACTGAGTGTGAATCATGGTCA GCATGGACTCAGCGTGACTGATACTTTGACAGGAGCTATATC |
| | 14542_tGPM1 + X2 flank | ATTCTCGCCAAGGCATTACCATCCCATGTAAGAACGGAATAA AACAGCATTCGAAGGTTATTGCTATAACATGTCATGTCACC |

Table S8D: gRNA oligos and primers for backbone amplification

| Target | Name | Sequence |
|---|---|---|
| ScHXK2 | 10205 HXK2 gRNA fw | TGCGCATGTTTCGGCGTTCGAAACTTCTCCGCAGTGAAAG<br>ATAAATGATCGGTAAGTCCGTTGGTATCATGTTTTAGAGCT<br>AGAAATAGCAAGTTAAAATAAGGCTAGTCCGTTATCAAC |
| | 10206 HXK2 gRNA rv | GTTGATAACGGACTAGCCTTATTTTAACTTGCTATTTCTAG<br>CTCTAAAACATGATACCAACGGACTTACCGATCATTTATCT<br>TTCACTGCGGAGAAGTTTCGAACGCCGAAACATGCGCA |
| ScPGI1 | 10080 PGI1 gRNA fw | TGCGCATGTTTCGGCGTTCGAAACTTCTCCGCAGTGAAAG<br>ATAAATGATCCAAAAAATTTATGAATCTCAGTTTTAGAGCT<br>AGAAATAGCAAGTTAAAATAAGGCTAGTCCGTTATCAAC |
| | 10081 PGI1 gRNA rv | GTTGATAACGGACTAGCCTTATTTTAACTTGCTATTTCTAG<br>CTCTAAAACTGAGATTCATAAATTTTTTGGATCATTTATCTT<br>TCACTGCGGAGAAGTTTCGAACGCCGAAACATGCGCA |
| ScPFK1 | 10207 PFK1 gRNA | TGCGCATGTTTCGGCGTTCGAAACTTCTCCGCAGTGAAAG<br>ATAAATGATCCATCATCTCTGAAGCAAGCAGTTTTAGAGCT<br>AGAAATAGCAAGTTAAAATAAG |
| ScPFK2 | 10208 PFK2 gRNA | TGCGCATGTTTCGGCGTTCGAAACTTCTCCGCAGTGAAAG<br>ATAAATGATCCCAGGTCATGTACAACAAGGGTTTTAGAGC<br>TAGAAATAGCAAGTTAAAATAAG |
| ScFBA1 | 12332 FBA1 gRNA | TGCGCATGTTTCGGCGTTCGAAACTTCTCCGCAGTGAAAG<br>ATAAATGATCCAAATCTTAAAGAGAAAGACGTTTTAGAGC<br>TAGAAATAGCAAGTTAAAATAAG |
| ScTPI1 | 10972 TPI1 gRNA fw | TGCGCATGTTTCGGCGTTCGAAACTTCTCCGCAGTGAAAG<br>ATAAATGATCCTTAGACTACTCTGTCTCTTGTTTTAGAGCT<br>AGAAATAGCAAGTTAAAATAAGGCTAGTCCGTTATCAAC |

| | | |
|---|---|---|
| | 10973 TPI1 gRNA rev | GTTGATAACGGACTAGCCTTATTTTAACTTGCTATTTCTAGCTCTAAAACAAGAGACAGAGTAGTCTAAGGATCATTTATCTTTCACTGCGGAGAAGTTTCGAACGCCGAAACATGCGCA |
| ScTDH3 | 10968 TDH3 gRNA fw | TGCGCATGTTTCGGCGTTCGAAACTTCTCCGCAGTGAAAGATAAATGATCTTTGAGTAGCAGTCAAAGAGGTTTTAGAGCTAGAAATAGCAAGTTAAAATAAGGCTAGTCCGTTATCAAC |
| | 10969 TDH3 gRNA rev | GTTGATAACGGACTAGCCTTATTTTAACTTGCTATTTCTAGCTCTAAAACCTCTTTGACTGCTACTCAAAGATCATTTATCTTTCACTGCGGAGAAGTTTCGAACGCCGAAACATGCGCA |
| ScPGK1 | 10970 PGK1 gRNA fw | TGCGCATGTTTCGGCGTTCGAAACTTCTCCGCAGTGAAAGATAAATGATCCAGACACGAATTGAGCTCTTGTTTTAGAGCTAGAAATAGCAAGTTAAAATAAGGCTAGTCCGTTATCAAC |
| | 10971 PGK1 gRNA rev | GTTGATAACGGACTAGCCTTATTTTAACTTGCTATTTCTAGCTCTAAAACAAGAGCTCAATTCGTGTCTGGATCATTTATCTTTCACTGCGGAGAAGTTTCGAACGCCGAAACATGCGCA |
| ScGPM1 | 10976 GPM1 gRNA fw | TGCGCATGTTTCGGCGTTCGAAACTTCTCCGCAGTGAAAGATAAATGATCATTGCCAAGGACTTGTTGAGGTTTTAGAGCTAGAAATAGCAAGTTAAAATAAGGCTAGTCCGTTATCAAC |
| | 10977 GPM1 gRNA fw | GTTGATAACGGACTAGCCTTATTTTAACTTGCTATTTCTAGCTCTAAAACCTCAACAAGTCCTTGGCAATGATCATTTATCTTTCACTGCGGAGAAGTTTCGAACGCCGAAACATGCGCA |
| ScENO2 | 10076 ENO2 gRNA fw | TGCGCATGTTTCGGCGTTCGAAACTTCTCCGCAGTGAAAGATAAATGATCCAAGGCCAACCTAGATGTTAGTTTTAGAGCTAGAAATAGCAAGTTAAAATAAGGCTAGTCCGTTATCAAC |
| | 10077 ENO2 gRNA rv | GTTGATAACGGACTAGCCTTATTTTAACTTGCTATTTCTAGCTCTAAAACTAACATCTAGGTTGGCCTTGGATCATTTATCTTTCACTGCGGAGAAGTTTCGAACGCCGAAACATGCGCA |

| | | |
|---|---|---|
| ScPYK1 | 10974 PYK1 gRNA fw | TGCGCATGTTTCGGCGTTCGAAACTTCTCCGCAGTGAAAGATAAATGATCTATCAACTTCGGTATTGAAAGTTTTAGAGCTAGAAATAGCAAGTTAAAATAAGGCTAGTCCGTTATCAAC |
| | 10975 PYK1 gRNA rev | GTTGATAACGGACTAGCCTTATTTTAACTTGCTATTTCTAGCTCTAAAACTTTCAATACCGAAGTTGATAGATCATTTATCTTTCACTGCGGAGAAGTTTCGAACGCCGAAACATGCGCA |
| HsHK4 | 13696 gRNA HsHK4II fw | TGCGCATGTTTCGGCGTTCGAAACTTCTCCGCAGTGAAAGATAAATGATCATGTGTTCTGCTGGTTTGGCGTTTTAGAGCTAGAAATAGCAAGTTAAAATAAGGCTAGTCCGTTATCAAC |
| | 13697 gRNA HsHK4II rev | GTTGATAACGGACTAGCCTTATTTTAACTTGCTATTTCTAGCTCTAAAACGCCAAACCAGCAGAACACATGATCATTTATCTTTCACTGCGGAGAAGTTTCGAACGCCGAAACATGCGCA |
| ScSTT4 (I) | 16748 gRNA STT4 (IMS0990) fw | TGCGCATGTTTCGGCGTTCGAAACTTCTCCGCAGTGAAAGATAAATGATCAACATTATGTACGATGATCAGTTTTAGAGCTAGAAATAGCAAGTTAAAATAAGGCTAGTCCGTTATCAAC |
| | 16749 gRNA STT4 (MS0990) rev | GTTGATAACGGACTAGCCTTATTTTAACTTGCTATTTCTAGCTCTAAAACTGATCATCGTACATAATGTTGATCATTTATCTTTCACTGCGGAGAAGTTTCGAACGCCGAAACATGCGCA |
| ScSTT4 (II) | 16755 gRNA STT4 (IMS0992) fw | TGCGCATGTTTCGGCGTTCGAAACTTCTCCGCAGTGAAAGATAAATGATCATTGTCTACATATCGATTTTGTTTTAGAGCTAGAAATAGCAAGTTAAAATAAGGCTAGTCCGTTATCAAC |
| | 16756 gRNA STT4 (IMS0992) rev | GTTGATAACGGACTAGCCTTATTTTAACTTGCTATTTCTAGCTCTAAAACAAAATCGATATGTAGACAATGATCATTTATCTTTCACTGCGGAGAAGTTTCGAACGCCGAAACATGCGCA |
| AmdS | 11588 gRNA AmdS fw | TGCGCATGTTTCGGCGTTCGAAACTTCTCCGCAGTGAAAGATAAATGATCATCACATCCGAACATAAACAGTTTTAGAGCTAGAAATAGCAAGTTAAAATAAGGCTAGTCCGTTATCAAC |

| | | |
|---|---|---|
| | 11589 gRNA AmdS rev | GTTGATAACGGACTAGCCTTATTTTAACTTGCTATTTCTAG CTCTAAAACTGTTTATGTTCGGATGTGATGATCATTTATCTT TCACTGCGGAGAAGTTTCGAACGCCGAAACATGCGCA |
| pROS13, pMEL13 and pMEL10 backbone amplification | 6005_p426 CRISP rv | GATCATTTATCTTTCACTGCGGAGAAG |
| | 6006_p426 CRISP fw | GTTTTAGAGCTAGAAATAGCAAGTTAAAATAAGGCTAGTC |

Table S8E: Repair fragments

| Name | Sequence |
|---|---|
| 5888 HXK2 repair oligo fw | TTTCTAATGCCTTTTCCATCATGTTACTACGAGTTTTCTGAACCTCCT CGCACATTGGTAGCTTAATTTTAAATTTTTTTGGTAGTAAAAGATGC TTATATAAGGATTTCGTATTTATTG |
| 5889 HXK2 repair oligo rv | CAATAAATACGAAATCCTTATATAAGCATCTTTTACTACCAAAAAAA TTTAAAATTAAGCTACCAATGTGCGAGGAGGTTCAGAAAACTCGTA GTAACATGATGGAAAAGGCATTAGAAA |
| 10084 PGI1 repair oligo fw | ATACACCGCTATGTATTTCAGGGCACTACTTCTACACATCAACGGTA CTAAACATTTCGCAAAAATTTTAAAAATTAGAGCACCTTGAACTTGC GAAAAAGGTTCTCATCAACTGTTTAA |
| 10085 PGI1 repair oligo rv | TTAAACAGTTGATGAGAACCTTTTTCGCAAGTTCAAGGTGCTCTAAT TTTTAAAATTTTTGCGAAATGTTTAGTACCGTTGATGTGTAGAAGTA GTGCCCTGAAATACATAGCGGTGTAT |
| 10209 PFK2 repair oligo fw | CCAGTCCCGCATACCCCCTTTGCAACGTTAACGTTACCGCTAGCGTT TACCATCTCCACGACTTATGTATACTGGAATATGTGATATAGACGAT TTAAAAGATAATTCCAATAAACGTCC |

| | |
|---|---|
| 10210 PFK2 repair oligo rv | GGACGTTTATTGGAATTATCTTTTAAATCGTCTATATCACATATTCCAGTATACATAAGTCGTGGAGATGGTAAACGCTAGCGGTAACGTTAACGTTGCAAAGGGGGTATGCGGGACTGG |
| 10211 PFK1 repair oligo fw | AATTAATATCTCATTAACAAAGTTATTGTACATAATCCGGTACAATATTCTTCAATGTACGTTTTAGGGTGTGCTTAATCTGCGTTGACAATGGTTCACGAAGACGACATCGGCAACTTT |
| 10212 PFK1 repair oligo rv | AAAGTTGCCGATGTCGTCTTCGTGAACCATTGTCAACGCAGATTAAGCACACCCTAAAACGTACATTGAAGAATATTGTACCGGATTATGTACAATAACTTTGTTAATGAGATATTAATT |
| 12333 FBA1 repair oligo fw | TCTTCTGTTCTTCTTTTTCTTTTGTCATATATAACCATAACCAAGTAATACATATTCAAAGTTAATTCAAATTAATTGATATAGTTTTTTAATGAGTATTGAATCTGTTTAGAAATAATG |
| 12334 FBA1 repair oligo rev | CATTATTTCTAAACAGATTCAATACTCATTAAAAAACTATATCAATTAATTTGAATTAACTTTGAATATGTATTACTTGGTTATGGTTATATATGACAAAAGAAAAAGAAGAACAGAAGA |
| 10980 TPI1 repair oligo fw | TGTTTGTATTCTTTTCTTGCTTAAATCTATAACTACAAAAAACACATACATAAACTAAAAGATTAATATAATTATATAAAAATATTATCTTCTTTTCTTTATATCTAGTGTTATGTAAAA |
| 10981 TPI1 repair oligo rev | TTTTACATAACACTAGATATAAAGAAAAGAAGATAATATTTTTATATAATTATATTAATCTTTTAGTTTATGTATGTGTTTTTTGTAGTTATAGATTTAAGCAAGAAAAGAATACAAACA |
| 10978 TDH3 repair oligo fw | TTTTTTTAGTTTTAAAACACCAAGAACTTAGTTTCGAATAAACACACATAAACAAACAAAGTGAATTTACTTTAAATCTTGCATTTAAATAAATTTTCTTTTTATAGCTTTATGACTTAG |
| 10979 TDH3 repair oligo rv | CTAAGTCATAAAGCTATAAAAAGAAAATTTATTTAAATGCAAGATTTAAAGTAAATTCACTTTGTTTGTTTATGTGTGTTTATTCGAAACTAAGTTCTTGGTGTTTTAAAACTAAAAAAA |

| | |
|---|---|
| 10986 PGK1 repair oligo fw | AAGTTCGTTCGATCGTACTGTTACTCTCTCTCTTTCAAACAGAATTGTCCGAATCGTGTGATTTATATACGTATATATAGACTATTATTTATCTTTTAATGATTATTAAGATTTTTATTA |
| 10987 PGK1 repair oligo rev | TAATAAAAATCTTAATAATCATTAAAAGATAAATAATAGTCTATATATACGTATATAAATCACACGATTCGGACAATTCTGTTTGAAAGAGAGAGAGTAACAGTACGATCGAACGAACTT |
| 10984 GPM1 repair oligo fw | AATTTCAGCTGACAGCGAGTTTCATGATCGTGATGAACAATGGTAACGAGTTGTGGCTGTTTTTTCCCTCCATTTTTCTTACTGAATATATCAATGATATAGACTTGTATAGTTTATTAT |
| 10985 GPM1 repair oligo rev | ATAATAAACTATACAAGTCTATATCATTGATATATTCAGTAAGAAAAATGGAGGGAAAAAACAGCCACAACTCGTTACCATTGTTCATCACGATCATGAAACTCGCTGTCAGCTGAAATT |
| 10086 ENO2 repair oligo fw | TTTTCTTTTCTTAGTTTCTTTCATAACACCAAGCAACTAATACTATAACATACAATAATATTTAACTAAGAATTATTAGTCTTTTCTGCTTATTTTTTCATCATAGTTTAGAACACTTTA |
| 10087 ENO2 repair oligo rv | TAAAGTGTTCTAAACTATGATGAAAAAATAAGCAGAAAAGACTAATAATTCTTAGTTAAATATTATTGTATGTTATAGTATTAGTTGCTTGGTGTTATGAAAGAAACTAAGAAAAGAAAA |
| 10982 PYK1 repair oligo fw | ATTATTCTCTCTTGTTTCTATTTACAAGACACCAATCAAAACAAATAAAACATCATCACAAAAAAGAATCATGATTGAATGAAGATATTATTTTTTTGAATTATATTTTTTAAATTTTAT |
| 10983 PYK1 repair oligo rev | ATAAAATTTAAAAAATATAATTCAAAAAAATAATATCTTCATTCAATCATGATTCTTTTTTGTGATGATGTTTTATTTGTTTTGATTGGTGTCTTGTAAATAGAAACAAGAGAGAATAAT |
| 6075 COUNTER SELECT oligo fw | TTTTTCTCATCTCTTGGCTCTGGATCCGTTATCTGTTCTGTTACACAAGAAATCGTACATACTAGAGCAAGATTTCAAATAAGTAACAGCAGCCATACGTTGAAACTACGGCAAAGGATT |

| | |
|---|---|
| 6076 COUNTER SELECT oligo rv | AATCCTTTGCCGTAGTTTCAACGTATGGCTGCTGTTACTTATTTGAAATCTTGCTCTAGTATGTACGATTTCTTGTGTAACAGAACAGATAACGGATCCAGAGCCAAGAGATGAGAAAAA |
| 16750 STT4 repair IMS990 G1766R fw | CGTAATTTCGTATTTGTTGCAATTCAAGGATAGACATAATGGTAACATTATGTACGATGATCAA**A**GACATTGTCTACATATCGATTTTGGGTTTATTTTTGATATTGTCCCAGGTGGTAT |
| 16751 STT4 repair IMS990 G1766R rv | ATACCACCTGGGACAATATCAAAAATAAACCCAAAATCGATATGTAGACAATGTCTTTGATCATCGTACATAATGTTACCATTATGTCTATCCTTGAATTGCAACAAATACGAAATTACG |
| 16757 STT4 repair IMS992 F1775I fw | GATAGACATAATGGTAACATTATGTACGATGATCAAGGACATTGTCTACATATCGATTTTGG**CA**TTATTTTTGATATTGTCCCAGGTGGTATCAAGTTTGAAGCAGTACCATTCAAGCTG |
| 16758 STT4 repair IMS992 F1775I rv | CAGCTTGAATGGTACTGCTTCAAACTTGATACCACCTGGGACAATATCAAAAATAATGCCAAAATCGATATGTAGACAATGTCCTTGATCATCGTACATAATGTTACCATTATGTCTATC |
| 11590 Repair KL fw | AAGATAGTCGCCGAACTCGCAAGAGTCATTAACACCTCGCAATTGATGGGAAGTCCTCGCATATGACCTGAACCGACGGCAAATGCTCTTCAACTACGGCATACTTGCGGAAGCTACGGC |
| 11591 Repair KL rev | GCCGTAGCTTCCGCAAGTATGCCGTAGTTGAAGAGCATTTGCCGTCGGTTCAGGTCATATGCGAGGACTTCCCATCAATTGCGAGGTGTTAATGACTCTTGCGAGTTCGGCGACTATCTT |

Table S8F: primers for construction of IMX165 and IMX2015

| Fragment | Name | Sequence |
|---|---|---|
| *HXK2* deletion cassette | 2788 HXK2 deletion cassette fw | ATTGTAGGAATATAATTCTCCACACATAATAAGTACGTTAATTAAATAAACAGCTG AAGCTTCGTACGC |
| | 2789 HXK2 deletion cassette rv | TTAAAAAAAGGGCACCTTCTTGTTGTTCAAACTTAATTTACAAATTAAGTGCATAGGCCA CTAGTGGATCTG |
| *HXK1* deletion cassette | 1710 hxk1 deletion cassette fw | AAACTCACCCAAACAACTCAATTAGAATACTGAAAAAATAAGATGATGACAAGAGGGTCGAACTCCAGCTGAAGCTTCGTACGC |
| | 1711_hxk1 deletion cassette rv | AGGGAGGGAAAAACACATTTATATTTCATTACATTTTTTTCATTAGCCTAAGTCGTAATTGAGTCGCATAGGCCACTAGTGGATCTG |
| *pPDC1* amplification | 14670_pPDC1Sc+HXK2flank fw | TTTCTAATGCCTTTTCCATCATGTTACTACGAGTTTTCTGAACCTCCTCGCACATTGGTAGCGACTGGGTGAGCATATG |
| | 14671_pPDC1Sc+HXK2flank rev | TGGCACATCGGCCATGGAACCCTTTCTGGCTTGTGGTTTTTTTGGACCTAAATGAACCATTTTGATTGATTTGACTGTGTTATTTTG |
| Confirmation promoter replacement | 3238 HXK2 outside fw | GCCTTTTCCATCATGTTACTAC |
| | 4834 HXK2 rev | ACCCAATGGAATTGGCTCAG |

Table S8G: Primers for amplification *URA3, HsPKL* and diagnostic primers

| Fragment | Name | Sequence |
|---|---|---|
| *ScURA3* fragment | 11766_URA3 + TDH1 flank fw | GATATTTCACCAACACACACAAAAAACAGTACTTCACTAAATTTACACACAAAACAAAATTGAGTATTTTCAATAAATTTGTAGAGGACT |
| | 11767_URA3 + TDH1 flank rev | CGGTAGTATTTATGTATATTCAAAAAAAAATCATTATCCTCATCAAGATTGCTTTATTTATTGCTTTTGTTCCACTACTTTTTG |
| Confirmation integration *URA3* in *tdh1* | 1989_URA3 outside fw | CCACGTGCAGAACAACATAG |
| | 8306_URA3 rev | TGCTCCTTCCTTCGTTCTTC |
| | 8377_URA3 fw | GGGAATCTCGGTCGTAATG |
| | 2347_URA3 outside rev | GTCACATATTGTGGGTATGTGC |
| Confirmation removal SinLoG cassette | 11898_SeqFW_SGA | CGCGGAAACGGGTATTAGGG |
| | 11899_SeqRV_SGA | CTAGATCCGGTAAGCGACAG |
| Confirmation of replacement *HK4* with *HK2* | 2794 HXK2-FW KO conformation | CACCTTCGCCACTGTCTTATCTAC |
| | 2923 HXK2-RV wca del conf2 | GGGCACCTTCTTGTTGTTCAAACT |
| | 1452 HXK2FW1 | TTCGCCACTGTCTTATCTAC |
| | 13508 HK2 rev | ATCCTTGATTTGCAACTTGTC |
| *HsPKL* gene | 10846 HsPKL gene fw | CCATAGGTCTCATATGGAAGGTCCAGCTGGTTATTTGAG |
| | 10847 HsPKL gene rev | GGCCGGTCTCAGGATTCAGGAGATG |

#### Table S9 – List of plasmids used in this study

Table S9A: Plasmids containing human genes

| Name | Fragment | Source |
|---|---|---|
| pGGKp001 | HK1 | GeneArt |
| pGGKp002 | HK2 | GeneArt |
| pGGKp003 | HK3 | GeneArt |
| pGGKp004 | HK4 | GeneArt |
| pGGKp005 | GPI | GeneArt |
| pGGKp006 | PFKM | GeneArt |
| pGGKp007 | PFKP | GeneArt |
| pGGKp008 | PFKL | GeneArt |
| pGGKp009 | ALDOA | GeneArt |
| pGGKp010 | ALDOB | GeneArt |
| pGGKp011 | ALDOC | GeneArt |
| pGGKp012 | TPI | GeneArt |
| pGGKp013 | GAPDH | GeneArt |
| pGGKp014 | GAPDHS | GeneArt |
| pGGKp015 | PGK1 | GeneArt |
| pGGKp016 | PGK2 | GeneArt |
| pGGKp017 | PGAM1 | GeneArt |
| pGGKp018 | PGAM2 | GeneArt |

| | | |
|---|---|---|
| pGGKp019 | ENO1 | GeneArt |
| pGGKp020 | ENO2 | GeneArt |
| pGGKp021 | ENO3 | GeneArt |
| pGGKp022 | PKM1 | GeneArt |
| pGGKp023 | PKM2 | GeneArt |
| pGGKp024 | PKR | GeneArt |

Table S9B: Plasmids containing yeast promoter or terminator

| Name | Fragment | Source |
|---|---|---|
| pUD565 | Entry vector, CamR | GeneArt |
| pGGKp025 | pPDC1 sc | This study |
| pGGKp026 | pGPM1 sc | This study |
| pGGKp027 | pFBA1 sc | This study |
| pGGKp028 | pENO2 sc | This study |
| pGGKp029 | pADH1 sc | This study |
| pGGKp030 | pTPI1 sc | This study |
| pGGKp031 | pPFK2 sc | This study |
| pGGKp032 | pTEF1 sc | This study |
| pGGKp033 | pPGI1 sc | This study |
| pGGKp034 | pPYK1 sc | This study |
| pGGKp035 | pTDH3 sc | This study |

| | | |
|---|---|---|
| pGGKp036 | pPGK1 sc | This study |
| pGGKp037 | tADH1 sc | This study |
| pGGKp038 | tTEF2 sc | This study |
| pGGKp039 | tTEF1 sc | This study |
| pGGKp040 | tPYK1 sc | This study |
| pGGKp041 | tTDH3 sc | This study |
| pGGKp042 | tTPI1 sc | This study |
| pGGKp043 | tPGK1 sc | This study |
| pGGKp044 | tPGI1 sc | This study |
| pGGKp045 | tPDC1 sc | This study |
| pGGKp046 | tFBA1 sc | This study |
| pGGKp047 | pACT1 sc | This study |
| pGGKp048 | tGPM1 sc | This study |
| pGGKp096 | pHXK2 | [13] |
| pGGKp097 | tHXK2 | [13] |
| pYTK055 | tENO2 | [14] |
| pYTK014 | pTEF2 | [14] |

Table S9C: Plasmids used to construct pGGKd002 and pGGKd003

| Name | Fragment | Source |
|---|---|---|
| pYTK002 | ConLS | [14] |
| pYTK047 | GFP dropout | [14] |
| pYTK067 | ConR1 | [14] |
| pYTK074 | URA3 | [14] |
| pYTK086 | URA3 3' Homology | [14] |
| pYTK090 | KanR-ColE1 | [14] |
| pYTK092 | URA3 5' Homology | [14] |
| pGGKd002 | GFP dropout integration plasmid made from pYTK002,47,67,74,86,90,92 | This study |
| pGGKd003 | GFP dropout plasmid made from pYTK002,47,67,74,82,84 | This study |

Table S9D: Integration plasmids containing human transcriptional unit

| Name | Construct | Source |
|---|---|---|
| pUDI133 | ScPDC1p-HK1-ScPDC1t | This study |
| pUDI134 | ScPDC1p-HK2-ScPDC1t | This study |
| pUDI135 | ScPDC1p-HK3-ScPDC1t | This study |
| pUDI136 | ScPDC1p-HK4-ScPDC1t | This study |
| pUDI206 | ScHXK2-HK2-tHXK2 | This study |

| | | |
|---|---|---|
| pUDI207 | ScHXK2-HK4-tHXK2 | This study |
| pUDI137 | ScTEF2p-GPI1-ScTEF2t | This study |
| pUDI138 | ScTEF1p-PFKM-ScTEF1t | This study |
| pUDI139 | ScTEF1p-PFKP-ScTEF1t | This study |
| pUDI140 | ScTEF1p-PFKL-ScTEF1t | This study |
| pUDI141 | ScFBA1p-ALDOA-ScFBA1t | This study |
| pUDI142 | ScFBA1p-ALDOB-ScFBA1t | This study |
| pUDI143 | ScFBA1p-ALDOC-ScFBA1t | This study |
| pUDI144 | ScTPI1p-TPI-ScTPI1t | This study |
| pUDI145 | ScTDH3p-GAPDH-ScTDH3t | This study |
| pUDI146 | ScTDH3p-GAPDHS-ScTDH3t | This study |
| pUDI147 | ScPGK1p-PGK1-ScPGK1t | This study |
| pUDI148 | ScPGK1p-PGK2-ScPGK1t | This study |
| pUDI149 | ScGPM1p-PGAM1-ScGPM1t | This study |
| pUDI150 | ScGPM1p-PGAM2-ScGPM1t | This study |
| pUDI151 | ScENO2p-ENO1-ScENO2t | This study |
| pUDI152 | ScENO2p-ENO2-ScENO2t | This study |
| pUDI153 | ScENO2p-ENO3-ScENO2t | This study |
| pUDI154 | ScPYK1p-PKM1-ScPYK1t | This study |
| pUDI155 | ScPYK1p-PKM2-ScPYK1t | This study |
| pUDI156 | ScPYK1p-PKR-ScPYK1t | This study |

| | | |
|---|---|---|
| pUDI157 | ScPYK1p-PKL-ScPYK1t | This study |

Table S9E: Plasmids used as PCR template in this study

| Name | Relevant construct | Source |
|---|---|---|
| Plasmids used for construction of IMX165 | | |
| pSH47 | PGAL1-Cre-TCYC1, KlURA3 | [15] |
| pUG73 | loxP-*KlLEU2*-loxP cassette | [15] |
| pUG6 | loxP-*KanMX*-loxP cassette | [15] |
| Plasmids used for construction of IMX1076 | | |
| pUG-natNT1 | NatNT1 | [15] |
| p414-TEF1p-Cas9-CYC1t | TEF1p-Cas9-CYC1t | [16] |
| Multicopy plasmid used as PCR template | | |
| pUDE750 | ScHXK2p-HK4-ScHXK2t | This study |

Table S9F: gRNA plasmids for CRISPR Cas9 genome editing

| Name | Construct | Source |
|---|---|---|
| pMEL13 | 2µm ampR KanMX gRNA-CAN1.Y | [17] |
| pROS13 | 2µm ampR KanMX gRNA-CAN1.Y gRNA-ADE2.Y | [17] |

| | | |
|---|---|---|
| pMEL10 | 2µm ampR KlURA3 gRNA-CAN1.Y | [17] |
| pUDE342 | URA3 SNR52p-gRNA.SGA1-SUP4t RECYCLE SinLoG | [9] |
| pUDE327 | URA3 SNR52p-gRNA.*HXK2*-SUP4t | [9] |
| pUDR265 | 2µm ampR KanMX gRNA-PFK1 gRNA-PFK2 | This study |
| pUDR337 | 2µm | This study |
| pUDR338 | 2µm ampR KanMX gRNA-FBA1 gRNA-FBA1 | This study |
| pUDR371 | 2µm | This study |
| pUDR387 | 2µm | This study |
| pUDR376 | 2µm ampR AmdS gRNA-X2 | [18] |
| pUDR666 | 2µm | This study |
| pUDR667 | 2µm | This study |

### Appendix 1: Computational modelling of human hexokinases complementation

The human hexokinases *Hs*HK1 and *Hs*HK2 can only complement the yeast hexokinase upon mutations in the vicinity of the catalytic pocket. Therefore we aimed at investigating (1) why the native human hexokinases cannot complement the yeast hexokinase and (2) how mutations of the human hexokinases allow growth. In order to tackle these questions we have taken a computational modelling approach starting from the yeast glycolysis model from van Heerden *et al*. [19] which is based on the model by Teusink *et al*. [20] Since growth cannot be directly addressed based exclusively on a metabolic model such as the glycolysis model, our approach was to evaluate changes in glycolytic fluxes as an index with a direct causal relation to the growth rate of cells. Starting from the van Heerden model [19], a step-by-step procedure was followed, resulting in the 5 models described in the flowchart 1 below.

Flowchart 1: Overview of models created in Copasi.

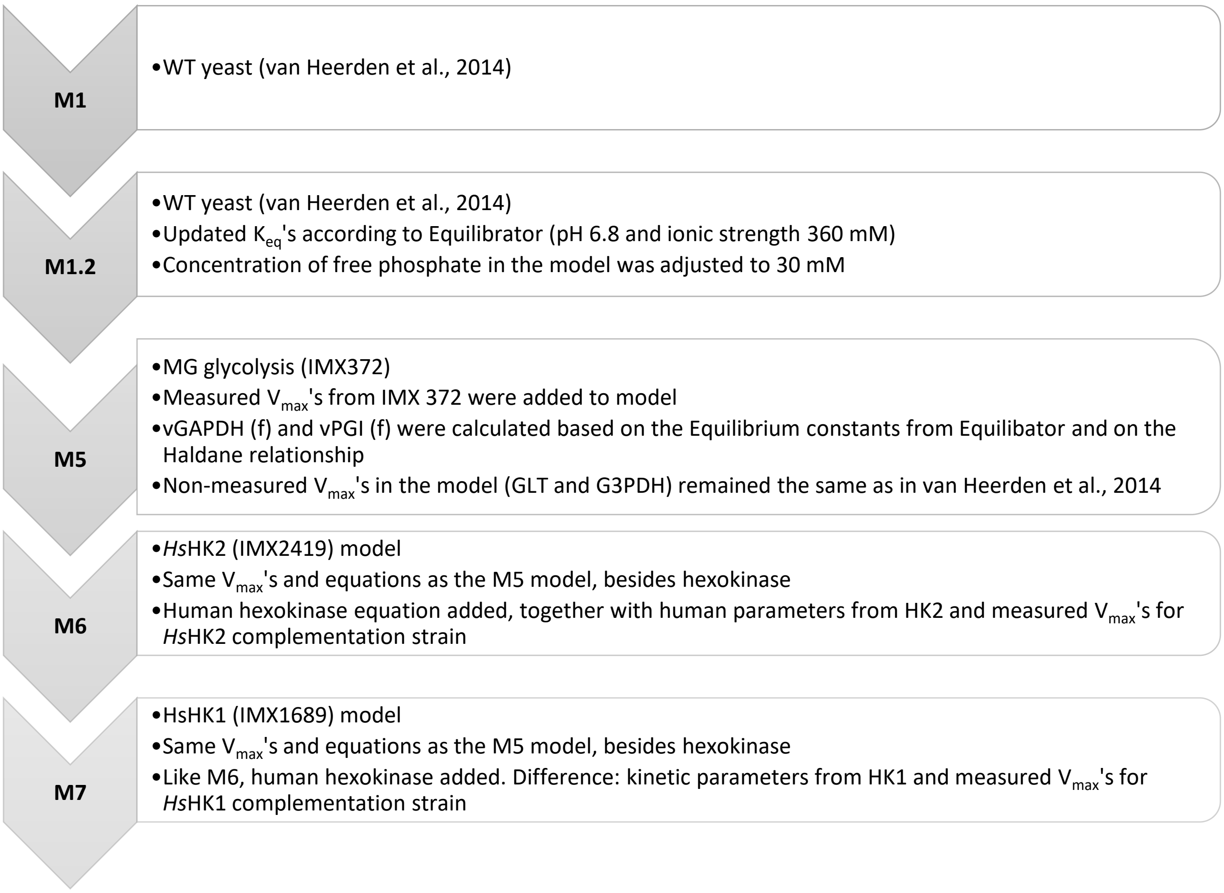


First, the Equilibrium constants of the model were updated based on values computed with Equilibrator (http://equilibrator.weizmann.ac.il/) at pH 6.8 and ionic strength 360 mM [21]. The list of Equilibrium constants used can be found below in Table A1. These values are necessary to calculate the reverse $V_{max}$ when only the forward $V_{max}$ was measured or vice-versa. To make the model more robust to changes and still physiologically relevant, the steady-state concentration of intracellular free phosphate was set as 30 mM. This value is within the range of 10-75 mM reported by van Eunen *et al.* (2010) [21], and consistent with the fact that the growth medium contained 22 mM phosphate. These proposed changes were necessary for the MG model (M5) to reach a stable steady-state.

Table A1: Equilibrium constants ($K_{eq}$) for the forward reactions (glycolysis as reference) catalysed by the enzymes in the first column

| Enzyme | Equilibrator* |
|---|---|
| HK | 548 |
| PGI | 0.36 |
| FBA | 0.00014 |
| TPI | 0.11 |
| GAPDH | 0.103 |
| PGK | 1,700 |
| PGAM | 0.184 |
| ENO | 5.190 |
| PYK | 88,000 |
| ADH (reverse reaction) | 0.00016 |
| AK | 0.471 |
| G3PDH | 43,000 |

*No $K_{eq}$ used for PFK and PDC in the van Heerden model, as these reactions were treated as irreversible.

From Teusink *et al.* [20], we have the following relationship for volume and protein content in yeast: 3.75 µL cell vol/mg protein ( = $3.75 \cdot 10^{-6}$ L/mg protein). The measured $V_{max}$'s for the IMX372 strain (Minimal Glycolysis, MG) are shown below both in µmol/(min · mg prot) (Table A2) as well as the mM/min equivalents (Table A3) calculated based on the intracellular volume.

Table A2: $V_{max}$'s in µmol/(min · mg prot)

| Enzyme | MG strain (IMX372) | HK2 compl. strain (IMX2419)[#] | HK1 compl. strain (IMX1689)[#] |
|---|---|---|---|
| HK | 1.32 | 0.567 | 0.526 |
| PGI (rev) | 3.37 | 3.37 | 3.37 |
| PFK | 0.72 | 0.72 | 0.72 |
| FBA | 1.60 | 1.60 | 1.60 |
| GAPDH (rev) | 4.26 | 4.26 | 4.26 |
| PGK (rev) | 9.46 | 9.46 | 9.46 |
| GPM | 9.66 | 9.66 | 9.66 |
| ENO | 1.85 | 1.85 | 1.85 |
| PYK | 8.90 | 8.90 | 8.90 |
| ADH | 1.37 | 1.37 | 1.37 |
| PDC | 1.49 | 1.49 | 1.49 |

[#]Assumed to be the same as the MG strain IMX372, except for hexokinase in the complementation strains.

Table A3: $V_{max}$'s in mM/min

| Enzyme | MG strain (IMX372) | HK2 compl. strain (IMX2419) | HK1 compl. strain (IMX1689) |
|---|---|---|---|
| HK | 351 | 151 | 140 |
| PGI (rev) | 897 | 897 | 897 |
| PFK | 191 | 191 | 191 |
| FBA | 428 | 428 | 428 |
| GAPDH (rev) | 1135 | 1135 | 1135 |
| PGK (rev) | 2522 | 2522 | 2522 |
| GPM | 2576 | 2576 | 2576 |
| ENO | 493 | 493 | 493 |
| PYK | 2372 | 2372 | 2372 |
| ADH | 365 | 365 | 365 |
| PDC | 398 | 398 | 398 |

For PGI and GAPDH the forward $V_{max}$'s were calculated from the measured reverse $V_{max}$'s based on the Haldane relation. Table 4 summarizes the $V_{max}$'s used for the simulations followed by the calculations performed according to the Haldane relationship.

Table A4: $V_{max}$'s in mM/min used as input for the simulations

| Enzyme | MG strain (IMX372) | HK2 compl. strain (IMX2419)# | HK1 compl. strain (IMX1689) # |
|---|---|---|---|
| HK | 351 | 151 | 140 |
| PGI (f) | 1507 | 1507 | 1507 |
| PFK | 191 | 191 | 191 |
| FBA | 428 | 428 | 428 |
| GAPDH (f) | 3758 | 3758 | 3758 |
| GAPDH (rev) | 1135 | 1135 | 1135 |
| PGK (rev) | 2522 | 2522 | 2522 |
| GPM | 2576 | 2576 | 2576 |
| ENO | 493 | 493 | 493 |
| PYK | 2372 | 2372 | 2372 |
| ADH | 365 | 365 | 365 |
| PDC | 398 | 398 | 398 |

For GAPDH, the following Haldane relationship applies:

$$K_{eq} = \frac{V_f\ K_{mBPG} K_{mNADH}}{V_r\ K_{mNAD}\ K_{mGAP}\ K_{mPi}}$$

a. $K_mBPG$ = 0.0098 mM, $K_mNADH$ = 0.06 mM, $K_mNAD$=0.09 mM, $K_mGAP$ = 0.21 mM, $K_mPi$ = 1 mM, $K_{eq}$ = 0.103 $mM^{-1}$
b. $V_r$ = 1135 mM/min
c. ∴ $V_f$ = 3758 mM/min

For PGI, the following Haldane relationship applies:

$$K_{eq} = \frac{V_f \, K_{mF6P}}{V_r \, K_{mG6P}}$$

a. $K_mF6P$ = 0.3 mM, $K_mG6P$ = 1.4 mM, $K_{eq}$ = 0.36
b. $V_r$ = 897 mM/min
c. ∴ $V_f$ = 1507 mM/min

A comparison between the original van Heerden model (M1) and the model with updated $K_{eq}$'s and free phosphate concentrations (M1.2) shows no effect on steady-state fluxes and small effects on steady-state metabolite concentrations, with the exception of TRIO (the sum of GAP and DHAP) and NADH (Fig. A1). This can be explained by a 18-fold difference in the $K_{eq}$ for GAPDH used for model M1.2 when compared to M1.

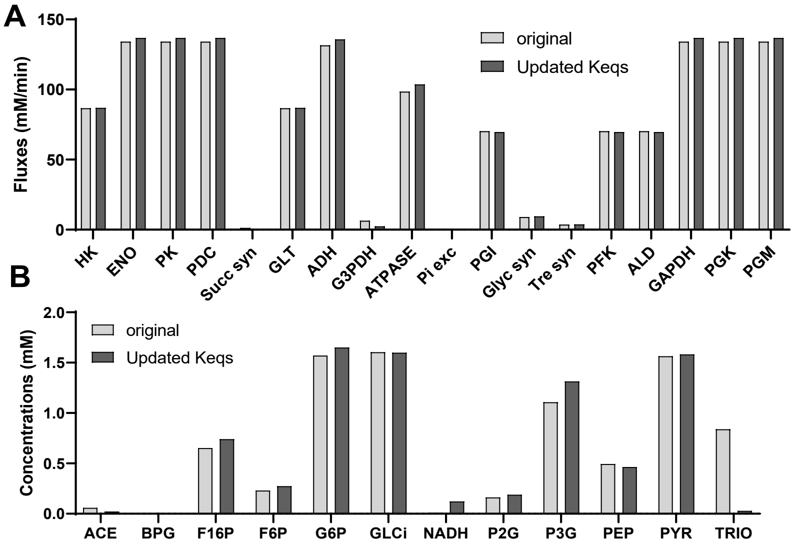


Figure A1: Comparison of steady-state fluxes (A) and concentrations (B) between the van Heerden model and the model with updated $K_{eq}$'s and phosphate concentrations.

The equation for yeast hexokinase (below) followed a generic bi-substrate equation with random binding order. A similar equation was used for the human hexokinase, with G6P binding competitive with ATP and ADP and parameters (Table A5) as described below.

$$v_{YeastHK} = \frac{\frac{V_{max}}{K_{m,Glc_i} \cdot K_{m,ATP}} \cdot \left( [Glc_i] \cdot [ATP] - \frac{[G6P] \cdot [ADP]}{K_{eq}} \right)}{\left( 1 + \frac{[Glc_i]}{K_{m,Glc_i}} + \frac{[G6P]}{K_{m,G6P}} + \frac{[G6P]}{K_{i,G6P}} \right) \cdot \left( 1 + \frac{[ATP]}{K_{m,ATP}} + \frac{[ADP]}{K_{m,ADP}} \right)}$$

$$v_{HumanHK1/HK2} = \frac{\frac{V_{max}}{K_{m,Glc_i} \cdot K_{m,ATP}} \cdot \left( [Glc_i] \cdot [ATP] - \frac{[G6P] \cdot [ADP]}{K_{eq}} \right)}{\left( 1 + \frac{[Glc_i]}{K_{m,Glc_i}} + \frac{[G6P]}{K_{m,G6P}} + \frac{[ADP]}{K_{i,ADP}} \right) \cdot \left( 1 + \frac{[ATP]}{K_{m,ATP}} + \frac{[ADP]}{K_{m,ADP}} + \frac{[G6P]}{K_{i,G6P}} \right)}$$

Equation yeast hexokinase:

- $K_i$G6P: proxy for trehalose 6-P (T6P) inhibition, competitive for glucose [19]

Equation human hexokinase:

- $K_i$G6P: glucose 6-P inhibition, competitive for ATP [22]
- $K_i$ADP: ADP inhibition. This inhibition is described to be mixed, but for simplicity only the competitive term was included in the rate equation [22]

Table A5: Differences between yeast and hexokinase kinetic parameters.

| Parameter | *Sc*HXK [19] | *Hs*HK2 [22] | *Hs*HK1 [22] |
|---|---|---|---|
| $K_m$ ATP (mM) | 0.15 | 0.78 | 0.4 |
| $K_m$ Glucose (mM) | 0.08 | 0.23 | 0.045 |
| $K_m$ ADP (mM) | 0.23 | 2.2 | 0.62 |
| $K_m$ G6P (mM) | 30 | 0.16 | 0.21 |
| $K_i$ ADP (mM) | - | 5.4 | 2.0 |
| $K_i$ G6P (mM) | 0.07 ($K_i$ T6P) | 0.021 (native)[#]<br>0.063 (mutated)* | 0.026 (native)[#]<br>0.21 (mutated)* |

[#]Literature reference values

*Estimated from the $IC_{50}$ measurements from strain IMX1690 for HsHK2 and from strain IMS1137 for HsHK1.

Figure A2 shows a comparison between the four generated models (M1.2, M5, M6 and M7). Fluxes, metabolite concentrations and the control of glucose uptake and ethanol fluxes are depicted. When either human hexokinase 2 or 1 were added to the model (M6 and M7), steady-state glycolytic fluxes (quantified as the flux through the glucose transporter GLT, and alcohol dehydrogenase ADH) were around one third of those in the WT and MG models (M1.2 and M5) (Fig. A2-A). In the human hexokinase models (M6 and M7), predicted ATP concentrations at steady state were around 50% lower than in the WT and MG controls (Fig. A2-B). Panel C of Fig. A2 shows that the human hexokinase models differ from the WT and MG in terms of the distribution of glycolytic intermediates. An apparent bottleneck at hexokinase in the human hexokinase models (M6 and M7 can be recognized from the accumulation of more intracellular glucose ($Glc_i$) than in the control models. This is confirmed by a higher flux control coefficient of the human hexokinases on the glycolytic fluxes in models M6 and M7 when compared to the control models (Panels D and E of Fig. A2). This is accompanied by higher flux control at the PFK and GAPDH nodes, which is compensated by lower flux control by glucose uptake (GLT) and ATP utilization (ATPase).

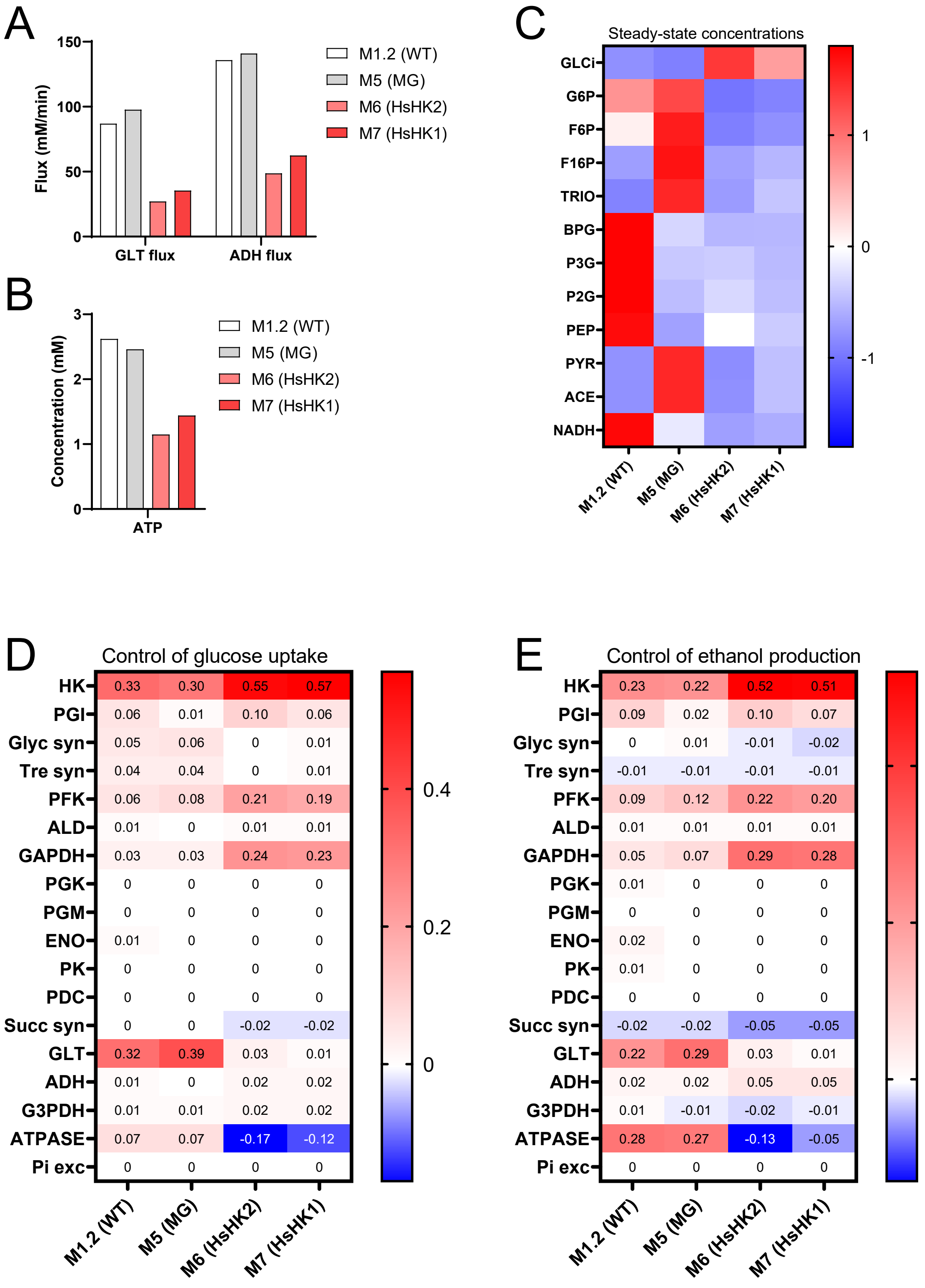


Figure A2. A: Predicted steady-state fluxes. B: ATP concentrations at steady state. C: heatmap for glycolytic intermediates in each model (z-score normalized per metabolite). D-E: Flux control coefficients for vGLT (glucose uptake) and vADH (ethanol production), respectively.

To evaluate whether trehalose-6P (T6P) would inhibit *Hs*HK2, the activity of the enzyme was measured in the presence of 1 mM  T6P. This inhibited both *Hs*HK2 (native) and *Hs*HK2$^{L776F}$ (mutated) by roughly 50% (Fig. S4B). We used this information to evaluate whether T6P inhibition would have a further impact on the glycolytic fluxes of the *Hs*HK2 complementation strain with the use of model M6. The $IC_{50}$T6P for human hexokinase 2 can be estimated to be 1 mM at the experimental conditions used. Assuming competitive inhibition, we can estimate the range of $K_i$T6P for the human enzyme given the following equation [23]:

$$IC_{50} = K_i\left(1 + \frac{[S]}{K_m}\right)$$

Given that the concentration of glucose used in the assay is 1 mM and its $K_m$ is 0.23 mM, the term 1+ $[S]/K_m$ can be approximated to 5, which means that the $K_i$ would be roughly $IC_{50}/5$, or 0.2 mM. This is 3 times higher than the Ki for T6P reported for the yeast hexokinase in the van Heerden model (0.07 mM). Including this parameter in the human hexokinase equation (representing a competitive term in the glucose binding pocket) results in no changes in glycolytic fluxes. A parameter scan performed with the *Hs*HK2 model (M6) shows little to no effect of the parameter within this range (Fig A3).

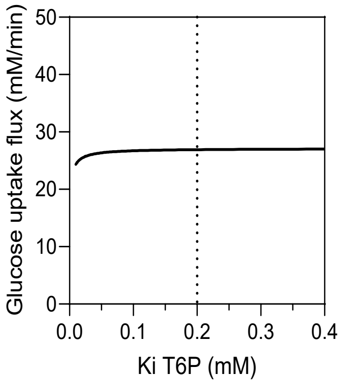


Figure A3: $K_i$T6P scan for HsHK2 model (M6). Dashed line: estimated $K_i$T6P for HsHK2.
